# Chronic inflammation drives epididymal tertiary lymphoid structure formation and autoimmune fertility disorders

**DOI:** 10.1101/2024.11.12.623224

**Authors:** Maia L. Elizagaray, Ferran Barrachina, Maria C. Avenatti, Isinsu Bastepe, Angela Chen, Ainize Odriozola, Oluchi Ukairo, Vanina Da Ros, Kiera Ottino, Nerea Subiran, Maria A. Battistone

**Author notes:** Correspondence: Maria Agustina Battistone, PhD, MGH CNY, 149 13th St. Suite 4.325H, Boston, MA 02129. equal contribution (Ferran Barrachina, Maria C. Avenatti).

## Abstract

The incomplete understanding of epididymal mucosal immunity is a significant contributing factor to the classification of many male infertility cases as idiopathic. Conditions that disrupt the immune balance in the male reproductive tract, such as vasectomy and infections, can expose sperm to the immune system, leading to increased production of anti-sperm antibodies (ASAs) and subsequent reproductive challenges. Regulatory T cells (Tregs) regulate inflammation and maintain sperm tolerance. In a murine model, we demonstrated that disrupting sperm immunotolerance induces chronic autoimmune responses characterized by antibody production targeting sperm and reproductive tissue autoantigens and unique tissue-specific immune cell signatures in the epididymis and testis. Such inflammatory features impair sperm function, contribute to epididymal damage, and drive sustained male subfertility. Tertiary lymphoid structures (TLSs) were formed within the epididymis after Treg depletion, defined by clusters of heterogenous B and T cells, fibroblasts, and endothelial cells. These ectopic structures perpetuate inflammation and lower the activation threshold for future immune threats. Similar isotypes of autoantibodies were detected in the seminal plasma of infertile patients, suggesting shared mechanistic pathways between mice and humans. Overall, we provide an in-depth understanding of the diverse B- and T-cell dynamics and TLS formation during epididymitis to develop precision-targeted therapies for infertility and chronic inflammation. Additionally, this immunological characterization of the epididymal microenvironment has the potential to identify novel targets for the development of male contraceptives.

**One Sentence Summary:** Understanding the epididymal immune cell landscape dynamics aids in developing targeted therapies for infertility and contraception.

## Introduction

Chronic inflammation and autoimmune conditions in the male reproductive system lead to impaired sperm production, function, and transport, thereby contributing significantly to male infertility. Male infertility accounts for roughly 50% of the 17% of infertility cases worldwide (*1*). Immunological factors are believed to contribute to approximately 15% of male infertility cases, though this may be an underestimation, as immunological aspects in infertile patients remain underexplored (*2*). The incomplete understanding of epididymal immunity is a major reason why many cases of male infertility remain classified as “idiopathic.” Immunological infertility is defined by the presence of elevated anti-sperm antibody (ASA) levels that interfere with reproductive function (*3, 4*). Vasectomy, congenital absence of vas deferens, infections, or trauma can result in the exposure of sperm to the immune system, thereby increasing the production of ASAs (*3, 4*). However, the immune mechanisms underlying the presence of ASAs remain uncertain (*1*).

Regulatory T cells (Tregs) control inflammation and prevent immune responses against sperm in the testis, where spermatozoa are produced, and the epididymis, where they mature and are stored (*5–7*). We previously reported that severe autoimmune orchitis and epididymitis developed 2 weeks after Treg depletion in a murine model (*5, 6*). This breakdown in immunotolerance affects sperm quantity and quality and increases pro-inflammatory heterogeneous immune cell infiltration, ultimately leading to severe reproductive impairment (*5*).

The recognition of sperm antigens as foreign by immune cells is attributed to their delayed establishment, occurring long after the self-tolerance setup process, during puberty. When protective epithelial barriers are compromised, the typically compartmentalized autoantigens are exposed to the immune system, potentially inducing autoreactive B-lymphocytes to produce autoantibodies (*6–8*). Autoreactive B cell proliferation is a defining characteristic of autoimmune disorders and a process largely driven by affinity maturation against autoantigens within B cell follicles, typically in secondary lymphoid organs. However, B cells in follicles and germinal centers (GCs) can also mature in ectopic locations, defined as tertiary lymphoid structures (TLSs), which form in response to chronic inflammation, autoimmune diseases, localized infections, and solid tumors (*9, 10*). TLSs mimic secondary lymphoid organs by featuring distinct T and B cell zones, follicular dendritic cell (DC) networks, high endothelial venules, and specialized immunofibroblasts. TLSs are fitted to support localized adaptive immune responses (*11*). For B cell activation and the effectiveness of antibody production, B cells must capture, process, and present antigens while receiving signals from other cells, such as mononuclear phagocytes (MPs), epithelial cells, and T lymphocytes.

Heterogeneity is a distinguishing feature of the different segments of the epididymis, highlighting the complexity of their segment-specific immune responses (*12–14*). In this study, we conducted a region-specific analysis of the local and systemic immune landscape in autoimmune-induced epididymitis. Our investigation focused on characterizing the distinct humoral, T- and B-cell-mediated immune responses which are crucial for maintaining sperm immunotolerance. We uncovered that immune tolerance disruption triggered an early and persistent pro-inflammatory response, adversely impacting the epididymis and culminating in sustained male subfertility. This condition might be partially maintained by the TLS formation within the epididymis, a feature that had never been reported in this organ. Similar autoantibody isotypes were detected in the seminal plasma of infertile individuals, suggesting shared mechanistic pathways underlying in both mice and humans. Deciphering these immunoregulatory mechanisms expands our understanding of the largely unexplored post-testicular environment and its critical role in perpetuating infertility. These insights could lead to innovative therapeutic approaches for male infertility and help identify new targets for male immuno-contraception.

## Results

### Treg depletion induced early epididymal epithelial changes

Using transgenic mice that express the diphtheria toxin (DT) receptor (DTR) under the control of the Foxp3 promoter (Foxp3-DTR/EGFP mice), we disrupted immunotolerance by ablation of Tregs using a DT-based depletion protocol (*5, 6*) (Fig. 1Ai). The total Treg cell number decreased in the proximal (initial segments (IS), and caput) and distal (corpus and cauda) epididymis and the testis 1 day after DT injection (Fig. 1A-B). The Treg depletion within the epididymis was further confirmed by confocal microscopy (Fig. 1C). To discriminate blood-borne circulating Tregs from tissue-resident cells, *in vivo* intravascular CD45^+^ cell staining was performed (*15*) (Fig. 1D*i*). The absolute number of resident (PE^-^) Tregs within the proximal and distal epididymis (Fig. 1D*ii-iii*) and testis (Fig. 1D*iv*) were significantly lower in the DT-treated Foxp3-DTR compared to WT. On the other hand, the local-circulating (PE^+^) Tregs in the epididymis and the testis showed no significant changes following DT injection (Suppl. Fig. 1).

**Figure 1.**
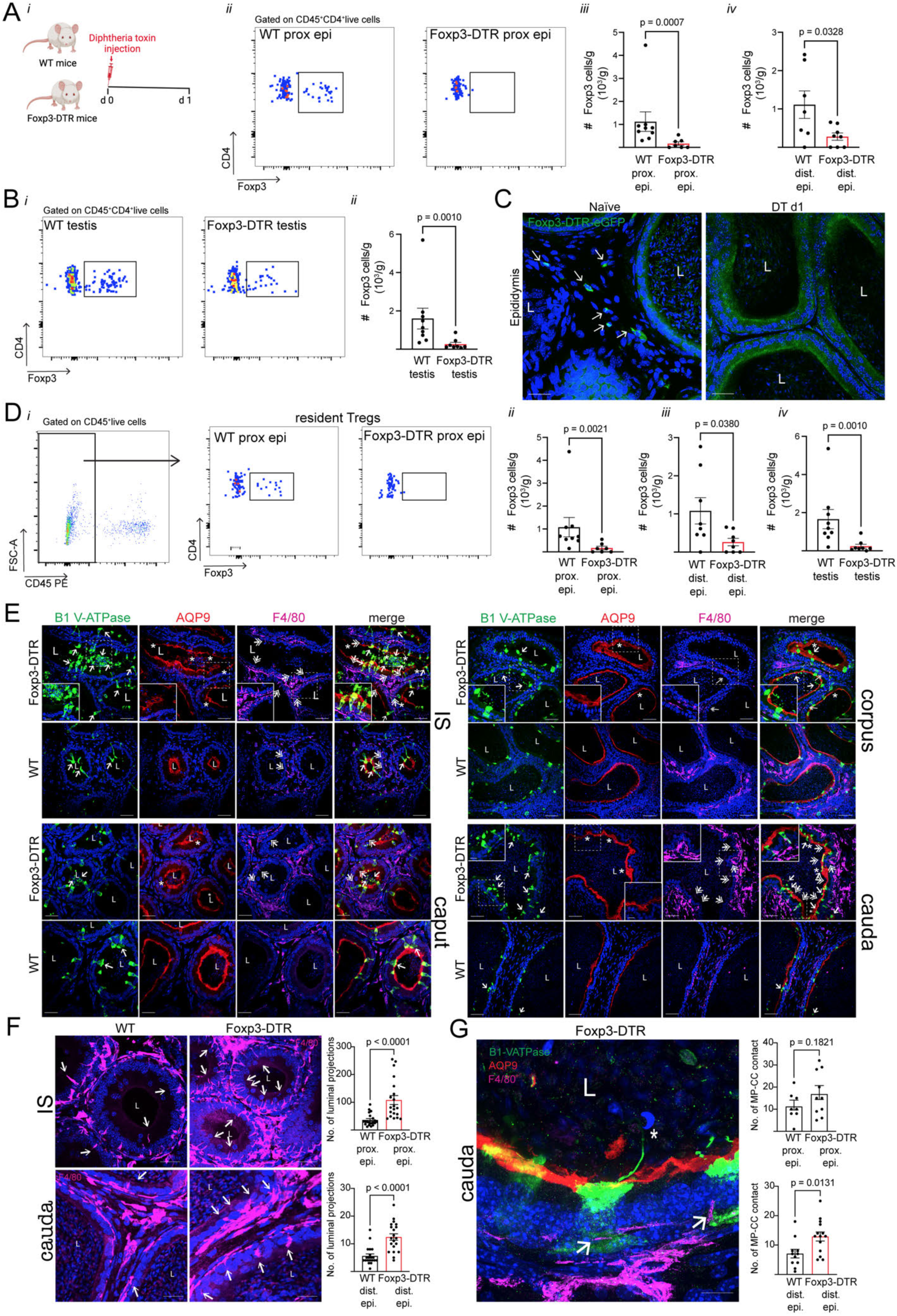
Regulatory T cell (Treg) depletion in the epididymis and testis and epididymal epithelial damage of Foxp3-DTR mice 1 day after diphtheria toxin (DT) treatment. **A)** Diagram of Treg depletion protocol and subsequent analysis 1 day after DT treatment (*i*). Flow cytometry gating strategy for identifying Tregs (Foxp3^+^CD4^+^CD45^+^) in the proximal and distal epididymides (*ii).* Absolute count (#) of total Tregs per gram of proximal and distal epididymis (*iii-iv)*. **B)** Flow cytometry gating strategy for identifying Tregs in the testis (*i)*. Absolute count of total Tregs per gram of testis (*ii)*. **C)** Confocal microscopy images showing Foxp3-EGFP^+^ Tregs (green, arrows) in the epididymis of naïve and 1-day DT-injected Foxp3-DTR transgenic mice. **D)** Flow cytometry gating strategy for identifying tissue-resident Tregs (CD45-PE^-^ Foxp3^+^CD4^+^CD45-BV711^+^) (*i*). Absolute count (#) of tissue-resident Tregs per gram of proximal and distal epididymis (*ii-iii*) and the testes (*iv*). Each dot represents a pool of 2 proximal epididymides, 2 distal epididymides, or 2 testes from each mouse. **E)** Confocal microscopy images showing B1 V-ATPase^+^ cells (clear cells (CCs); green), AQP9^+^ cells (principal cells; red), and F4/80^+^ mononuclear phagocytes (MPs, magenta) in all epididymal regions of WT and Foxp3-DTR mice 1 day after DT injection. Proximal epididymis comprises initial segments (IS) and caput; Distal epididymis comprises corpus and cauda. Arrows show apical cellular protrusions (blebs) from B1 V-ATPase^+^ cells (green; CCs) in different regions of the epididymis from DT-injected WT (control) and Foxp3-DTR mice. Asterisks show AQP9 morphological alteration in epididymal epithelia. Double-head arrows indicate F4/80^+^ luminal-reaching projections. **F)** Imaging and quantification of the number of F4/80^+^ projections (arrows) in the proximal (IS) and distal (cauda) epididymis of DT-injected WT and Foxp3-DTR mice per area of tissue (110,540 µm^2^). Each image quantification is represented as a dot. **G)** Imaging of B1 V-ATPase^+^ CC (green) and F4/80^+^ MP (magenta) contacts (arrows) in the cauda region. Notice the B1 V-ATPase^+^ nanotube reaching a sperm cell in the lumen (asterisk). Quantification of the number of CC-MP contacts in the proximal (IS and caput) and distal (cauda) (arrows) of DT-injected WT and Foxp3-DTR mice per area of tissue (110,540 µm^2^). Each image quantification is represented as a dot. Nuclei are labeled with DAPI (blue). Bars: 20 μm. L: Lumen. Data were analyzed using Student’s t-test (A*iv*, D*iii,* and G) or Mann-Whitney test (A*iii*, B*ii,* D*ii*, D*iv,* and F). Data are shown as means ± SEM.

Hematoxylin and eosin (H&E) staining revealed a higher incidence of damaged sites in both the proximal and distal regions of the Treg-depleted epididymis versus controls (Suppl. Fig. 2A-B). The Treg-depleted epididymal epithelium presented injury features versus the control: clear cells (CCs, B1 V-ATPase^+^ cells) changed their morphology, increased their membrane protrusions and shed several of them into the lumen (Fig. 1E, green (arrows), inset in the green panel); and principal cells (PCs, aquaporin 9^+^ (AQP9) cells) showed epithelial discontinuation (Fig. 1E, red (asterisks), inset in the red panel) and thicker microvilli. We observed that Treg-depleted cauda presented multiple F4/80^+^ MP luminal-reaching projections like the ones usually detected in the IS (Fig. 1E, magenta (double arrowheads), inset in the magenta panel). Quantification of these projections was statistically higher in the Treg-depleted epididymis (Fig. 1F). MP luminal-reaching projections were observed in close interaction with the CCs (Fig. 1E and 1G). The number of CC-MP contacts was amplified in the distal (Fig. 1G, lower panel) but not in the proximal (Fig. 1G, upper panel) epididymal regions, suggesting increased cell-to-cell crosstalk in this area (*13, 16*). Please note that CCs extend a nanotube (*13*) that touches the sperm head (positive for DAPI, Fig. 1G, asterisk). We didn’t detect testicular damage or spermatogenesis impairment at this early time point (Suppl. Fig. 2C).

### Systemic and local autoantibody production 2 weeks after DT treatment

Our previous study revealed that at 2 weeks post-Treg depletion, there is a notable antibody deposition in both the epididymis and testis, together with elevated total IgG autoantibody levels in the serum (*5*). In this study, we further investigated the molecular intricacies of these humoral responses post-Treg depletion by isotyping the autoantibodies by ELISA. This showed distinct humoral profiles based on the antigen tested. In Treg-depleted mouse serum, increases were observed in IgM, IgG2b, IgG2c, and IgA against testicular antigens (Suppl. Fig. 3A). IgM, IgG1, IgG2b, and IgG2c were elevated against epididymal antigens (Suppl. Fig. 3B). Notably, IgM and IgG2c levels rose against proximal sperm antigens (Suppl. Fig. 3C). At the same time, IgM, IgG1, IgG2b, IgG2c, IgG3, and IgA were higher against distal sperm antigens (Suppl. Fig. 3D).

Regarding the local presence of autoantibodies within tissues, ELISA results revealed that in testicular homogenates IgG1, IgG2b, IgG2c, and IgG3 levels were significantly increased against testicular antigens 2 weeks after Treg depletion (Fig. 2A). Additionally, epididymal homogenates displayed elevated levels of IgM, IgG1, IgG2b, IgG2c, IgG3, and IgA against epididymal antigens (Fig. 2B). Confocal microscopy showed the accumulation of the different isotypes in the testicular (Suppl. Fig 4, green) and epididymal (Fig. 2C-F, green; for other epididymal regions and isotypes, refer to Suppl. Fig. 5-9) interstitium of DT-injected Foxp3-DTR mice. Notably, in the epididymal lumen of Treg-depleted mice, we observed agglutinated sperm containing IgG1^+^ and IgG2c^+^ ASAs (green in Fig. 2G, indicated by arrowheads). Interestingly, some of these clusters of ASA-bound sperm (green) were surrounded by F4/80^+^ MPs (red) (asterisks in Fig. 2G), potentially indicating phagocytosis of antibody-bound sperm. Concomitantly, anti-sperm IgG2c (red) antibodies were attached to both the head and tail of sperm from Treg-depleted mice (Fig. 3A). Antibody-secreting B lymphocytes (CD19^+^) were localized in the interstitium of the epididymis (Fig. 2C-F, and Suppl Fig. 5-9). A conglomeration of different cell types (Fig. 2F, Supp. Fig. 7 and 8, dashed circles), in particular, B cells, was found in the epididymal interstitium.

**Figure 2.**
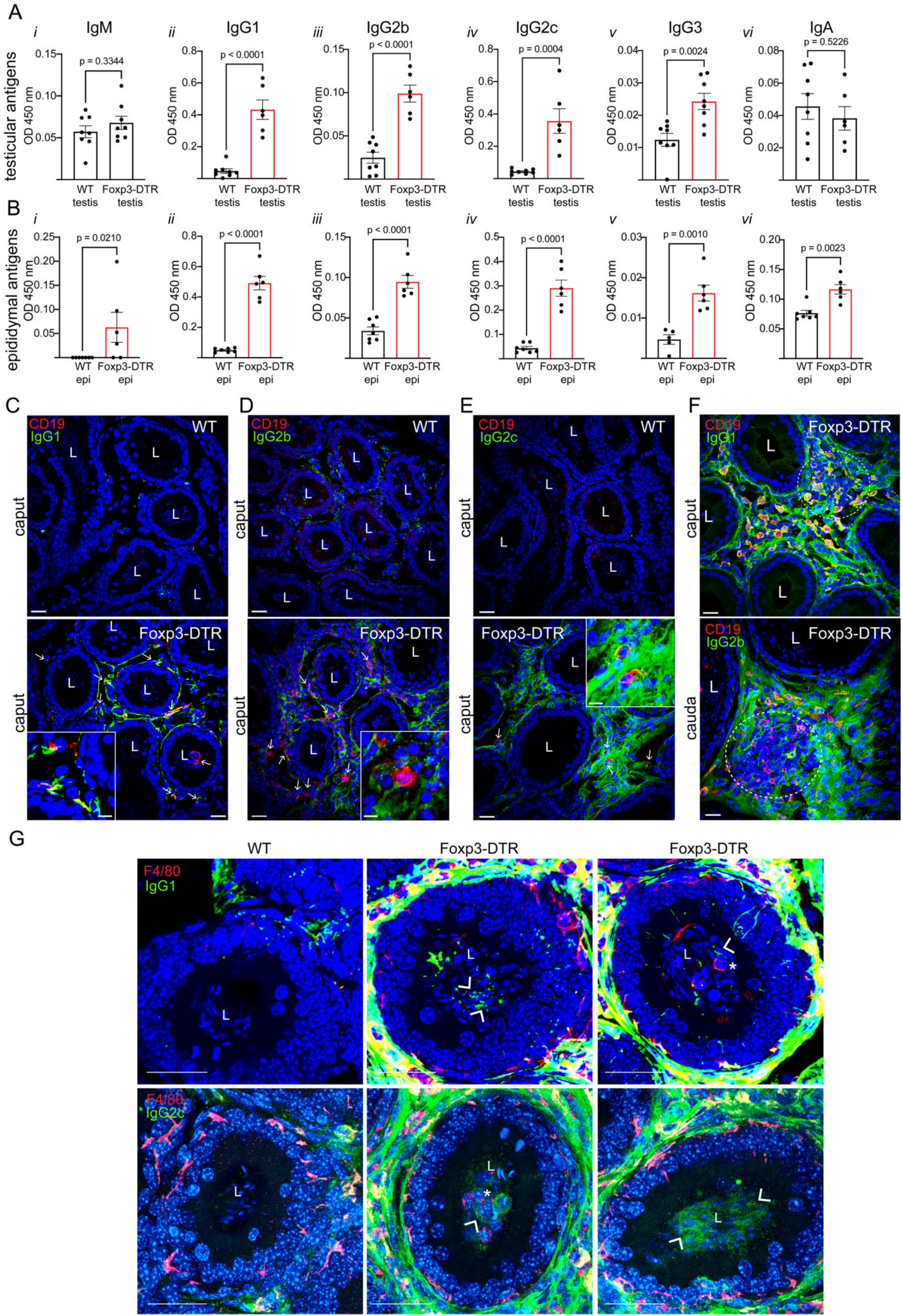
Isotypification of autoantibody levels in the testis and epididymis of Foxp3-DTR mice 2 weeks after Treg depletion. Isotypification of autoantibody levels (IgM, IgG1, IgG2b, IgG2c, IgG3, and IgA) in testicular homogenates against testicular antigens (**A**), and epididymal homogenates against epididymal antigens (**B**). Immunolabeling of CD19^+^ B cells (red, arrows) and IgG1 (green) **(C)**; IgG2b (green) **(D)** or IgG2c (green) **(E)**, in the caput region of DT-injected WT and Foxp3-DTR mice. **F)** Immunolabeling of CD19^+^ B cells (red) and IgG1 (green, upper panel) in the caput region and IgG2b (green, lower panel) in the cauda region of DT-injected WT and Foxp3-DTR mice. A dashed circle indicates aggregation of cells in the interstitium. Antibody deposition can be observed in the epididymal interstitium of the different regions. See images of testes and other epididymal regions in Supp. Fig. 4-9. **G)** Immunolabeling of F4/80^+^ MPs (red) and IgG1 (green, upper panels) and IgG2c (green, bottom panels) in the epididymis caput regions of DT-injected WT and Foxp3-DTR mice. Arrowheads show agglutination of sperm positive for IgG1 and IgG2c (green), surrounded by F4/80^+^ MPs (red, asterisks) in the DT-injected Foxp3-DTR mice. Nuclei are labeled with DAPI (blue). Bars: 20 μm. L: Lumen. Data were analyzed using Student’s t-test (A*i-vi*, B*ii-v*) or Mann-Whitney test (B*i* and *vi*). Data are shown as means ± SEM.

**Figure 3.**
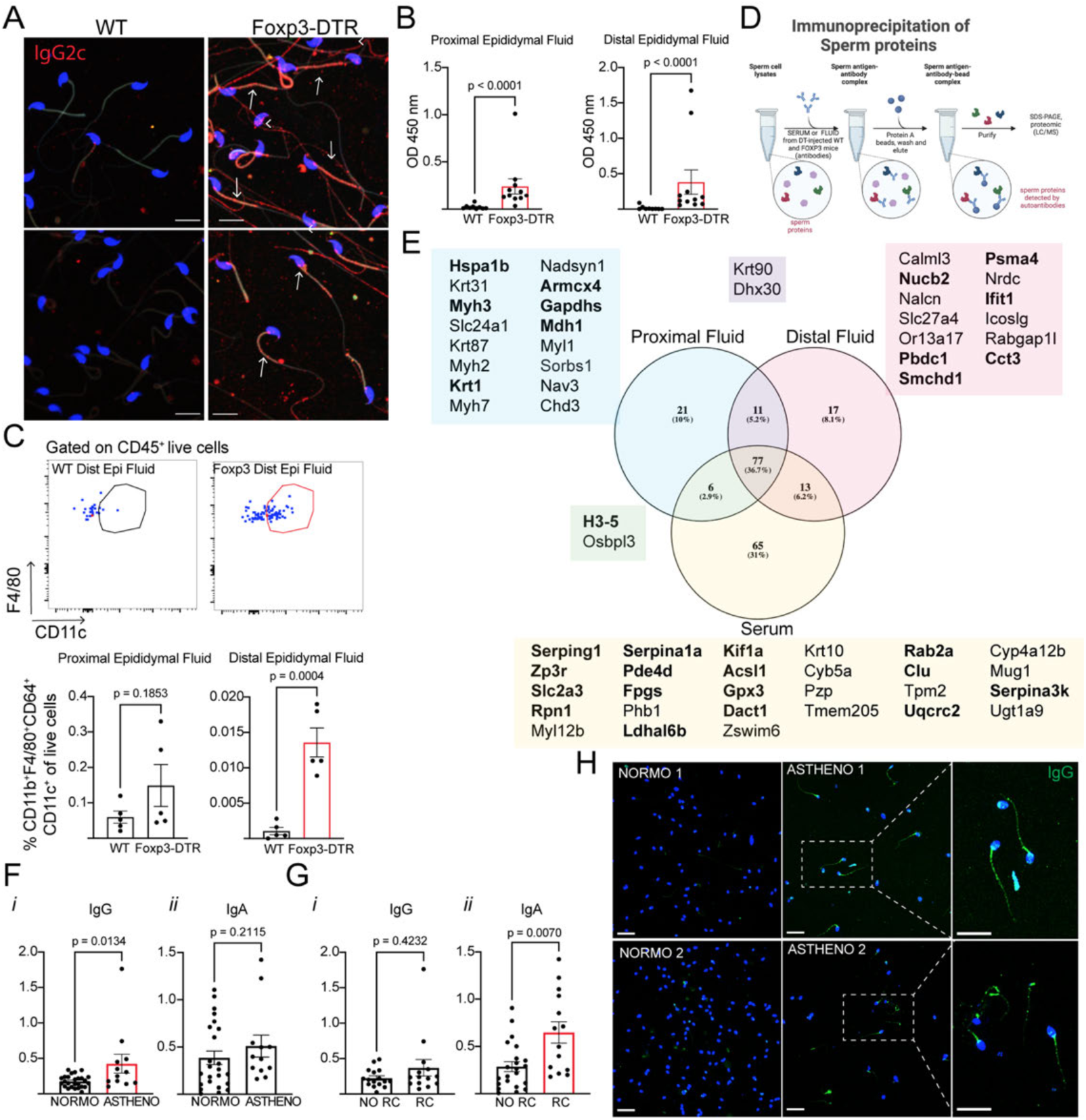
Anti-sperm antibodies (ASAs) in the epididymis of Foxp3-DTR mice 2 weeks after Treg depletion and in pathological human semen samples. **A)** Immunolabeling of IgG2c (red) on distal sperm from DT-injected WT and Foxp3-DTR mice. **B)** Total IgG autoantibodies against proximal and distal sperm antigens were detected in proximal and distal epididymal fluids of Treg-depleted mice by ELISA. **C)** Relative abundance of M1-like macrophages (CD11b^+^F4/80^+^CD64^+^CD11c^+^ of total live cells) in the proximal and distal epididymal fluids of DT-treated WT and Foxp3-DTR mice measured by flow cytometry. **D)** Sperm proteins immunoprecipitation with serum, proximal, and distal epididymal fluids protocol diagram. **E)** Venn diagram showing the serum/epididymal fluid-immunoprecipitated sperm proteins detected in the DT-treated Foxp3-DTR samples by proteomics. In bold are the proteins reported to be involved in sperm function. See also Supp. Tables I and II. **F)** Levels of total IgG (*i*) and IgA ASAs (*ii*) in seminal plasma from healthy (normozoospermic, normo) and pathological (asthenozoospermia, astheno) patients. **G)** Total IgG (i) and IgA ASA (ii) levels in seminal plasma of patients with low (no RC) versus high (RC) percentages of round cells. **H)** Total IgG (green) immunofluorescence imaging in the sperm fraction from normozoospermic and pathological (astheno) patients. Nuclei are labeled with DAPI (blue). Bars: 20 μm. Data were analyzed using Student’s t-test (C, G*ii*), or Mann-Whitney test (B, F, G*i*). Data are shown as means ± SEM.

### Sperm antigens detected by ASAs

Two weeks after Treg depletion, total IgG ASAs were higher in the proximal and distal epididymal fluids of the Treg-depleted mice compared to WT (Fig. 3B). Flow cytometry revealed an increase in the relative abundance of M1 pro-inflammatory macrophages (CD11b^+^F4/80^+^CD64^+^CD11c^+^ cells) in the distal epididymal fluid of the Foxp3-DTR group (Fig. 3C). Immunoprecipitation of sperm proteins using serum, as well as proximal and distal epididymal fluids from DT-treated WT and Foxp3-DTR mice, confirmed the presence of ASAs after Treg depletion (Fig. 3D-E and Suppl. Tables I and II). Seventy-seven proteins were identified across serum and epididymal fluids from both proximal and distal regions. Of these, 13 proteins were shared between serum and distal fluid, 11 between proximal and distal fluids, and 6 between serum and proximal fluid. In DT-treated Foxp3-DTR mice, 37 immunoglobulin (Ig)-related proteins were detected in serum. Meanwhile, 78 and 85 were found in proximal and distal fluids, respectively. Proteomics identified proteins involved in sperm maturation, motility, capacitation, and sperm-egg interaction, such as MDH1, GADPHS, KRT1, PSMA4, PDE4D, ZP3R, Rab2A, NUCB2, and CALML3 (*17–23*). Other key identified proteins are crucial for embryo development: H3-5 (*24*), SMCHD1 (*25, 26*), HSPA1B (*27*), and ARMCX4 (*28*). Some detected proteins, such as SORBS1 and TRM2, have been reported in the male reproductive tract but not specifically in spermatozoa; others, like PBDC1, are known to be present in sperm, but their role in fertility remains unclear. Additionally, we revealed novel sperm proteins, such as NALCN, which might interact with the CatSper channel (*29*), a key player in sperm motility and fertility (*30*). In addition, serum ASAs recognized complement-related proteins, some of which have already been described in sperm and semen (*31*). However, their direct impact on sperm function and fertility is not well-described.

Serum and epididymal fluid ASAs recognize proteins functioning as immune modulators, such as IFIT1, MUG1, ICOSLG, SERPINA1A, and CLU. IFIT1 has been detected in spermatozoa, with expression levels differing between semen samples that achieved pregnancy and those that did not (*32*). MUG1 regulates immune responses by facilitating the internalization of viral proteins (*33*); its role in spermatozoa is unknown. ICOSLG is essential for T-cell activation, differentiation, and cytokine production, enabling B-cell antibody secretion. Additionally, the interaction between ICOSLG and its receptor is crucial for TLS formation (*34, 35*). Although ICOSLG is expressed in the male reproductive system (*12, 13*), its role in fertility has not yet been described. Epididymal CCs, which are involved in immune responses induced by epididymitis, show increased *Icolg* expression in the distal region of the organ. They may also transfer this ligand to sperm during maturation (*12, 13*). SERPINA1A functions as a protease inhibitor, specifically targeting neutrophil elastase, and plays a critical role in regulating the enzymatic activities released by inflammatory cells. CLU is a crucial seminal plasma glycoprotein involved in sperm maturation, capacitation, and female immune tolerance (*36*). CLU has diverse immune functions, such as regulating complement activity. The fact that some of the sperm proteins detected by the ASAs are immune regulators may indicate that after injury, accessible sperm antigens may be triggering primordial signals of immune activation.

### Human ASAs detected in patients with fertility challenges

Given that Treg depletion in mice leads to an autoimmune response with ASA production, we next investigated whether similar ASAs are present in patients with compromised sperm function, particularly asthenozoospermic individuals facing fertility challenges. To characterize human ASA isotypes, we conducted ELISA on seminal plasma from 30 patients from assisted reproduction units at Cruces and Galdakao Hospital, Spain (Suppl. Table III). Our findings revealed elevated ASA levels in 20% of the samples. Asthenozoospermic patients exhibited significantly higher IgG ASA levels than normozoospermic patients (Fig. 3F*i*). However, no significant differences in IgA ASAs were observed between these groups (Fig. 3F*ii*). Conversely, samples with a high content of round cells (RC) showed a considerable increase in IgA ASAs (Fig. 3G*i-ii*). Additionally, we detected IgG ASAs on sperm heads and tails of patients with high ASA levels in their seminal plasma (Fig. 3H).

### B cell dynamics 2 weeks after Treg depletion

Two weeks after Treg ablation, B cells localized in the proximal (Fig. 4A and Suppl. Fig. 10A) and, to a lesser extent, in the distal epididymal interstitium (Suppl. Fig. 10A). t-SNE (t-distributed Stochastic Neighbor Embedding) and FlowSOM (clustering algorithm) analysis of the immune cell node (CD45^+^), identified 12 distinct cell subtypes and a shift in the immune cell landscape in the different regions of the epididymis (Fig. 4B*i-ii*) and the testis (Suppl. Fig. 11A) compared to the WT. At this time point and consistent with our previous findings (*5*), we observed an increase in the CD4^+^ T cells in both organs (Fig. 4B and Suppl. Fig. 11A). Additionally, the analysis highlighted a rise in different CD8^+^ T and CD20^+^ B cells across both tissues (Fig. 4B and Suppl. Fig. 11A). Remarkably, t-SNE and FlowSOM analysis of the B lymphocyte node (B220^+^, CD19^+^, CD20^+^) and B cell markers, identified 8 distinct B cell clusters following 2 weeks post-Treg-depletion in the proximal epididymis (Suppl. Fig. 10B*i*-*ii*) and the testis (Suppl. Fig. 11B*i*-*ii*), highlighting the diversity of B cell populations in these organs. Moreover, we observed higher counts of total and tissue-resident B cells in the testis (Suppl. Fig. 11C*i-iii*, D*i-iii*) and proximal epididymis (Fig. 4*Ci, iii, v, Di, iii, v*). Regarding the local-circulating cells, CD19^+^ and CD20^+^ B cells in the testis (Suppl. Fig. 11E*ii-iii*), B220^+^, CD19^+^, and CD20^+^ B cells in the proximal, and CD20^+^ B cells in the distal epididymis (Suppl. Fig. 10C*i-iii*) were elevated after Treg depletion. The relative abundance of total Plasma B cells (CD3⁻CD19⁺IgD⁻CD138⁺) raised in the proximal and distal regions and the testis (Fig. 4E*i-ii*, Supp. Fig. 11F), whereas local circulating cells were amplified in both epididymal regions (Supp. Fig. 10D*i-ii*). However, tissue-resident Plasma B cell expansion was observed only in the distal parts (Fig. 4F*ii*) and the testis (Suppl. Fig. 11G). These results suggest that autoantibodies are being produced *in situ* in these organs by the recruited B cells (note co-localization of CD19^+^ cells and IgG1, IgG2b, and IgG2c in the epididymis, Fig. 2C-G and Supp. Fig. 7A, 8A).

**Figure 4.**
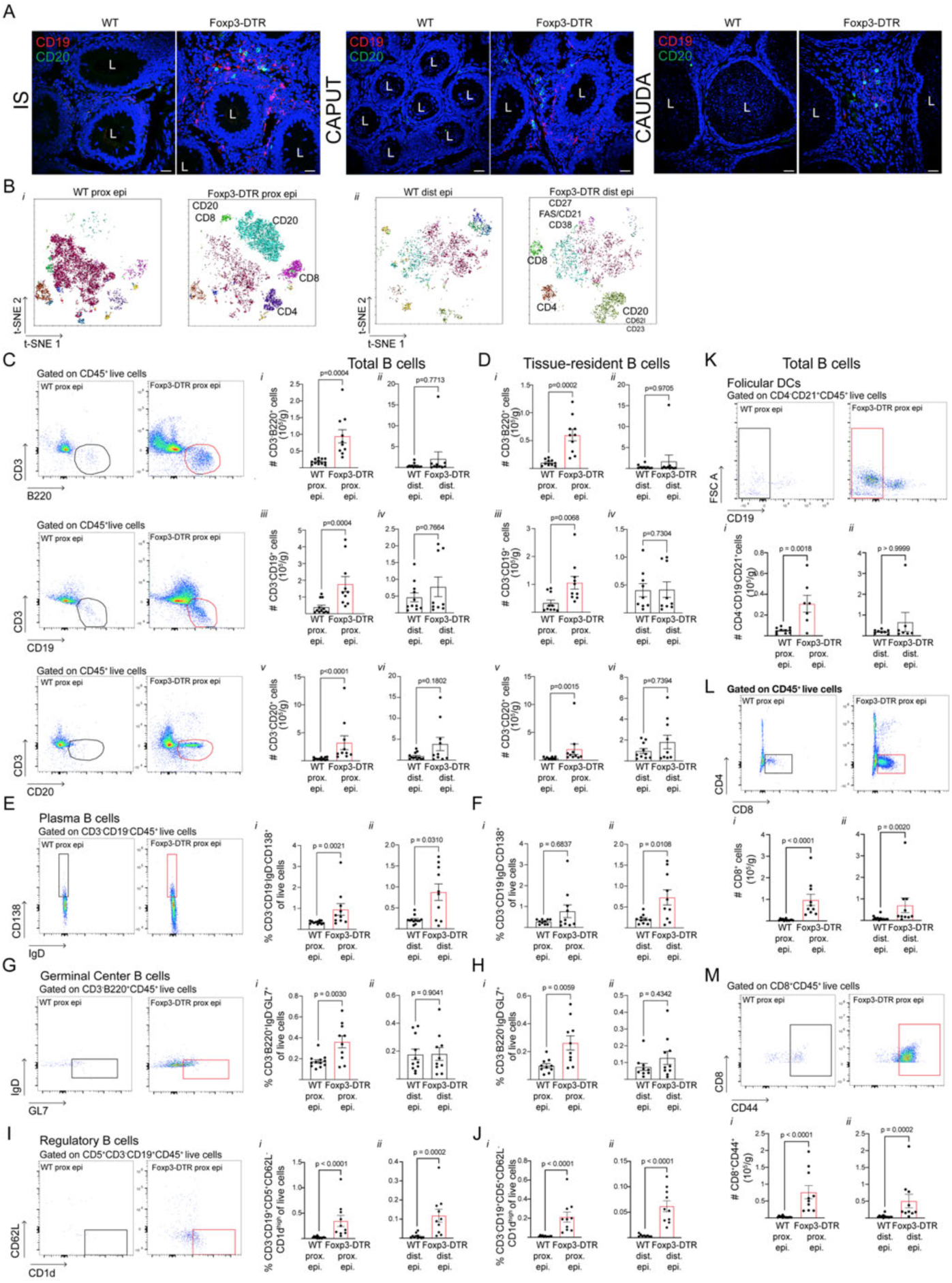
B cell analysis in the epididymis of Foxp3-DTR mice 2 weeks after Treg depletion. **A)** Immunolabeling of CD19 (red) and CD20 (green) in the IS, caput, and cauda regions of DT-injected WT and Foxp3-DTR mice. **B)** tSNE and FlowSOM analysis of proximal (*i*) and distal (*ii*) epididymides from WT and Foxp3-DTR mice. **C)** Flow cytometry gating strategy and absolute count (#) of total B cells (CD3^-^B220^+^, CD3^-^CD19^+^, and CD3^-^CD20^+^ cells) per gram of proximal and distal epididymis. **D)** Absolute count (#) of tissue-resident B cells (CD3^-^B220^+^, CD3^-^CD19^+^, and CD3^-^CD20^+^ cells) per gram of proximal and distal epididymis. Flow cytometry gating strategy and relative abundance of total **(E)** and tissue-resident **(F)** Plasma B cells (CD3⁻CD19⁺IgD⁻CD138⁺ of live cells) in the proximal and distal epididymides. Flow cytometry gating strategy and relative abundance of total **(G)** and tissue-resident **(H)** Germinal center (GC) B cells (CD3⁻B220⁺IgD⁻GL7⁺ of live cells) in the proximal and distal epididymides. Flow cytometry gating strategy and relative abundance of total **(I)** and tissue-resident **(J)** Regulatory B cells (Bregs, CD3^-^ CD19^+^CD5^+^CD62L^-^CD1d^hi^ of live cells) in the proximal and distal epididymides. **K)** Flow cytometry gating strategy and relative abundance of total Follicular Dendritic cells (FDC, CD4^-^CD19^-^CD21^+^ of live cells) in the proximal and distal epididymides. Flow cytometry gating strategy and absolute count (#) of total CD8^+^ T cells (CD4^-^CD8^+^) (**L**) and activated CD8^+^ T cells (CD4^-^CD8^+^CD44^+^) (**M**) per gram of proximal and distal epididymis. Nuclei are labeled with DAPI (blue). Bars: 20 μm. Data were analyzed using Student’s t-test (C*i*, D*i*, F*ii*, G*i*, H*i*, K*i*), or Mann-Whitney test (*Cii-vi*, *Dii-vi*, E, F*i*, G*ii*, H*ii*, I, J, K*ii*, L, M). Data are shown as means ± SEM.

We also observed a higher abundance of total, tissue-resident, and local-circulating CD3^-^ B220^+^IgD^-^GL7^+^ B cells, indicative of GC, in the Treg-depleted proximal epididymis (Fig. 4G-H, Suppl. Fig. 10E*i*), but not in the total and tissue-resident cells of the distal region (Fig. 4G-H, Suppl. Fig. 11E*ii*). In the testis, only resident GC B cells expanded 2 weeks after Treg depletion (Suppl. Fig. 11I-K). Moreover, B regulatory cells (Bregs, CD3^-^CD19^+^CD5^+^CD62L^-^CD1dhi) increased in both proximal and distal epididymal regions (Fig. 4I-J, Suppl. Fig. 10F), while only total and tissue-resident Bregs were elevated in the testis (Suppl. Fig. 11L-N). Alongside the expansion of B cells, we noted infiltration of follicular DC (FDCs, CD4^-^CD19^-^CD21^+^) in the proximal epididymis and testis 2 weeks post-Treg ablation (Fig. 4K*i*, Supp. Fig. 10G*i*, *iii,* Suppl. Fig. 11O-Q), with no changes in the distal regions (Fig. 4K*ii*, Supp. Fig. 10G*ii*, *iv*). Remarkably, CD8^+^ and activated CD8^+^CD44^+^ T lymphocytes also raised in both the epididymis and testis (Fig. 4L-M, Suppl. Fig. 10H-I and Suppl. Fig. 11R-W).

### Chronic inflammation in the testis and epididymis 8 weeks after Treg ablation

The immune response was assessed in the epididymis and testis 8 weeks post-DT treatment to examine the long-term effects of tolerance disruption using *in vivo* staining of circulating CD45^+^ cells (Fig. 5A). Total CD45^+^ cells were prominent in the distal epididymis and testis (Fig. 5B, Suppl. Fig. 14B*ii*). Additionally, local-circulating CD45^+^ cells were found to be elevated in both the proximal and distal epididymis, as well as in the testis (Suppl. Fig. 12A*iii-iv*, Suppl. Fig. 14B*iv*). Tissue-resident CD45^+^ cells were also amplified in the DT-treated Foxp3-DTR testis (Suppl. Fig. 14B*iii*). t-SNE and FlowSOM analysis successfully identified 12 distinct cell subtypes in the epididymis (Fig. 5C-D) and testis (Suppl. Fig. 14A). While the most dramatic immune cell landscape changes were observed in the proximal epididymis 2 weeks post-DT, by 8 weeks, these changes were more pronounced in the distal regions. Interestingly, 8 weeks after depletion, we observed a Treg expansion (total, resident, and circulating) in the epididymis and the testis (Fig. 5E*ii*; Suppl. Fig. 12B*i-iv*; Suppl. Fig. 14C*ii-iv*). The Treg rebound was also evident by confocal microscopy (Fig. 5F, green, arrows). In the distal regions, a surge was observed only in the tissue-resident CD4^+^Foxp3^-^ cells. (Suppl. Fig. 12C*ii*). However, in the proximal epididymis (Fig. 5E*iii* and Supp. Fig. 12C*i, iii*) and testis (Suppl. Fig. 14C*v-vii*), total, resident, and circulating CD4^+^Foxp3^-^ cells were greater compared to controls. Monocyte (CD11b^+^Ly6G^-^Ly6C^+^) and macrophage (CD11b^+^Ly6G^-^Ly6C^-^F4/80^+^) infiltration previously noted 2 weeks after Treg depletion (*5*), had been resolved in the epididymis (Supp. Fig. 13A*i* and B*i*) at this time point. In the testis, monocyte infiltration remained elevated (Suppl. Fig. 13A*ii*), whereas macrophage infiltration was fully resolved (Suppl. Fig. 13A*ii*, D). Conversely, luminal-reaching projections of MPs persisted at a high level (Suppl. Fig. 13C) in the proximal epididymis, indicating sustained antigen sensing, capture, and processing activity.

**Figure 5.**
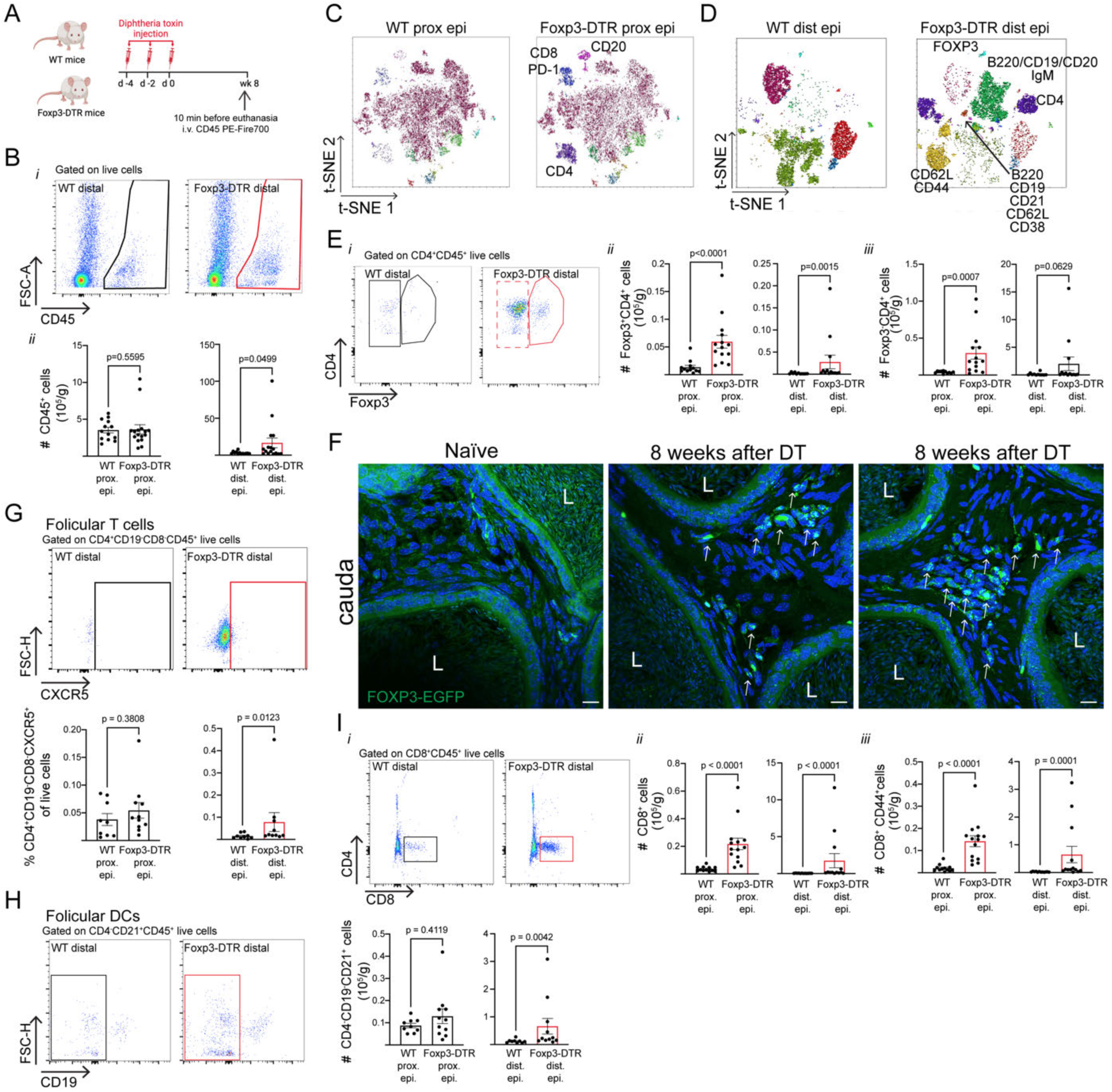
T cell and DC infiltration in the epididymis of DT-injected Foxp3-DTR mice 8 weeks after Treg ablation. **A)** Diagram of Treg depletion protocol, *in vivo* staining of CD45^+^ circulatory cells, and subsequent analysis 8 weeks after DT treatment. **B)** Flow cytometry gating strategy and absolute count (#) of total CD45^+^ cells per gram of proximal and distal epididymis. tSNE and FlowSOM analysis of proximal (**C**) and distal (**D**) epididymides from WT and Foxp3-DTR mice. **E)** Flow cytometry gating strategy (*i*) and absolute count (#) of total Tregs (CD4^+^Foxp3^+^) (*ii*) and T CD4^+^ cells (CD4^+^Foxp3^-^) (*iii*) per gram of proximal and distal epididymis. **F)** Confocal microscopy of distal regions of Foxp3-DTR-eGFP mice. Notice the Treg Foxp3^+^ (green, arrows) cell accumulation in the interstitium of the DT-treated Foxp3-DTR mice. **G)** Flow cytometry gating strategy and relative abundance of T follicular helper cells (Tfh, CD4^+^CD19^-^CD8^-^CXCR5^+^ of live cells) in the proximal and distal epididymides. **H)** Flow cytometry gating strategy and absolute count (#) of total follicular dendritic cells (FDC, CD4^-^CD19^-^CD21^+^ live cells) per gram of proximal and distal epididymides. **I)** Flow cytometry gating strategy (*i*), absolute count (#) of total CD8^+^ T cells (CD4^-^CD8^+^) (*ii*) and activated CD8^+^ T cells (CD4^-^CD8^+^CD44^+^) (*iii*) per gram of proximal and distal epididymis. Nuclei are labeled with DAPI (blue). L: Lumen. Bars: 20 μm. Data were analyzed using the Mann-Whitney test. Data are shown as means ± SEM.

Strikingly, total, tissue-resident and -circulating follicular helper T cells (Tfh, CD4^+^CD19^-^ CD8^-^CXCR5^+^) were higher only in the distal epididymis (Fig. 5G and Supp. Fig. 12D*ii*, *iv*) and in the local circulation of the testis (Supp. Fig. 14D*iv*). In contrast, total and tissue-circulating FDCs (CD4^-^CD19^-^CD21^+^) were found in the distal and testis 8 weeks after DT treatment (Fig. 5H, Supp. Fig. 12E*iv* and Supp. Fig. 14E*ii, iv*). Additionally, in the epididymis (Fig. 5I*i-iii*, Suppl. Fig. 12F*i-iv,* G*i-iv*) and testis (Suppl. Fig. 14F*ii-vii*), the total, tissue-resident, and local-circulating CD8^+^ and activated CD8^+^CD44^+^T lymphocyte counts were elevated at this time point.

Analyzing the B cell diversity of the distal epididymis 8 weeks after Treg elimination, 8 clusters with different phenotypes to the ones defined at 2 weeks were found (Fig. 6A-B and Suppl. 15A). In particular, the total and local-circulating B220^+^, CD19^+^, and CD20^+^ cells and the resident B220^+^ cell populations were higher (Fig. 6B and Supp. Fig. 15C*iii, vi*, D*ii, iv*, E*ii, iv*). However, no differences were observed in B cell counts in the proximal epididymis (Supp. Fig. 15B, Cii, v, Di*, iii*, E*i, iii*). In the testis, besides identifying 8 B cell populations (Supp. Fig. 16A), we found a higher incidence of total, resident, and circulating CD19^+^ and CD20^+^ cells (Suppl. Fig. 16B-D). We also detected that total and circulating plasma B cells amplified in the distal regions and only the circulating ones in the testis (Fig. 6C, Suppl. Fig. 16E). Additionally, GC B cells showed an elevation in the distal regions but not in the testis nor proximal epididymis (Fig. 6D, Suppl. Fig. 15I-K and Supp. Fig. 16F). Significantly, Bregs raised across all compartments in the epididymis, as well as in the total and local-resident Bregs within the testis (Fig. 6E, Suppl. Fig. 16G). Memory B cells, both total and resident, expanded in the testis (Suppl. Fig. 16H) and throughout the epididymis, except for the circulating cells in the proximal region (Fig. 6F). Together, region-specific B cell analysis at 2 and 8 weeks after Treg ablation revealed a shift of naïve B cells to memory and GC B cells, suggesting local B cell maturation in the distal regions after 8 weeks of DT treatment.

**Figure 6.**
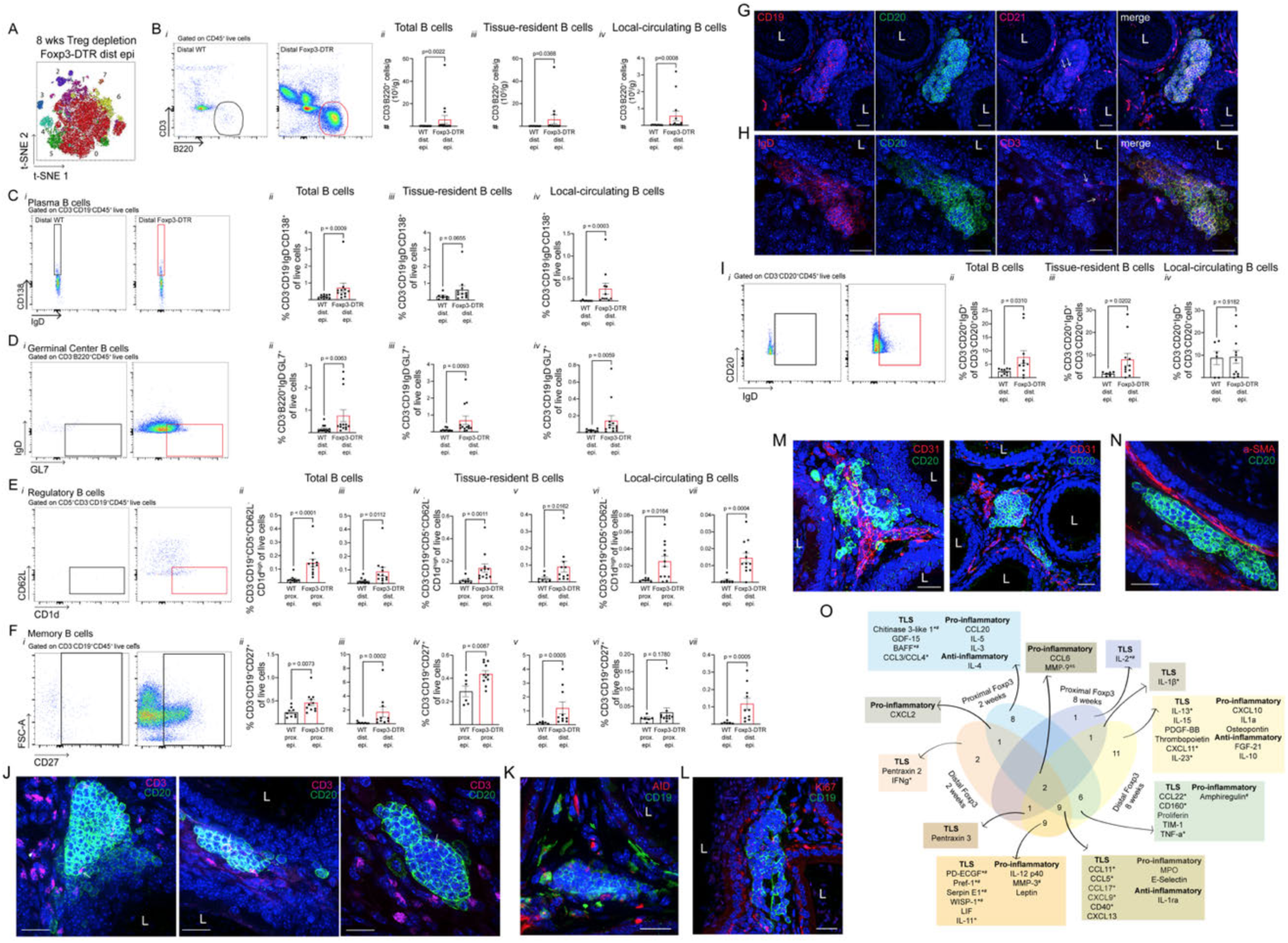
B cell analysis and TLS characterization in the epididymis of Foxp3-DTR mice 8 weeks after Treg depletion. **A)** tSNE and FlowSOM analysis of distal epididymides from Foxp3-DTR mice. **B)** Flow cytometry gating strategy (*i*) and absolute count (#) of total (*ii*), tissue-resident (*iii*), and local-circulating (*iv*) B cells (CD3^-^B220^+^ cells) per gram of distal epididymis. **C)** Flow cytometry gating strategy (*i*) and relative abundance of total (*ii*), tissue-resident (*iii*), and local-circulating (*iv*) plasma B cells (CD3⁻CD19⁺IgD⁻CD138⁺ of live cells) in the distal epididymides. **D)** Flow cytometry gating strategy (*i*) and relative abundance of total (*ii*), tissue-resident (*iii*), and local-circulating (*iv*) Germinal center (GC) B cells (CD3⁻B220⁺IgD⁻GL7⁺ of live cells) in the distal epididymides. **E)** Flow cytometry gating strategy (*i*) and relative abundance of total (*ii, iii*), tissue-resident (*iv, v*), and local-circulating (*vi, vii*) regulatory B cells (B regs, CD3- CD19^+^CD5^+^CD62L⁻CD1d^hi^ of live cells) in the proximal and distal epididymides. **F)** Flow cytometry gating strategy (*i*) and relative abundance of total (*ii, iii*), tissue-resident (*iv, v*), and local-circulating (*vi, vii*) memory B cells (memB cells, CD3^-^CD19^+^CD27^+^) in the proximal and distal epididymides. **G)** Immunolabeling of CD19 (red), CD20 (green), and CD21 (magenta, arrows) in the distal epididymal region of DT-injected Foxp3-DTR mice. **H)** Immunolabeling of IgD (red), CD20 (green), and CD3 (magenta, arrows) in the distal epididymal region of DT-injected Foxp3-DTR mice. **I)** Flow cytometry gating strategy (*i*) and relative abundance of total (*ii*), tissue-resident (*iii*), and local-circulating (*iv*) IgD^+^ CD20^+^ B cells (CD3⁻CD20⁺IgD⁺ of CD20^+^ cells) in the distal epididymides. **J)** Immunolabeling of CD20 (green) and CD3 (magenta) in the distal epididymal region of DT-injected Foxp3-DTR mice. **K)** Immunolabeling of CD19 (green) and AID (red) in the distal epididymal region of DT-injected Foxp3-DTR mice. **L)** Immunolabeling of CD19 (green) and Ki-67 (red) in the distal epididymal region of DT-injected Foxp3-DTR mice. **M)** Immunolabeling of CD20 (green) and CD31 (red) in the corpus and cauda epididymal region of DT-injected Foxp3-DTR mice. **N)** Immunolabeling of CD20 (green) and a-SMA (red) in the distal epididymal region of DT-injected Foxp3-DTR mice. **O)** Venn diagram of the soluble mediators measured in the proximal and distal epididymides at 2 and 8 weeks after DT treatment. Molecules were grouped by general functions or inflammatory abilities such as TLS formation (^&^, cell recruitment, tissue remodeling, angiogenesis capacity) and pro- (*) and anti-inflammatory (^#^) mediators. See also Supp. Fig. 18. Nuclei are labeled with DAPI (blue). L: Lumen. Bars: 20 μm. Data were analyzed using Student’s t-test (E*vi*, F*iv,* I*iv*), or Mann-Whitney test (B*ii-iv*, C*ii-iv*, D*ii-iv*, E*ii-v, vii*, F*ii-iii,v-vii,* I*ii, iii*). Data are shown as means ± SEM.

### Treg depletion triggered TLS formation in the distal epididymis

Confocal microscopy revealed clusters of B cells (CD19^+^ and CD20^+^ positive) resembling TLSs in the corpus and cauda epididymis 8 weeks after Treg depletion (Fig. 6G, H, J-N and Suppl. Fig. 17A-G). Within these TLSs, CD21^+^ follicular B and dendritic-like cells were also present (Fig. 6G). Notably, several B cells within the TLSs were IgD-positive, indicating the presence of mature naïve B cells (Fig. 6H, Suppl. Fig. 17G). The relative abundance of total and tissue-resident IgD^+^ B cells was higher in the distal epididymis (Fig. 6I). We identified various stages of TLS development at 8 weeks (Suppl. Fig. 17G) after Treg depletion. The epididymal TLSs presented features of secondary lymphoid organs such as T cells (CD3^+^ cells, Fig. 6J, Suppl. Fig. 17A and F) surrounding B cell zones (CD19^+^ and CD20^+^ cells). Some TLS cells were positive for AID (activation-induced cytidine deaminase) and Ki67 (Fig. 6K-L and Suppl. Fig. 17B-C). AID plays a crucial role in regulating somatic hypermutation, isotype switching, and the ultimate effector function of Igs (*9, 10*), whereas Ki67 is a well-established proliferation marker. Epididymal TLSs were observed surrounding endothelial cells (CD31^+^ cells, Fig. 6M and Suppl. Fig. 17D) and fibroblasts (α-SMA^+^) (Fig. 6N and Suppl. Fig. 17E). These findings confirm the development of TLSs within the epididymis.

We conducted a multiplex proteome profiling assay to further explore the molecular players involved in the immune response triggered in the different epididymal regions during Treg depletion. This analysis revealed significant differences in mediator abundances between the proximal and distal regions at 2 and 8 weeks following Treg depletion (Fig. 6O and Suppl. Fig. 18A-B). At 2 weeks, 25 molecules were 2-fold differentially increased in the proximal and 18 in the distal epididymis compared to WT, whereas 1 and 6 mediators were found exclusively in the Foxp3-DTR proximal and distal regions, respectively (Fig. 6O). The 8-week Treg-depleted distal epididymis showed 30 molecules with a 2-fold increase, whereas only 5 mediators were found in the proximal epididymis at the same time point versus control. Furthermore, 9 mediators were found exclusively in the Foxp3-DTR distal regions at this time point (Fig. 6O). Many of the secreted factors play critical roles in signaling pathways associated with immune cell recruitment, displaying both pro-inflammatory and anti-inflammatory profiles and contributing to TLS formation.

CXCL13, detected in both regions at 2 weeks and in distal epididymis at 8 weeks post-DT treatment, is crucial for TLS assembly. It recruits CXCR5-expressing B cells, organizing them into follicular structures within TLSs and maintaining B cell zone architecture through this chemokine-receptor axis (*35, 37*). BAFF, upregulated only in the proximal regions at the shorter time point, is also associated with TLS formation and maintenance by promoting the survival and proliferation of B cells (*38*). Notably, the highest number of upregulated molecules was observed in the distal epididymis at 8 weeks, correlating with the presence of TLSs. Chemokines such as CXCL9, CXCL10, CCL3/CCL4/MIP-1α/β, CCL5/RANTES, CCL17/TARC, CCL20, CCL22, among others, are immune cell attractants and activators that are important to initiate signals that trigger TLS establishment (*10*). Together with pro-inflammatory cytokines such as IL-1β, IL-13, TNF-α, IFNγ, IL-12p40, IL-23, and costimulatory molecules such as CD40/TNFRSF5 that activate DCs and B cells, enhance immune cell activation and B-cell differentiation. Anti-inflammatory molecules, particularly IL-10, FGF-21, and IL-1ra, were observed in the distal regions 8 weeks after Treg depletion. These molecules help inhibit excessive inflammation by counterbalancing the significant inflammatory surge. Amphiregulin, Proliferin, MMP-9, Chitinase-3-like protein-1, and GDF-15, involved in wound healing, tissue repair, and angiogenesis, were consistently detected throughout the epididymis at both time points, actively remodeling the extracellular matrix and facilitating the initiation of TLS formation (*39, 40*). These molecules collectively orchestrated the recruitment, activation, and organization of different immune cells, promoting the establishment and functionality of TLSs in response to chronic epididymitis.

### Chronic inflammation Resulted in Severe Immunological Male Subfertility

Regarding the humoral response at 8 weeks after DT, increased serum and tissue Ig levels against sperm, testicular, and epididymal antigens were detected in the Treg-ablated mice. Isotyping of serum autoantibodies revealed distinct humoral profiles compared to 2 weeks after depletion and depending on the antigen tested. In 8-week Treg-depleted serum, increases were noted in IgM, IgG1, IgG2c, IgG3, and IgA against testicular antigens (Suppl. Fig. 19A), while elevated levels of IgM, IgG1, and IgG2c were displayed against epididymal (Suppl. Fig. 19B), proximal sperm (Suppl. Fig. 19C), and distal sperm antigens (Suppl. Fig. 19D). Similar to the findings 2 weeks after Treg ablation, at 8 weeks post-depletion, we revealed accumulation of IgG2b in the testis (Fig. 7A, D), and of IgM, IgG1, IgG2b, and IgG2c in the epididymis (Fig. 7B, C and Supp. Fig. 20A-C). Suppl. Fig. 20D summarizes the profiles of the detected autoantibodies. No antibody deposition nor sperm conglomeration was seen in the epididymal lumen 8 weeks after Treg ablation. Notably, serum IgG titers significantly decreased compared to serum of 2-week Treg-depleted mice (Fig. 7E). Interestingly, the proximal epididymal epithelium exhibited areas of injury 8 weeks post-DT in Foxp3-DTR mice, but such lesions were not observed in the distal region (Fig. 8A-B). At the same time, the IF of B1 V-ATPase CCs (green) and AQP9 (red) of the epididymis did not show major luminal epithelial defects (Suppl. Fig. 21). Importantly, whereas spermatogenesis was not chronically affected (Suppl. Fig. 13E), we observed reduced distal sperm counts (Fig. 8C). Fertility was severely impacted, as shown by a decreased litter size (Fig. 8E). The decline in sperm motility observed 2 weeks after Treg depletion (*5*) was fully restored by 8 weeks (Fig. 8D). These results unveiled that immune tolerance disruption triggered an early and persistent pro-inflammatory response, adversely impacting the epididymis and testis, culminating in chronic male subfertility.

**Figure 7.**
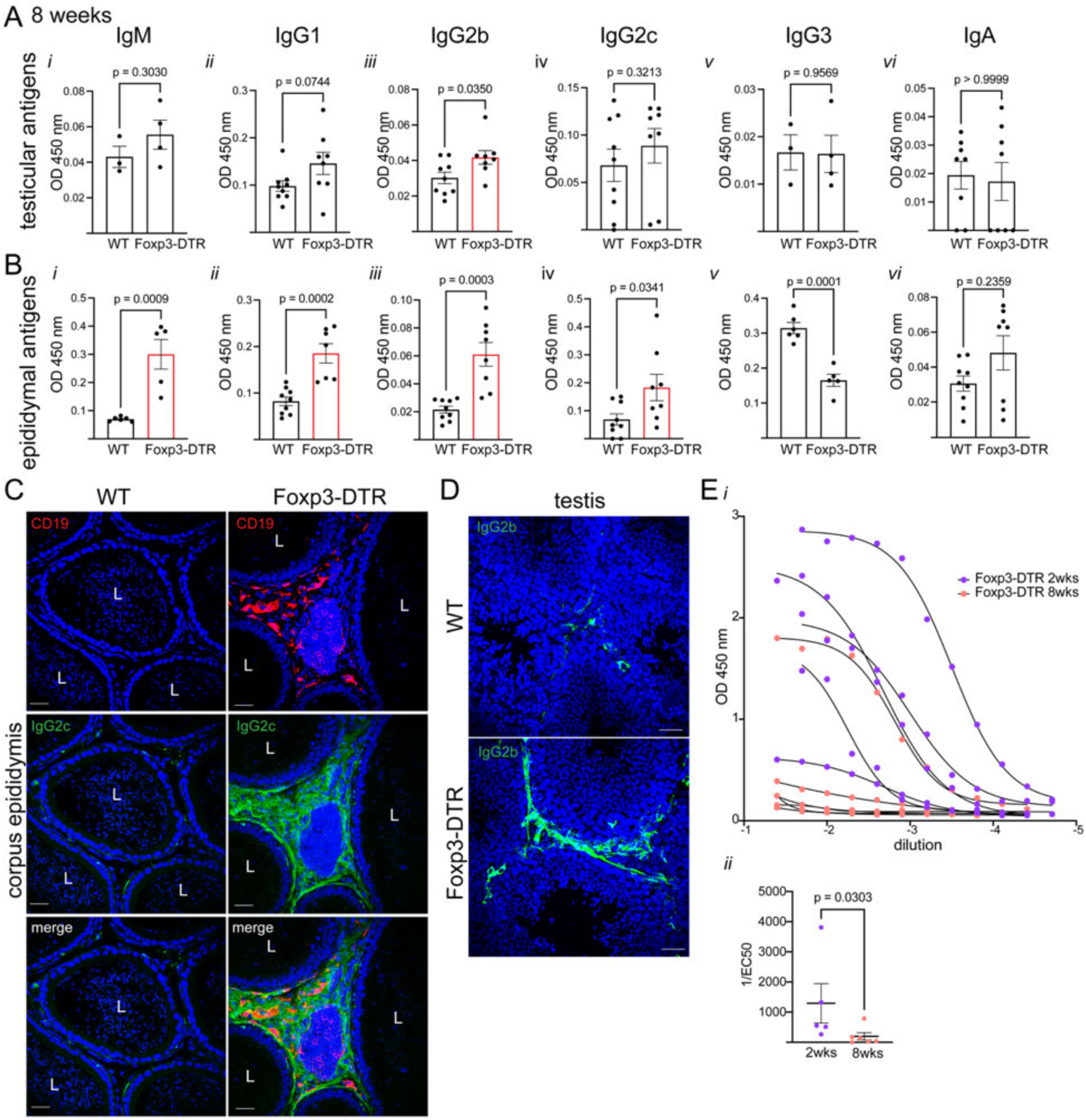
Autoantibody analysis in the testis and epididymis of 8 weeks DT-injected Foxp3-DTR mice. Isotypification of autoantibody levels (IgM, IgG1, IgG2b, IgG2c, IgG3, and IgA) in testicular homogenates against testicular antigens **(A)** and in epididymal homogenates against epididymal antigens **(B)**. **C)** Immunolabeling of CD19^+^ B cells (red) and IgG2c (green) in the corpus region of DT-injected WT and Foxp3-DTR mice. **D)** IgG2b (green) immunolabeling in the testis of DT-injected WT and Foxp3-DTR mice. **E)** Serum total IgG titration at 2- and 8-weeks post Treg ablation by ELISA. *i*) Dose-response curves of optical density vs. serial dilutions of Foxp3-DTR mice sera. *ii*) IgG titers are calculated as the inverse of the EC50 dilution. Nuclei are labeled with DAPI (blue). Bars: 20 μm. L: Lumen. Data were analyzed using Student’s t-test (A*i-iii, v*, B*i-v*), or Mann-Whitney test (A*iv, vi*; B*vi*; E*ii*). Data are shown as means ± SEM.

**Figure 8.**
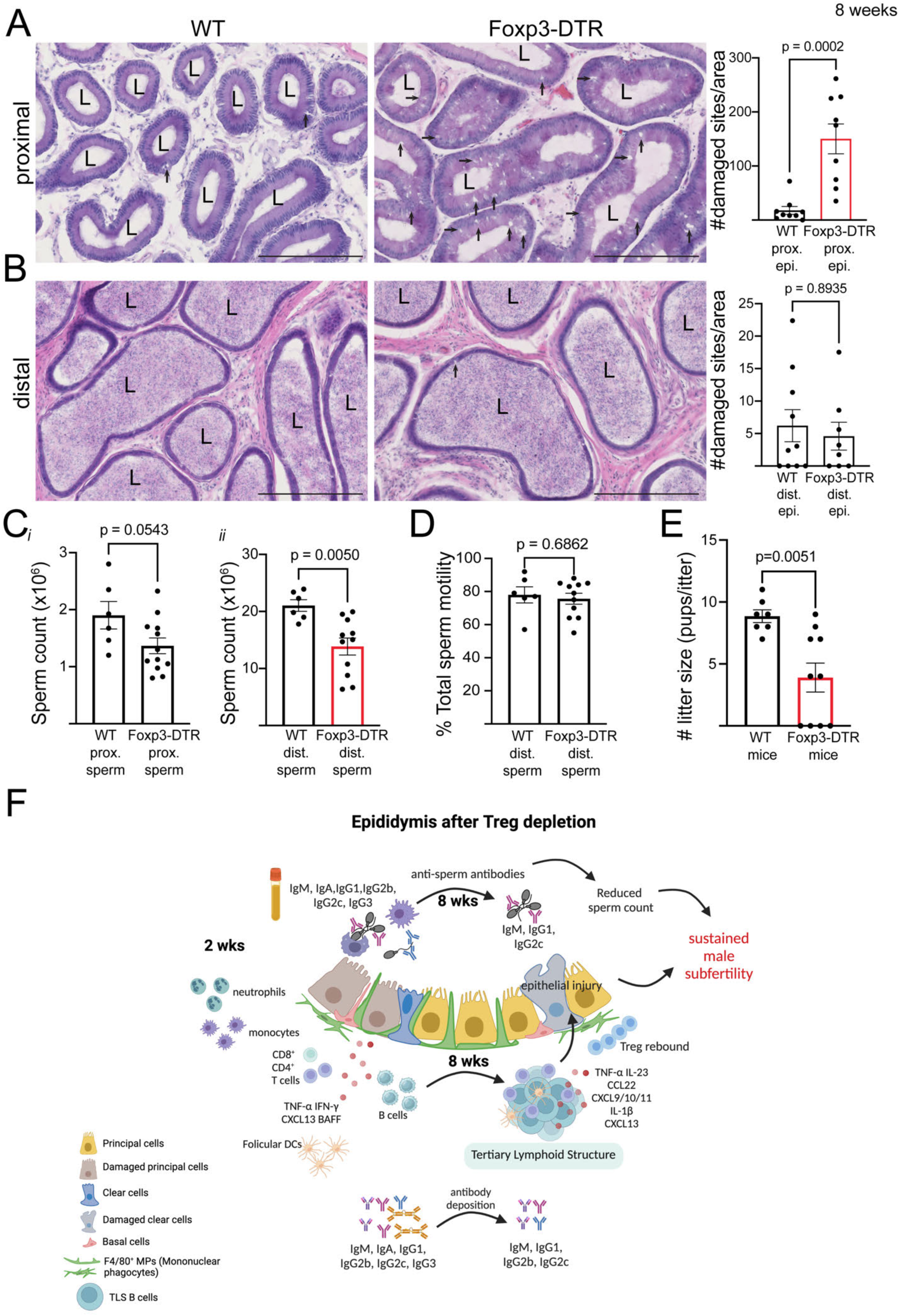
Damaged epididymis and impaired fertility of 8 weeks DT-injected Foxp3-DTR mice. H&E staining of the proximal ((IS) and caput) (**A**) and distal (corpus and cauda) (**B**) epididymal regions of DT-injected WT and Foxp3-DTR mice and quantification of the number of damaged sites normalized per tissue area (µm^2^). Any morphological alteration in epididymal epithelia (arrows) was counted as a damaged site. Each area quantified is represented as a dot. **C)** Computer-assisted sperm analysis (CASA) analysis of proximal sperm cells (*i*) and distal sperm cell (*ii*) counts**. D)** Percentage of total sperm motility analyzed by CASA of distal sperm cells. **D)** Fertility assessment by natural mating as the number of litter size (pups/litter). **F)** Graphical representation of the key findings at 2 and 8 weeks after Treg-depletion in the epididymis. The figure was created with BioRender.com. L: Lumen. Bars: 250 μm, Data were analyzed using Student’s t-test (C, D) or Mann-Whitney test (A, B, E). Data are shown as means ± SEM.

### Chronic inflammation in the epididymis triggers TLS formation, perpetuating impaired male fertility

In our working model (Fig. 8F), within 24 h of Treg depletion, epithelial damage initiates early responses from MPs and epithelial cells (CCs). By 2 weeks, the epididymis shows substantial infiltration of T and B lymphocytes, alongside monocytes, neutrophils, and MPs, particularly concentrated in the proximal regions, as previously described (5). This autoimmune response induces the production of various autoantibody isotypes and the release of pro-inflammatory mediators, including TNF-α and IFNγ. Crucially, both regions of the epididymis secrete factors, such as CXCL13 and BAFF, that facilitate B cell recruitment and maturation and promote TLS formation. At 8 weeks, persistent inflammation in the distal epididymis, characterized by mature and differentiated B and T cell accumulation, supports the establishment of TLSs. The secretion of key mediators, like CXCL9/10/11, IL-23, CCL22, and IL-1β, maintains these ectopic structures, perpetuating a chronic inflammatory environment. At this time, Treg rebound occurs to counteract the inflammation. As a result, the prolonged epithelial injury, along with the presence of ASAs, severely impairs sperm counts, ultimately causing sustained male subfertility.

## Discussion

Chronic inflammation in the epididymis involves complex immune responses that persist over time, causing tissue damage and impaired male fertility (*41, 42*). Tregs are key controllers of immunotolerance and excessive inflammation (*5–7*). Treg dysregulation contributes to autoimmune subfertility, characterized by unique immune responses and the production of autoantibodies targeting sperm and reproductive tissue antigens (*5*). Our current study revealed that B cells are instrumental to this process, producing autoantibodies *in situ* that exacerbate inflammation and epididymal injury, leading to sustained male subfertility. We also uncovered a precise cell-cell communication network between epithelial cells (CCs), lymphocytes, and MPs. In chronic epididymitis, we observed TLSs within the distal regions of the organ. These structures serve as hubs for ongoing immune responses, enabling local B cell activation and autoantibody production, perpetuating inflammation, and highlighting the intricate interplay between myeloid cells, T lymphocytes, Tregs, B cells and autoantibodies, in the pathogenesis of autoimmune impaired fertility.

The epididymis is crucial for sperm maturation and storage; inflammation in this organ can disrupt sperm function (*16, 42, 43*). Improper management of epididymitis may lead to chronic inflammation and long-term complications, including infertility. Mouse models of epididymitis typically involve induced inflammation, such as infection (*Escherichia coli*), non-infectious lipopolysaccharide (LPS)-induced epididymitis, or the injection of immune-stimulating substances like Freund’s adjuvant (*13, 42, 44, 45*). However, the formation of TLSs was not observed in any of these mouse models of epididymitis or orchitis. Our study focused on understanding how epididymitis influences the composition and function of the B and T cell populations. Chronic epididymitis stimulates the expansion of specific subsets of B cells, such as GC, memory B cells, and plasma B cells. These cells may continuously produce antibodies locally, contributing to sustained inflammation and epididymal injury. It is known that B cells undergo class switching and affinity maturation in response to prolonged antigen exposure. Chronic epididymitis, with tissue damage and persistent presence of spermatozoa and autoantigens, may potentially lead to the generation of high-affinity anti-sperm and autoantibodies. Interestingly, we found that the expansion of regulatory B cells, which have immunosuppressive functions, may counteract excessive immune activation and reduce inflammation.

Heterogeneity is a defining characteristic of epididymal segments, underscoring differences in immune responses across regions (*5, 12, 13, 16, 44, 46*). Immune cell diversity and soluble mediator profiles vary markedly between proximal and distal epididymal regions and across 2 and 8 weeks following Treg depletion. The exclusive development of TLSs in distal areas highlights region-specific cues within the epididymis. This diversity reflects the impact of the local microenvironment, or niche, on shaping immune landscapes over time, similar to what is documented in the gut (*47, 48*). The proximal epididymis exhibited a unique immune cell composition and mediator profile compared to the distal region, suggesting that local players— such as epithelial and endothelial cells, fibroblasts, and resident immune cells—drive the recruitment and activation of specific immune populations. Even within the IS, caput, corpus, and cauda, distinct immune cell signatures are present under steady-state conditions (*12, 16, 44, 46, 49, 50*). Characterizing the immunological region-specific landscape of the epididymal microenvironment is essential to advancing patient outcomes in various infertility contexts.

The composition of TLSs varies depending on the tissue type and the specific inflammatory or immune responses (*10*). In the epididymis, TLSs feature diverse immune cells, including B cells, follicular DCs, and T cells, surrounded by endothelial cells and fibroblasts. These structures typically form in perivascular regions in response to disrupted tissue homeostasis. In the distal epididymis 8 weeks after Treg depletion, we observed endothelial cells surrounding the TLSs, elevated levels of pro-angiogenic mediators, and a marked increase in local circulating B cells, suggesting enhanced vascularization of the TLSs. We detected distinct stages of epididymal TLS development: an initial stage marked by dense lymphocyte aggregates, an intermediate stage with defined T and B cell areas, and a mature stage featuring well-organized T- and B-cell zones, accompanied by GCs, class-switched IgD, and fibroblasts (*10*). Early-stage TLSs, marked by B cells, endothelial cells, and fibroblasts, were identified in both proximal and distal epididymis at 2 weeks post-DT treatment. By 8 weeks post-DT, well-defined TLS-like structures with distinct B and T cell zones were observed exclusively in the distal region.

The kinetics of epididymal immune recruitment also differ between the proximal and distal regions, with enhanced B and T cell activation, proliferation, and infiltration initially observed in the proximal region and subsequently, in the distal segments. 2 weeks after Treg ablation, both epididymal regions secreted CXCL13, which promotes TLS formation by attracting T and B cells (*35*). In both regions, there was an upregulation of CD40, which promotes GC formation, Ig isotype switching, and the generation of plasma and memory cells (*51*). Additionally, CXCL2, secreted by MPs, was elevated and plays a crucial role in angiogenesis, a process essential for the maintenance of TLSs (*52*). Another region-specific signature is a more Th2-biased cytokine profile (IL-4, IL-5, TIM-1) as exhibited by the proximal epididymis, in contrast to distal regions, which showed a more Th1-like profile (IL-12 p40, IFNγ). At 8 weeks post-Treg ablation, numerous soluble pro-inflammatory mediators were detected in the distal regions, triggering TLS formation and maintenance. However, in this region, IL-10 and IL-1ra were also upregulated, indicating an anti-inflammatory response within the milieu where the TLSs are located. These contrasting factors highlight the complexity and precision of epididymal immune regulation. Understanding these region-specific, time-dependent variations is essential for unraveling local immune responses and their implications for tissue pathophysiology.

While TLSs have been identified in testicular tumors (*38, 53*), no data exists regarding these structures in epididymal cancer. Although testicular tumors are relatively common malignancies in young men, epididymal cancers are extremely rare, typically arising as secondary tumors from testicular origins. The available literature on epididymal cancers is limited, consisting primarily of case reports (*54, 55*). Notably, the testis—especially the Sertoli cells—can protect allografts from rejection. Key contributors to this phenomenon include various anti-inflammatory mediators, Tregs, and M2-like MPs (*56*). Further studies are needed to understand why TLSs form in the epididymis but not in the testis after Treg depletion. Comparative analyses of TLS composition induced by different conditions will help uncover shared and unique mechanisms across different reproductive tissues.

Tregs play a context-specific role in TLS formation. In a mouse lung adenocarcinoma model, Treg absence increases CD4^+^ and CD8^+^ T cell infiltration, particularly in TLSs, while in pemphigus skin TLSs, Tregs drive CXCL13 production through interaction with T cells, which in turn promotes TLS development. Breg involvement in TLS remains understudied (*10*). Our findings suggest that CD8^+^ cells may contribute to epididymitis-related damage, as dysregulated CD8^+^ T cells could target sperm or epididymal tissue, potentially causing reduced fertility (*57*). CD8^+^ cell prevalence also shifts in males with ASAs (*58, 59*). Multiple memory B cell phenotypes have been identified in TLSs, with IgG1^+^ B cells enriched in pancreatic cancer (*10*). Notably, IgG1, IgG2b, IgG2c, and IgM persist in the epididymis 8 weeks post-Treg depletion, indicating ongoing local antibody production. The distinct autoantibody profiles detected may be linked to specific stages of autoimmunity progression. These autoantibodies could mediate sperm opsonophagocytosis, antibody-dependent cellular cytotoxicity, immune complex formation, and complement activation, ultimately leading to sperm cell lysis (*3, 4, 60*). These defects likely contribute to the reduced sperm counts and smaller litter sizes observed after natural mating, 8 weeks post-Treg depletion. Additionally, ASAs may exacerbate the condition by promoting the release of inflammatory mediators, further impairing sperm fertilizing ability (*4, 8, 61*).

After Treg depletion, epididymal damage occurs within the first day and persists for 8 weeks, priming B cell maturation, ASAs production, and formation of TLSs. TLSs also may contribute to the local production of autoantibodies. Infertile patients, as well as most individuals who have undergone vasectomy, present ASAs (*4, 8, 62*). ASAs are associated with reduced sperm motility and concentration, impairing the ability of sperm to penetrate cervical mucus, interact with the egg, and potentially affect early embryonic development (*3, 4, 8, 58, 63*). The presence of IgG and IgA ASAs in the seminal plasma of patients with fertility challenges, along with ASA recognition of the head and tail of human sperm, suggests shared mechanistic pathways between murine and human pathological processes.

Identifying the specific targets of ASAs remains a critical area of research, as doing so could lead to better diagnosis and treatment of ASA-mediated immune infertility and the development of new immune-contraceptive methods. Importantly, the Human Contraception Antibody, an IgG1 anti-sperm monoclonal antibody, is currently under development as a novel non-hormonal contraceptive (*64*). Several sperm antigens have been identified in ASA from patients suffering from immune infertility (*4*). Our work focused on pinpointing the targets of ASAs and expanded the list of sperm antigens recognized by autoantibodies in the proximal and distal epididymal fluids and serum.

The breakdown of immunotolerance in the epididymis and testis triggers chronic inflammation and local autoimmune responses, marked by autoantibody deposition, region-specific immune signatures, soluble mediator secretion, and lymphocyte infiltration. This immune disruption impairs sperm function and drives prolonged male subfertility. Epididymal TLSs may further exacerbate inflammation by driving local antibody production against sperm cells. Our study identifies key molecular mediators involved in epididymal TLS formation and suggests that unexplained male infertility may be linked to immune dysregulation—characterized by pro-inflammatory cell recruitment, autoantibodies, and elevated soluble mediators. Overall, we provide an in-depth understanding of B- and T-cell diversity dynamics and TLS formation during epididymitis to develop precision-targeted therapies for infertility and chronic inflammation.

## Methods

### Mouse model

Female C57BL/6-Tg(Foxp3 DTR/EGFP)23.2Spar/Mmjax (Foxp3-DTR mice) and male C57BL/6 WT mice were acquired from Jackson Laboratory and bred at the Massachusetts General Hospital (MGH) animal facility to produce Foxp3-DTR or WT (control) male littermates(*5*). Foxp3-DTR transgenic mice express the diphtheria toxin receptor (DTR) fusion protein controlled by the endogenous forkhead box P3 (Foxp3) promoter/enhancer regions. Male mice aged between six to forty weeks were used for all experiments. Animal procedures adhered to the National Institutes of Health (NIH) Guide for the Care and Use of Laboratory Animals and were approved by the MGH Subcommittee on Research Animal Care. Similar to our previous studies(*5*), six-week-old male littermates, both Foxp3-DTR and WT, received intraperitoneal injections of diphtheria toxin (DT) from *Corynebacterium diphtheriae* (D0564, Sigma-Aldrich, St. Louis, MO) dissolved in phosphate-buffered solution (PBS) at a dosage of 40 μg/kg of body weight to deplete Tregs. DT injections were administered on days 0, 2, and 4, and mice were assessed 2 and 8 weeks after the final DT injection. DT-injected WT mice served as controls. Furthermore, some Foxp3-DTR and WT mice received a single DT injection to verify the depletion of Tregs after 24 hours of treatment.

### Mouse in vivo staining

The *in vivo* staining of circulating CD45^+^ cells was performed following a reported protocol(*15*). Briefly, mice were intravenously injected with anti-mouse CD45-PE or CD45-PE-Fire-700 (3 µg/100 µL saline) 10 min before euthanasia. As detailed below, tissue was collected for flow cytometry staining and fixation. The gating strategy used to distinguish circulating from resident CD45^+^ cells is shown in Figure 1D.

### Mouse testis and epididymis collection

Mice were anesthetized using a mixture of isoflurane (2%) and air (Baxter, Deerfield, IL) and then perfused through the left cardiac ventricle with PBS until the organs were free of blood (for ELISA and soluble mediator arrays). Subsequently, the epididymis and testes were dissected for further analysis. For tissues requiring fixation, perfusion continued with a 4% paraformaldehyde (PFA; 15714-S, Paraformaldehyde 32% Solution EM grade, Electron Microscopy Sciences) fixative for 7 min. Following harvesting, the organs were immersed in 4% PFA for 4 hours. For *in vivo* CD45 staining, animals were not subjected to PBS perfusion. Instead, the epididymides and testes were excised after euthanasia.

### Mouse sperm collection

Sperm were retrieved from the proximal (initial segments (IS) and caput) and distal (cauda) regions of the epididymis in Foxp3-DTR and WT littermates following DT injections. Sperm were obtained by excising the respective areas in modified Human Tubal Fluid (HTF) medium (#90126, FUJIFILM Irvine Scientific, Inc., Santa Ana, CA) enriched with 0.3 % of Bovine Serum Albumin (BSA; A0281, Sigma-Aldrich). After a 15-min incubation at 37°C, sperm were examined using CASA (computer analysis system assay) and/or fixed with 4% PFA for 10 min at room temperature.

### Mouse serum collection

Blood was drawn from the left ventricle of the mouse heart into BD Microtainer® blood collection tubes (365967, Becton, Dickinson and Company, Franklin Lakes, NJ). After allowing the samples to sit at room temperature for 30 min, they were centrifuged at 4,000g for 10 min to isolate the serum.

### Mouse tissue processing

Cleaned, perfused, and frozen tissues were thawed in PBS supplemented with a protease inhibitor cocktail (#04693159001, cOmplete™ Mini EDTA-free Protease Inhibitor Cocktail, Roche, Basel, Switzerland). Tissues were minced with scissors, homogenized with a pestle, and incubated on ice for 30 min. The supernatant was collected after centrifuging for 20 min at 10,000g. Samples were either used directly for ELISA or stored at −80°C until further use. For cytokine/chemokine kit measurements, the same procedure was followed. Tissues (4 proximal and 4 distal epididymes from DT-treated WT and Foxp3-DTR mice) were thawed and processed in PBS with Triton X-100 (Sigma-Aldrich) as stated in the manufacturer’s protocol (ARY028, Proteome Profiler Mouse XL Cytokine Array R&D Systems).

### Mouse epididymal fluid collection

Epididymal fluids were retrieved from the proximal (initial segments (IS) and caput) and distal (cauda) regions of the epididymis in Foxp3-DTR and WT littermates following DT injections. Fluids were obtained by excising the respective areas in PBS pH 7.4. After a 15-min incubation at 37 °C, tissues were discarded, and samples were centrifuged for 5 min at 500g to discard sperm cells. Supernatants were stored at −80C until use.

### Confocal microscopy

Similar to previous studies (*5, 49*), the fixed organs were preserved in 30% sucrose in PBS (with 0.02% NaAzide) for at least 48 h at 4°C and then embedded in Tissue-Tek OCT compound (Sakura Finetek, Torrance, CA). Tissue sections, either 5 or 25 µm in width, were prepared using a Reichert Frigocut microtome and mounted onto Fisherbrand Superfrost Plus microscope slides (Fisher Scientific, Pittsburgh, PA). Alternatively, sperm samples were fixed for 10 min in 2% PFA at room temperature and then washed twice with PBS before being smeared onto slides.

Primary antibodies utilized were rat anti-mouse-F4/80 (*5*), chicken anti-V-ATPase B1 subunit(*5*), rabbit anti-AQP9(*5*), goat anti-mouse-IgA (1mg/mL, 62-6720, Invitrogen), goat anti-mouse-IgG1 (1mg/mL, A10551, Invitrogen), goat anti-mouse-IgG2b (1mg/mL, M32407, Invitrogen), goat anti-mouse-IgG2c (0.8mg/mL, 115-035-208, Jackson ImmunoResearch), goat anti-mouse-IgG3 (M32707, Invitrogen), donkey anti-mouse-IgM (0.8mg/mL, 715-035-020, Jackson ImmunoResearch), rabbit anti-mouse-CD19 (0.2mg/mL: sc-8500-R, Santa Cruz), rat anti-mouse-CD21/CD35 (0.5mg/mL, 14-0211-81, Invitrogen), Armenian Hamster anti-mouse- CD3 (0.5mg/mL, 100303, Bioegend), rat anti-human/mouse-AID (0.5mg/mL, 14-5959-80, Invitrogen), rabbit anti-alpha smooth muscle actin (SMA) (0.2mg/mL, ab5694, Abcam), rabbit anti-mouse-CD31 (ab28364, Abcam), rabbit anti-Ki67 (0.031 mg/mL, MA5-14520, Thermofisher Scientific), rat anti-mouse-IgD (0.5mg/mL, 405702, Biolegend), goat anti-mouse-CD20 (0.5mg/mL, sc-7735, Santa Cruz), and donkey anti-human-IgG (1.5mg/mL, 709-545-149, Jackson ImmunoResearch). The secondary antibodies used were Alexa Fluor 488 goat anti-mouse IgG (20 μg/ml; A-11001, Invitrogen), Alexa Fluor 647 donkey anti-rat IgG (3 μg/ml; 712-606-150, Jackson ImmunoResearch, West Grove, PA), Alexa Fluor 488 donkey anti-mouse IgG (20 μg/ml; A21202, Invitrogen), Alexa Fluor 488 donkey anti-chicken IgY (15 μg/ml; 703-545-155, Jackson ImmunoResearch), Alexa Fluor 647 donkey anti-rabbit IgG (1.5 μg/ml; 711-606-152, Jackson ImmunoResearch), Alexa Fluor 488 goat anti-rabbit IgG (1:800 dilution; 111-545-144, Jackson ImmunoResearch), DyLight 649 goat anti-hamster (STAR104D649, AbD Serotec), FITC donkey anti-human IgG (15 μg/ml; 709-545-149, Jackson ImmunoResearch) and Cy3 donkey anti-rabbit IgG (7.5 μg/ml; 711-165-152, Jackson ImmunoResearch).

Different antigen retrieval methods were employed based on the antibody used: a 4-min treatment with PBS containing 1% SDS (for V-ATPase B1, CD19, CD20 CD3, and AQP9 antibodies), or PBS containing 1% SDS and 0.1% Triton (for anti-F4/80, anti-IgG and anti-IgA antibodies). Slides were then blocked for 30 min at room temperature in PBS containing 1% BSA and incubated with primary antibodies for 18 hours at 4 °C. All antibodies were diluted in DAKO medium (DAKO, Carpinteria, CA). Mounted slides were treated with SlowFade Diamond Antifade Mounting medium (Thermo Fisher Scientific, S36963) containing the DNA marker DAPI. Negative control slides underwent incubation with secondary antibodies alone for immunostaining.

### H&E evaluation

Sections (5 µm in width) were stained with Hematoxylin Solution, Harris Modified (HHS32, Sigma-Aldrich), and subsequently counterstained with Eosin Y Solution (HT110132, Sigma-Aldrich). The stained slides were digitally scanned using a NanoZoomer 2.0RS digital scanner (Hamamatsu, Japan).

### Flow cytometry analysis

The proximal and distal segments of the epididymis and testis were mechanically minced and enzymatically digested for 30 min at 37°C in RPMI 1640 medium containing collagenase type I (0.5 mg/mL) and collagenase type II (0.5 mg/mL), following established protocols(*5*). The resulting suspensions were filtered through a 70 μm nylon mesh strainer, rinsed with 2% fetal bovine serum (FBS) in PBS with 2 mM EDTA, and centrifuged for 5 min at 400 g. The pellet was treated with ACK lysing buffer (#A10492-01, Gibco, Grand Island, NY) for 1 min, followed by centrifugation for 5 min at 400 g. Cell suspensions were then incubated for 30 min with anti-mouse antibodies (1:500 dilution) against CD45 Brilliant Violet 711 (0.2 mg/ml; Clone 30-F11, 563709, BD Biosciences, San Jose, CA), CD45 PE-Fire^tm^700 (0.2 mg/ml; Clone 30-F11, 103177, BioLegend), CD45 PE (0.5 mg/ml; Clone 30-F11, 103105, BioLegend), CD45 BUV395 (0.2 mg/ml; Clone 30-F11; 565967, BD Biosciences), Ly6G PE (0.2 mg/ml; Clone 1A8, 551461, BD Biosciences), F4/80 PE/Cyanine7 (0.2 mg/ml; Clone BM8, 123113, BioLegend, San Diego, CA), Ly6C FITC (0.5 mg/ml; Clone AL-21; 553104, BD Biosciences), MHC Class II (I-A/I-E) Alexa Fluor 700 (0.2 mg/ml; Clone M5/114.15.2, 56-5321-82, Thermo Fisher Scientific, Waltham, MA), CD4 PE (0.2 mg/ml; Clone RM4-5, 100511, BioLegend), CD4 BV480 (0.2 mg/ml; Clone RM4-5; 565634, BD Biosciences), CD44 BUV737 (0.2 mg/ml; Clone IM7; 612799, BD Biosciences), CD11b APC/Cyanine7 (0.2 mg/ml; Clone M1/70, 101225, BioLegend), CD1d BUV805 (0.2 mg/ml; Clone IB1; 741965, BD Biosciences), CD19 BV750 (0.2 mg/ml; Clone 1D3; 747332, BD Biosciences), IgD BUV615 (0.2 mg/ml; Clone 217-170, 751474, BD Biosciences), CD138 BV510 (0.2 mg/ml; Clone 281-2, 563192, BD Biosciences), CD3 BV421 (0.2 mg/ml; Clone 17A2, 100227, BioLegend), CD20 PE (0.2 mg/ml; Clone SA275A11, 150409, BioLegend), B220 Pacific Blue (0.5 mg/ml; Clone RA3-6B2, 103230, BioLegend), FAS APC/Fire^TM^810 (0.2 mg/ml; Clone SA367H8, 152623, BioLegend), CD5 Alexa Fluor^®^594 (0.5 mg/ml; Clone 53-7.3, 100632, BioLegend), GL7 PerCP/Cyanine5.5 (0.2 mg/ml; Clone GL7, 144609, BioLegend), CD8 FITC (0.5 mg/ml; Clone 53-6.7, 100706, BioLegend), CXCR5 BV605 (0.1 mg/ml; Clone L138D7, 145513, BioLegend), CD62L PE/Cyanine5 (0.2 mg/ml; Clone MEL-14, 104410, BioLegend), CD21/CD35 Alexa fluor^®^647 (0.5 mg/ml; Clone 7E9, 123423, BioLegend), CD38 Alexa Fluor^®^ 700 (0.5 mg/ml; Clone 90, 102741, BioLegend), PD-1 APC/Cyanine7 (0.2 mg/ml; Clone 29F.1A12, 135223, BioLegend), CD23 PerCP (0.1 mg/ml; Clone #011, 50695-R011, SinoBiological Inc.), CD27 PerCP-eFluor^TM^710 (0.2 mg/ml; Clone LG.7F9, 46-0271-80, Invitrogen), diluted in 2 % FBS in PBS (with BD Horizon Brilliant Stain buffer (BD Biosciences). DAPI (62248, ThermoScientific), LIVE/DEAD™ Fixable Blue Dead Cell Stain Kit, for UV excitation (L23105, Invitrogen) or LIVE/DEAD™ Fixable Yellow Dead Cell Stain Kit, for 405 nm excitation (L34967, Invitrogen) were used as viability markers.

For intranuclear staining of the Foxp3 and intracellular staining of IgM and IgG, suspensions were incubated fixed for 20 min with 1x Fixation/Permeabilization working solution and washed with 1x Permeabilization Buffer, following the manufacturer’s instructions (#00-5523-00, Thermo Fisher Scientific). Fixed cells were then incubated overnight with the anti-Foxp3-APC (0.2 mg/ml; Clone FJK-16s, 17-5773-82, ThermoFisher Scientific), IgM BV786 (0.2 mg/ml; Clone R6-60.2, 564028, BD Biosciences) IgG PE/Cyanine7 (0.2 mg/ml; Clone Poly4053, 405315, BioLegend) diluted in 1x Permeabilization Buffer (00-5523-00, Thermo Fisher Scientific) containing BD Horizon Brilliant Stain buffer (563794, BD Biosciences)(102). After incubation, cells were washed with 2% FBS in PBS and filtered through a 40 μm cell strainer.

Flow cytometry data were collected using the BD FACSAria II flow cytometer (BD Biosciences) or AURORA flow cytometer (Cytek® Aurora System) and interpreted using FlowJo software version 10.8.1 (BD Biosciences). Control experiments including negative controls (unstained cells from each organ) and fluorescence minus one (FMO) controls were conducted to establish gating strategies, following established procedures. t-distributed Stochastic Neighbor Embedding (tSNE) and FlowSOM analysis were applied to flow data, enabling detailed characterization of cell populations. tSNE was used for dimensionality reduction to visualize high-dimensional flow cytometry data projected into a lower-dimensional space and FlowSOM self-organizing mapping clusters cells based on their phenotypic similarities, for predicting distinct cell populations.

### Enzyme-Linked Immunosorbent Assay (ELISA)

Proteins were extracted from the testis, epididymis, and sperm of WT mice and from healthy human sperm donors, using a buffer consisting of RIPA buffer (R26200-125.0, RPI Research Products International, Mt. Prospect, IL) supplemented with a protease inhibitor cocktail (#04693159001, cOmpleteTM Mini EDTA-free Protease Inhibitor Cocktail, Roche, Basel, Switzerland) and a phosphatase inhibitor cocktail (#4906837001, PhosSTOP, Roche) for 20 min at 4 °C. Subsequently, the samples were centrifuged at 10,000 g for 10 min, and the resulting supernatants containing soluble proteins were collected. 96-well plates were coated with tissue/cell proteins (testicular, epididymal, sperm) at a concentration of 6 µg/mL in PBS, pH 7.4, and left overnight at 4°C. The plates were then blocked with 3% BSA in PBS for 1 hour at 37°C before incubating with mouse serum diluted 1/25, tissue homogenates diluted 1/3, epididymal fluids diluted ½ or human seminal fluid diluted 1/5 in 1% BSA in PBS for 2 hours at 37°C. The different isotypes bound to the antigen-coated wells were detected following a 1-hour incubation with horseradish peroxidase (HRP)-anti-mouse IgG (1:5000; 115-035-205, Jackson ImmunoResearch), IgA (1:3000; 62-6720, Invitrogen), IgG1 (1:2000; A10551, Invitrogen), IgG2b (1:5000; M32407, Invitrogen), IgG2c (1:5000; 115-035-208, Jackson ImmunoResearch), IgG3 (1:2000; M32707, Invitrogen), IgM (1:5000; 715-035-020, Jackson ImmunoResearch) and human totIgG (1:10000, 709-035-149, Jackson ImmunoResearch) and IgA (1:10000; 109-035-011, Jackson ImmunoResearch). The enzymatic reaction was initiated with 3,3’,5,5’-tetramethylbenzidine (TMB) substrate (TMB substrate kit, 34021, Thermo Fisher Scientific) and stopped with H_2_SO_4_ 2N. Optical density readings were measured using a microplate reader (Promega GloMax Discover, Madison, WI) at 450 nm.

### Computer Assisted Sperm Analysis (CASA)

Sperm were harvested from the proximal/distal epididymis as described above. After a 15-min incubation at 37°C, proximal/distal sperm were collected and analyzed. Alternatively, distal sperm were obtained and then diluted (1:5 ratio) in HTF medium with 0.3% BSA, followed by a 60-min incubation at 37°C to induce sperm capacitation. Sperm analysis was conducted using Hamilton Thorne’s CASA version 14 (Hamilton Thorne Inc., Beverly, MA). Each sample underwent a minimum of two analyses.

### Assessment of fertility

Male littermates of both Foxp3-DTR and WT strains (8-week post-DT injections) were singly housed with a WT adult female for 5 days. The presence of a copulatory plug confirmed successful mating. Fertility assessment was determined based on litter size, representing the number of pups per litter.

### Immunoprecipitation using anti-sperm antibodies

Serum and proximal and distal epididymal fluids were collected from WT and Foxp3-DTR mice two weeks following DT treatment. These samples were incubated with Dynabeads® Protein G, washed, and then incubated with protein extracts (as previously described) derived from proximal or distal sperm cells of WT naïve mice, following the manufacturer’s protocol (Immunoprecipitation Kit – DynabeadsTM Protein G, 10006D, Invitrogen). Eluates were run on a 10% bis-acrylamide gel (NuPAGETM 10% Bis-Tris Gel, NP0302BOX, Invitrogen), then stained with Coomassie dye R-250 (ImperialTM Protein Stain, 24615, Thermo Scientific), bands were cut and submitted to the Taplin Biological Mass Spectrometry Facility, Harvard Medical School, Building C, Room 523, 240 Longwood Ave. Boston, MA 02115, similar to previously reported (*5*). Results are available in Supp. Tables I and II.

### Proteomic Data Analysis

Proteomic datasets from each experimental group were compared using the VENNY 2.1 Venn Diagram online tool (https://bioinfogp.cnb.csic.es/tools/venny/). Protein abundance was quantified by summing intensity values, calculated as the sum of intensities of all peaks from the same protein. Heat maps were generated using the web-based software Morpheus (https://software.broadinstitute.org/morpheus). The proteomics data generated will be deposited to theProteomeXchange Consortium (http://proteomecentral.proteomexchange.org) via the PRIDE partner repository with the dataset identifier xxx.

### Human sample collection

Human semen samples were obtained from patients undergoing routine semen analysis at the assisted reproduction unit of the Cruces and Galdakao University Hospital of Bizkaia (Spain). Ejaculates were collected by masturbation after 3-5 days of sexual abstinence. All patients gave signed informed consent to the Declaration of Helsinki. Semen parameters and the presence of round cells (RC) were evaluated using the Sperm Class Analyzer® CASA System (Microptics®).

After semen analysis, all samples were classified as normozoospermic or asthenozoospermic according to the World Health Organization guidelines (*1*).

### Human sample processing

Human semen samples were centrifuged at 300 g for 15 min to separate the seminal plasma and sperm fraction. Then, isolated sperm cells were washed with 2 mL of phosphate-buffered saline 1X (PBS 1X). Finally, sperm pellets and seminal plasma fractions were stored at −80°C until usage.

### Statistical Analysis

Data analysis was conducted using GraphPad Prism 10 version 10.3.1 (464) (GraphPad Software, La Jolla, CA). The Shapiro-Wilk test was employed to assess normality. Analysis of variance (F test) was utilized to compare two groups. Parametric tests, including Student’s t-test (two-tailed), were applied. For non-parametric analyses, the Mann-Whitney test (two-tailed) was used. Statistical significance was set at p-values < 0.05. Data were presented as means ± SEM.

## Acknowledgments

The authors thank the Microscopy Core of the Program in Membrane Biology (PMB) (MGH, Boston, MA) and the MGB Molecular Imaging Core (MGH, Charlestown, MA), the Taplin Mass Spectrometry Facility (Cell Biology Department, Harvard Medical School, Boston, MA), and the MGH Pathology Flow Cytometry Facility (MGH, Boston, MA). This work was supported by the National Institutes of Health (grant HD104672-01 to M.A.B.), the MGH Physician/Scientist Development and Claflin Distinguished Scholar Awards (to M.A.B), and the Lalor Foundation (to F.B. and M.L.E), the IBSA Foundation for Scientific Research (to F.B.), and the Ministry of Science and Innovation, Spain, grant numbers PID2020-119949RB-100 and TED2021-132681B-I00, CPP2021-008458, co-founded by the European Union and by Instituto de Salud Carlos III, co-funded by European Union (ERDF/ESF, “Investing in your future”, grant number PI20/01131) to N.S.

## Author Contributions

M.L.E, F.B., and M.A.B. were involved in the study design and conceptualization. M.L.E., F.B., K.O., M.C.A., I.B., A.O., V.D.R., O.U., and A.C. performed the experiments and data analysis. M.L.E., F.B., M.C.A., N.S, and M.A.B. were involved in data interpretation. M.L.E. and M.A.B. wrote the original manuscript. All authors contributed to the writing of the manuscript, made critical comments, and approved the final version.

## Competing Interest Statement

The authors declare no financial or non-financial competing interests.

## Ethics approval

All animal procedures were approved by the Massachusetts General Hospital (MGH) Subcommittee on Research Animal Care and followed the NIH Guide for the Care and Use of Laboratory Animals (National Academies Press, 2011; protocol 2003N000216).

The human study adhered to the ethical principles outlined in the Declaration of Helsinki (1975, revised in 1983) and was approved by The Medicine Research Ethics Committee of the Basque Country, Spain (protocol number: PI2019184).

## Supplementary Material

**Supplementary Figure 1.**
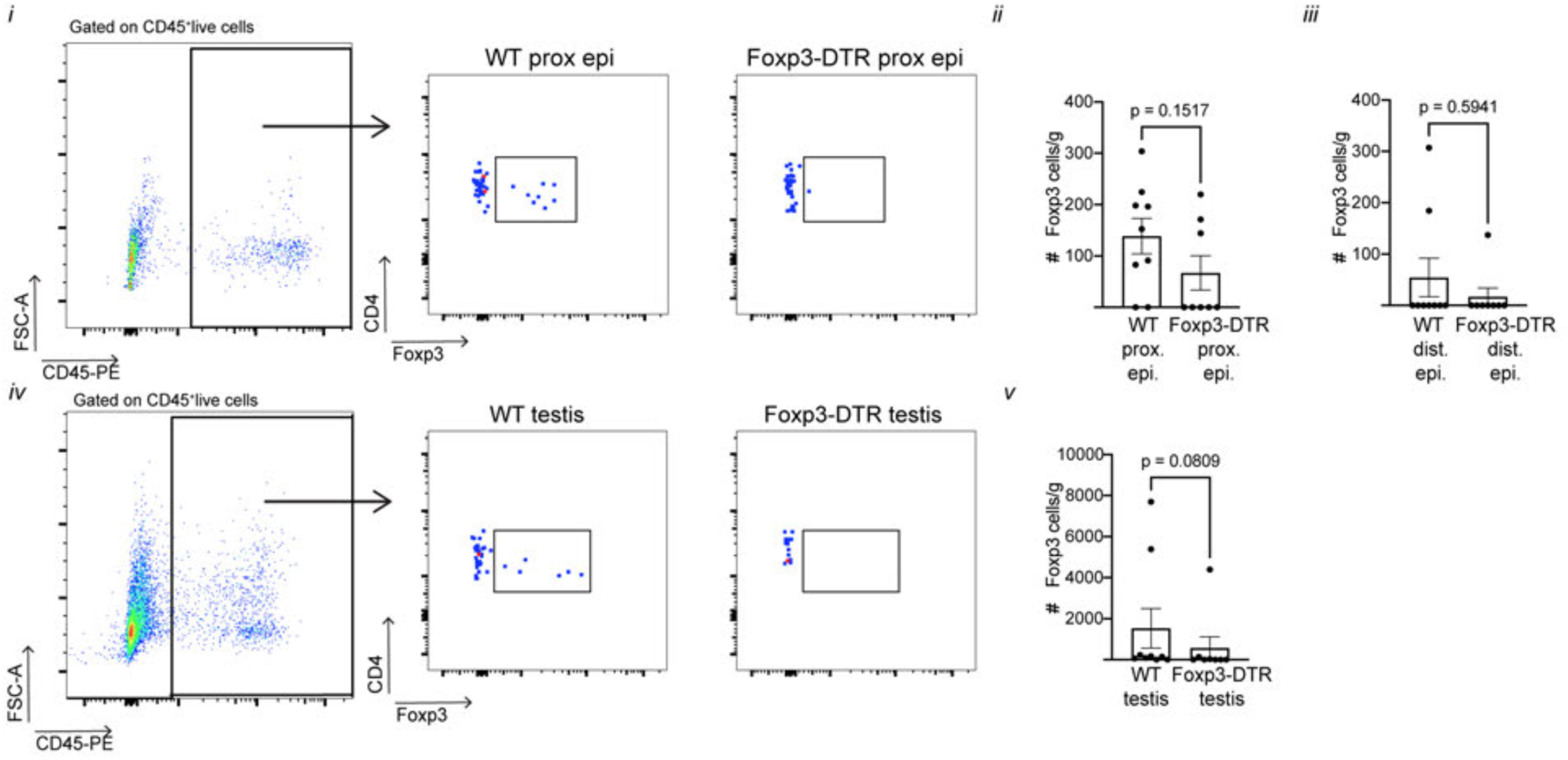
Circulating Treg population in the epididymis and testis 1 day after DT treatment. Flow cytometry gating strategy and analysis of circulating Tregs (CD45- PE^+^Foxp3^+^CD4^+^CD45-BV711^+^) per gram of proximal (*i-ii*) and distal epididymis (*iii*), and testis (*iv-v*). Each dot represents a pool of 2 proximal epididymides, 2 distal epididymides, or 2 testes from each mouse. Data were analyzed using the Mann-Whitney test. Data are shown as means ± SEM.

**Supplementary Figure 2.**
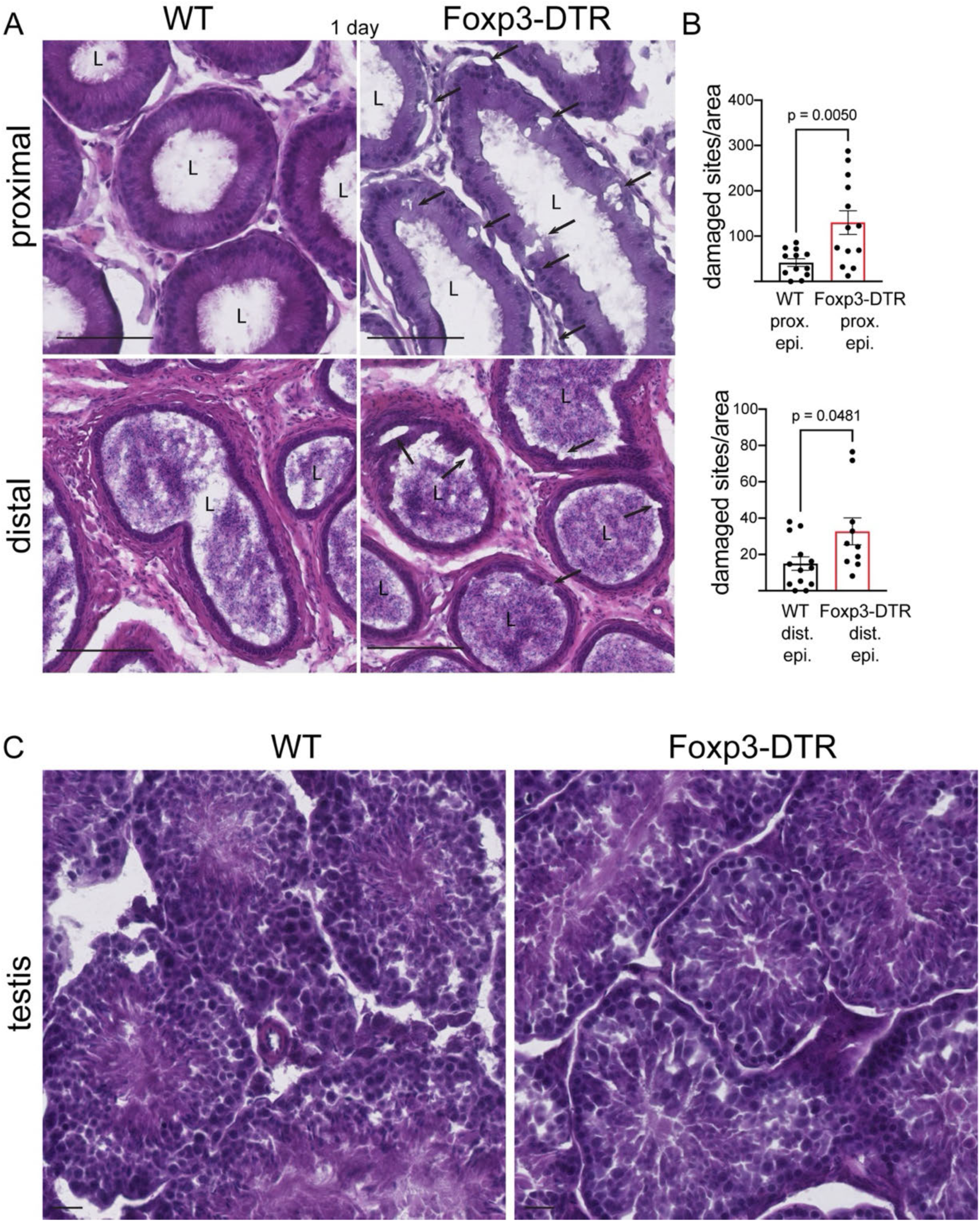
Histological evaluation of epididymis and testis of Foxp3-DTR mice 1 day after DT treatment. **A)** H&E staining of different epididymal regions: proximal (initial segments (IS) and caput) and distal (cauda) of DT-injected WT and Foxp3-DTR mice. **B)** Quantification of the number of damaged sites normalized per tissue area (nm^2^). Any morphological alteration in epididymal epithelia (arrows) was counted as a damaged site. Each area quantified is represented as a dot. Bars: 100 μm. L: Lumen. Data were analyzed using Student’s t-test (B, upper panel) or Mann-Whitney (B, lower panel) test. Data are shown as means ± SEM. **C)** H&E staining in the testis of DT-injected WT and Foxp3-DTR mice. A complete process of spermatogenesis was observed in both groups. Bars: 20 μm.

**Supplementary Figure 3.**
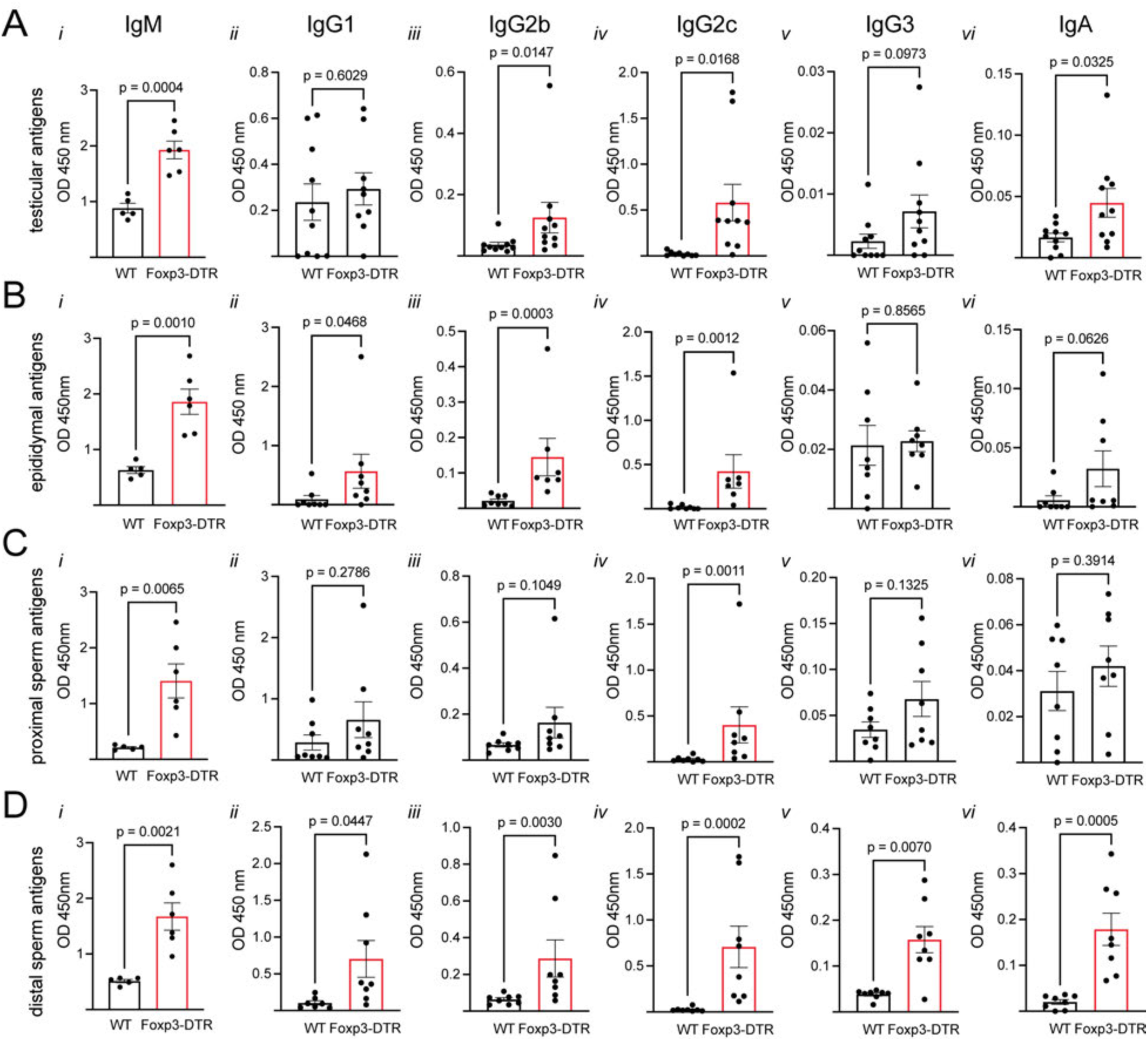
Serum autoantibody isotypification of Foxp3-DTR mice 2 weeks after DT treatment by ELISA. Serum IgM, IgG1, IgG2b, IgG2c, IgG3, and IgA levels of DT-injected Foxp3-DTR and WT mice against testicular antigens (**A**), epididymal antigens (**B**), proximal sperm antigens (**C**) and distal sperm antigens (**D**). Data were analyzed using Student’s t-test (A*i; iv; v*, B*i; v*, C*i; v; vi*, D*i; ii; vi*) or Mann-Whitney (A*ii; iii; vi*, B*ii; iii; iv; vi*, C*ii; iii; iv*, D*iii; iv; v*) test. Data are shown as means ± SEM.

**Supplementary Figure 4.**
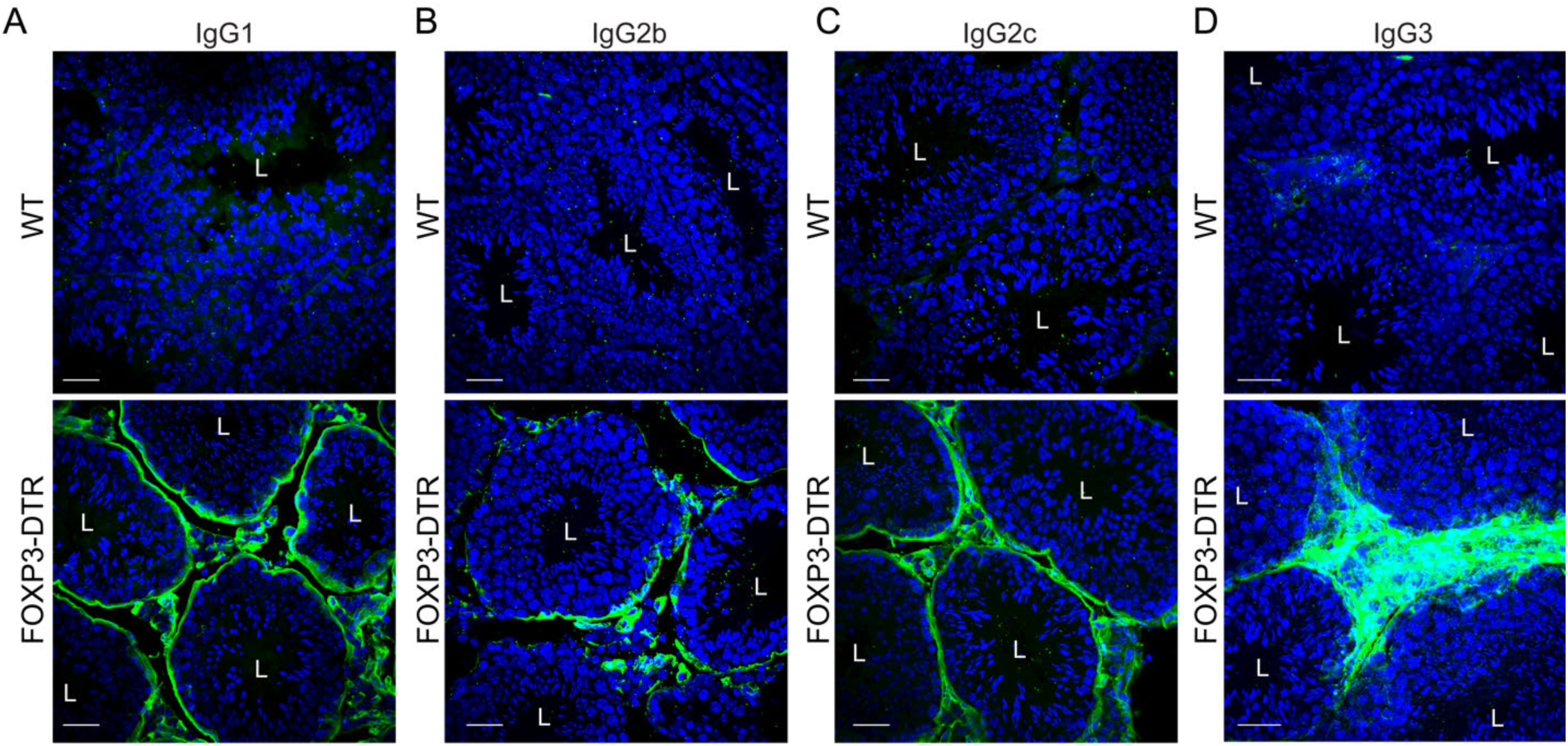
Evaluation of IgG1, IgG2b, IgG2c, and IgG3 on testis of Foxp3- DTR mice 2 weeks after Treg depletion by immunofluorescence. Confocal microscopy images showing immunolabeling of IgG1 (**A**), IgG2b (**B**), IgG2c (**C**), and IgG3 (**D**) on testis of WT and Foxp3-DTR mice 2 weeks after DT treatment. Nuclei are labeled with DAPI (blue). Bars: 20 μm. L: Lumen.

**Supplementary Figure 5.**
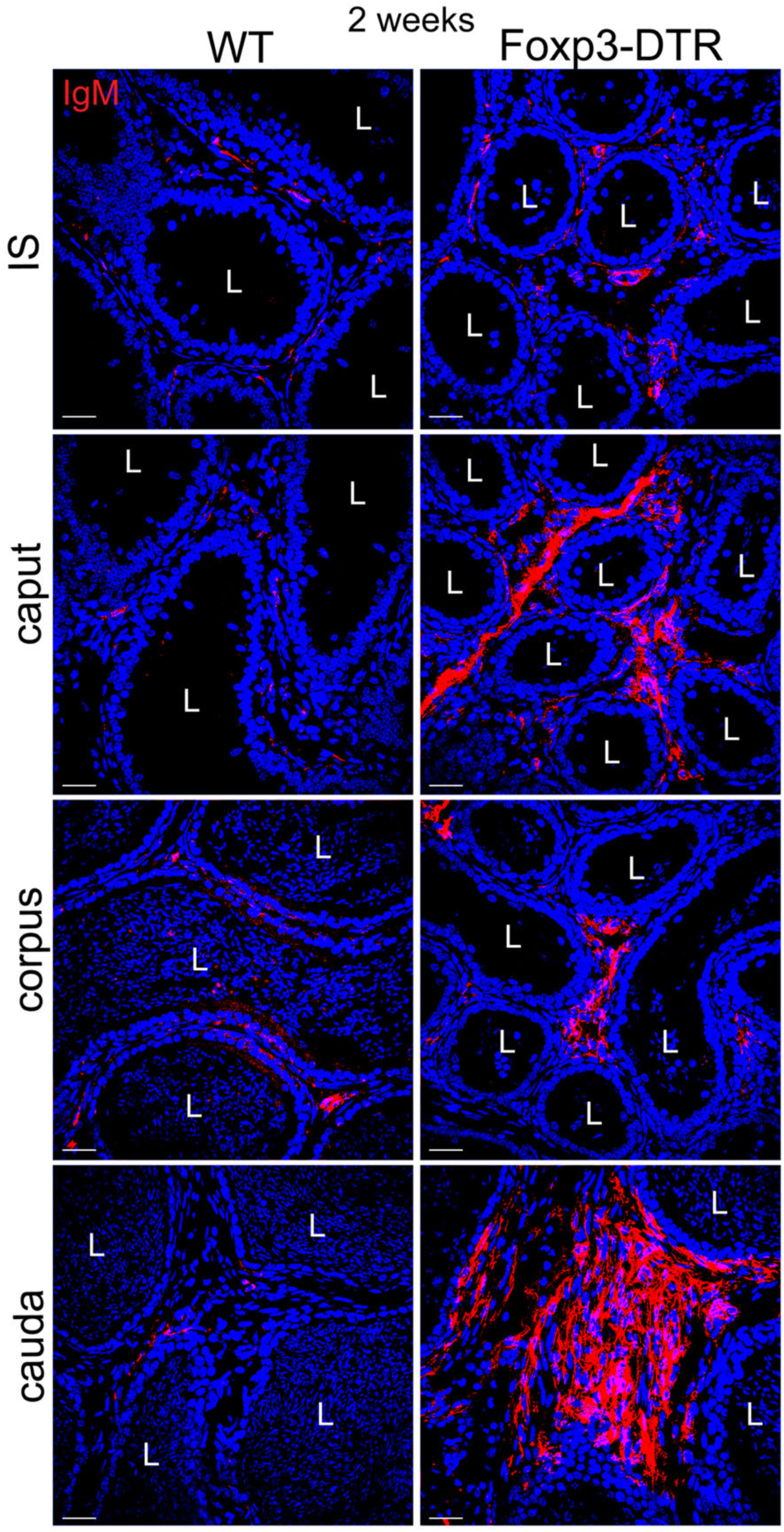
Evaluation of IgM on all epididymal regions of Foxp3-DTR mice 2 weeks after Treg depletion by immunofluorescence. Confocal microscopy images showing immunolabeling of IgM (red) on all epididymal regions of WT and Foxp3-DTR mice 2 weeks after DT treatment. Proximal epididymis comprises initial segments (IS) and caput; Distal epididymis comprises corpus and cauda. Nuclei are labeled with DAPI (blue). Bars: 20 μm. L: Lumen.

**Supplementary Figure 6.**
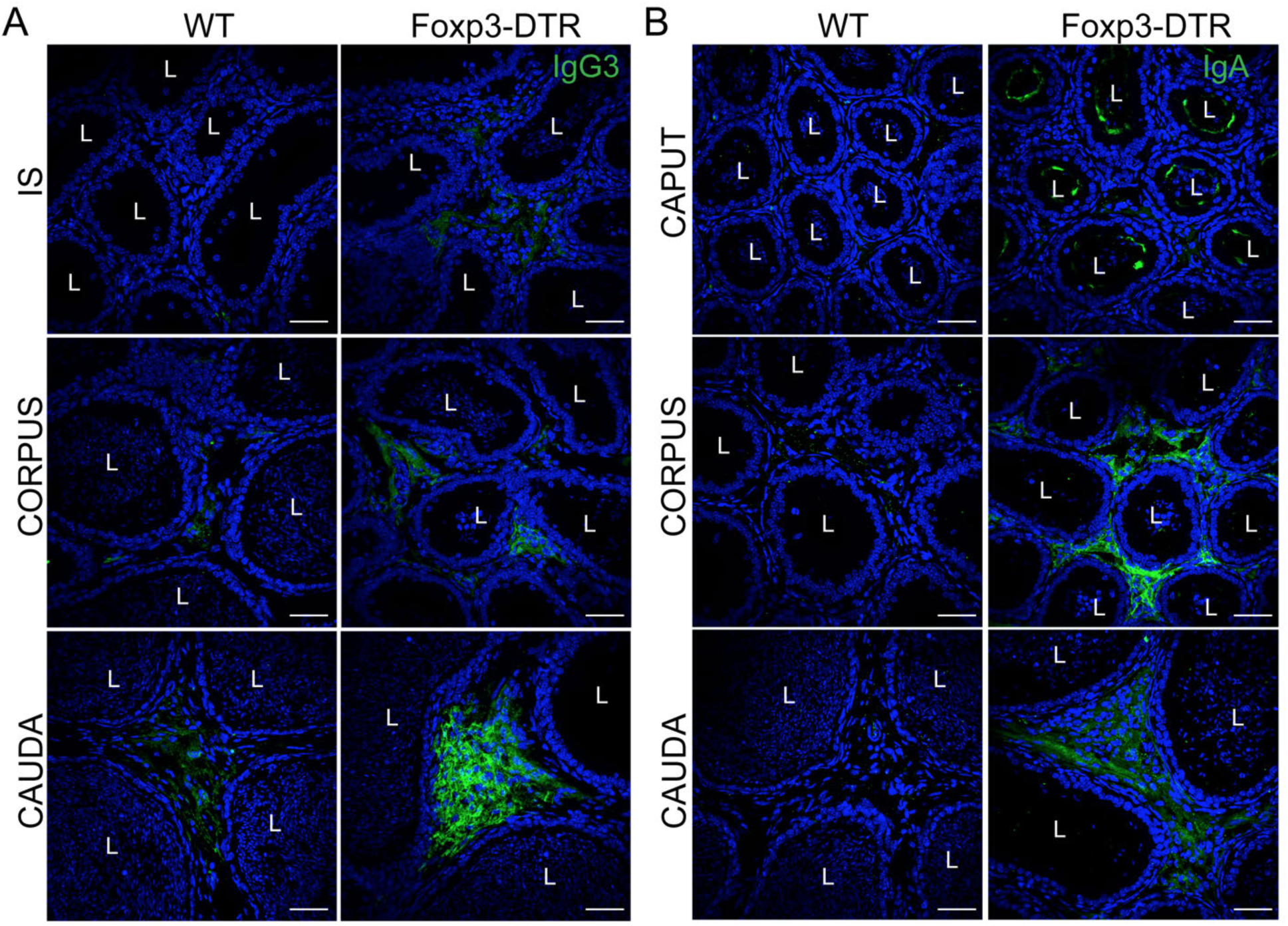
Evaluation of IgA and IgG3 on all epididymal regions of Foxp3- DTR mice 2 weeks after Treg depletion by immunofluorescence. Confocal microscopy images showing immunolabeling of IgG3 (**A**, green) and IgA (**B**, green) on all epididymal regions of WT and Foxp3-DTR mice 2 weeks after DT treatment. Nuclei are labeled with DAPI (blue). Bars: 20 μm. L: Lumen.

**Supplementary Figure 7.**
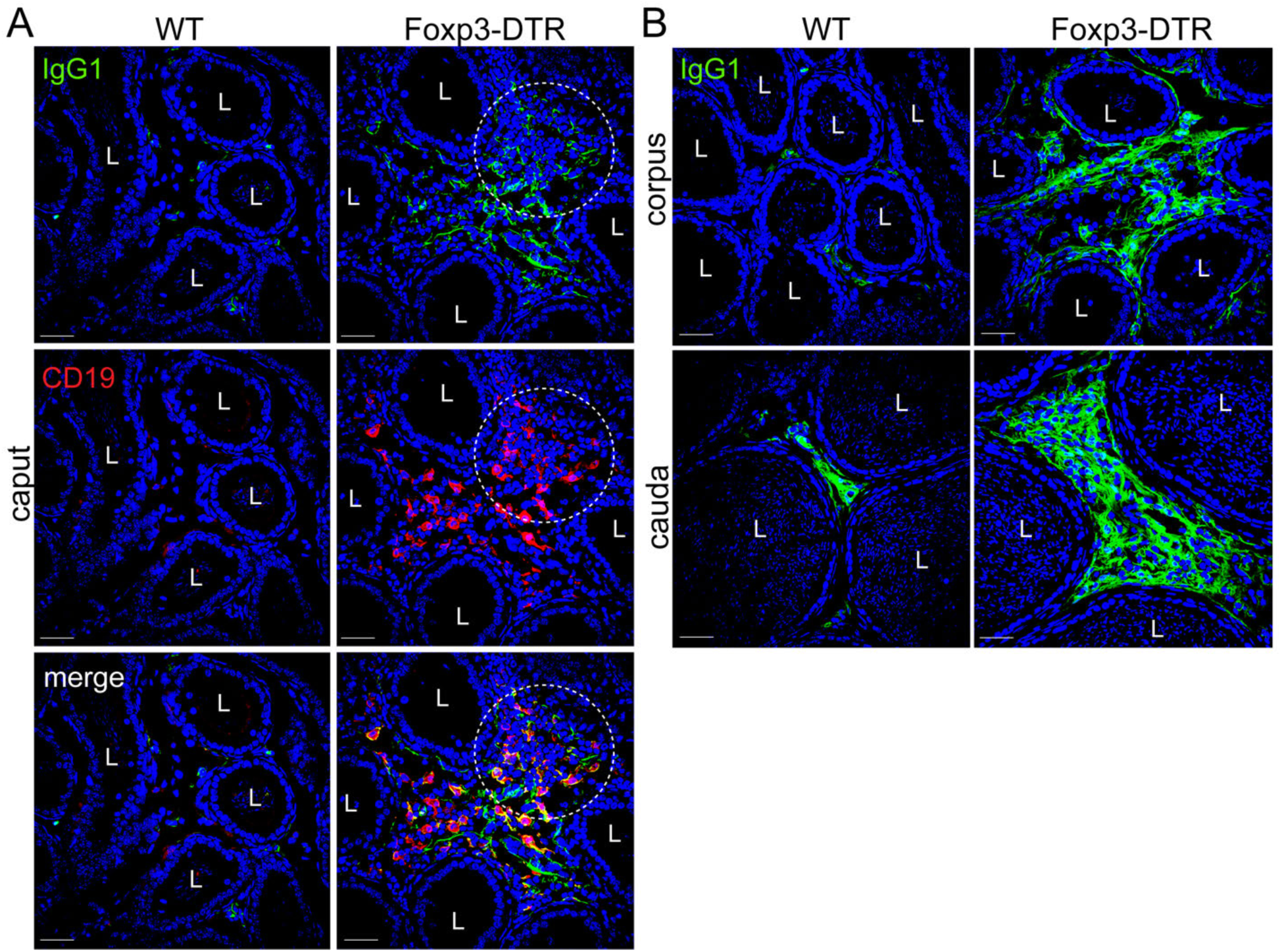
Evaluation of IgG1 on all epididymal regions of Foxp3-DTR mice 2 weeks after Treg depletion. **A)** Confocal microscopy images showing immunolabeling of IgG1 (green) and CD19 (red) on caput of WT and Foxp3-DTR mice 2 weeks after DT treatment. **B)** Confocal microscopy images showing immunolabeling of IgG1 (green) on corpus and cauda of WT and Foxp3-DTR mice 2 weeks after DT treatment. Nuclei are labeled with DAPI (blue). Bars: 20 μm. L: Lumen.\

**Supplementary Figure 8.**
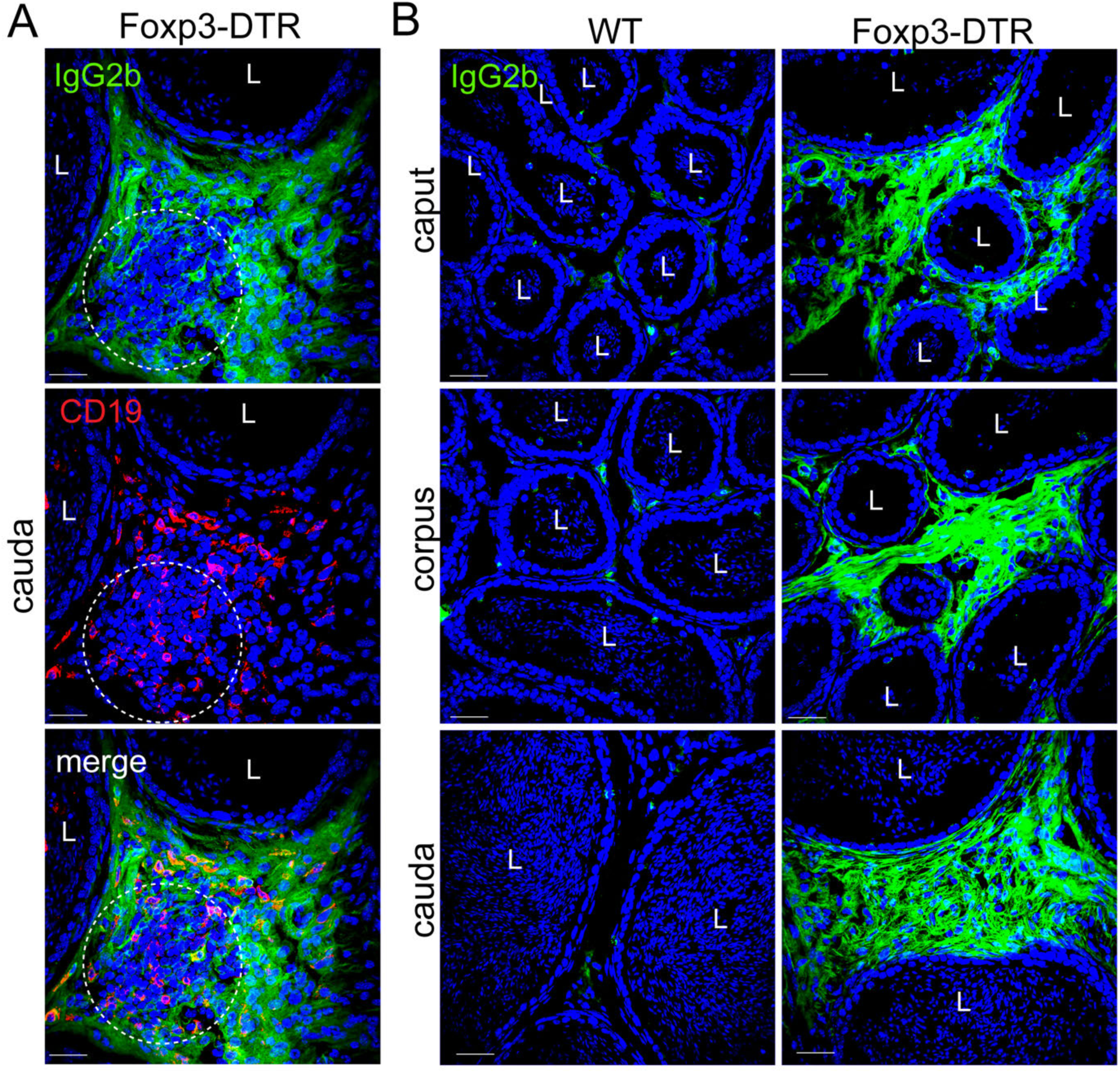
Evaluation of IgG2b on all epididymal regions of Foxp3-DTR mice 2 weeks after Treg depletion. **A)** Confocal microscopy images showing IgG2b (green) immunolabeling and CD19 (red) on the cauda region of WT and Foxp3-DTR mice 2 weeks after DT treatment. **B)** Confocal microscopy images showing IgG2b (green) immunolabeling on caput, corpus, and cauda of WT and Foxp3-DTR mice 2 weeks after DT treatment. Nuclei are labeled with DAPI (blue). Bars: 20 μm. L: Lumen.

**Supplementary Figure 9.**
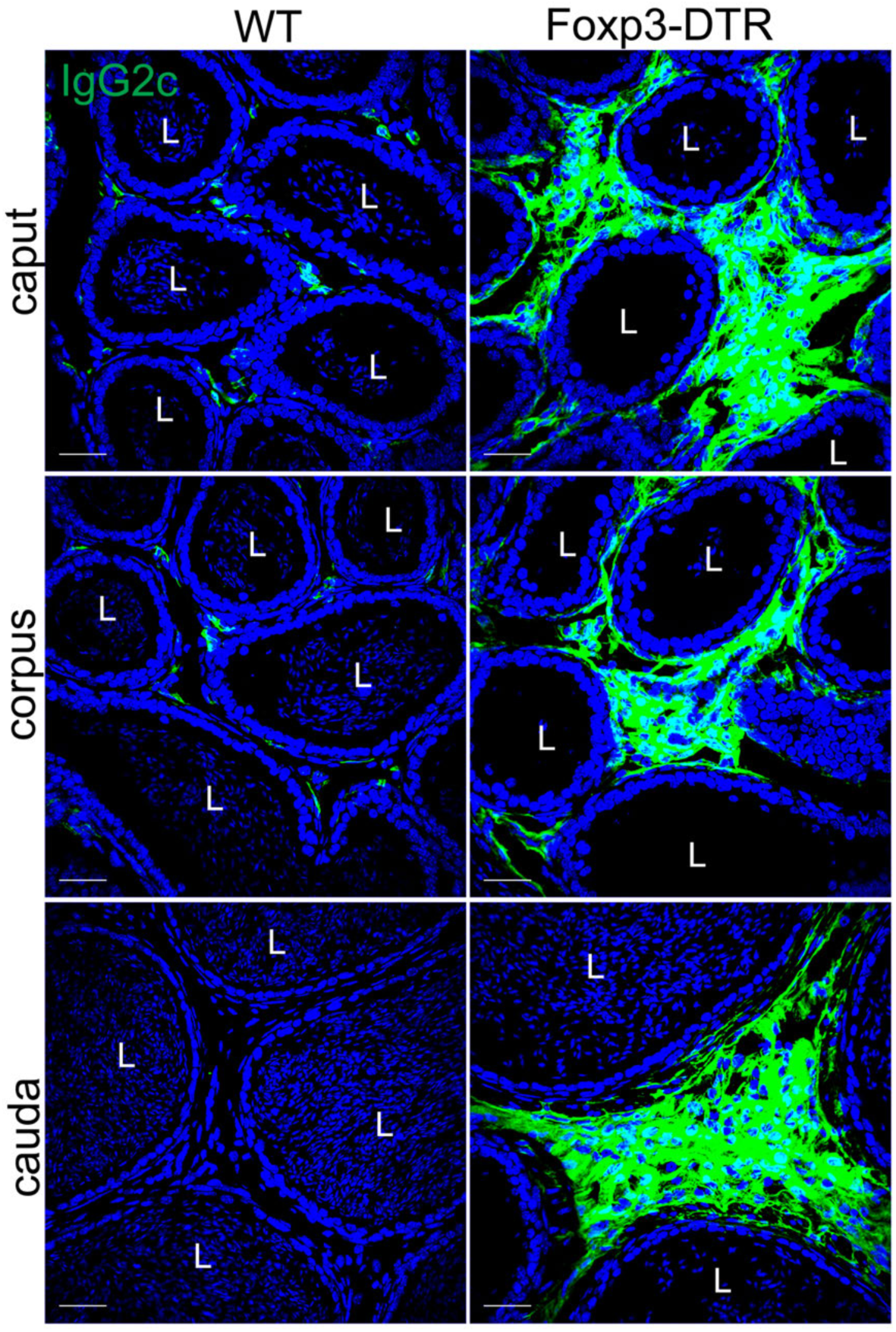
Evaluation of IgG2c on all epididymal regions of Foxp3-DTR mice 2 weeks after Treg depletion. Confocal microscopy images showing IgG2c (green) immunolabeling on epididymal regions of WT and Foxp3-DTR mice 2 weeks after DT treatment. Nuclei are labeled with DAPI (blue). Bars: 20 μm. L: Lumen.

**Supplementary Figure 10.**
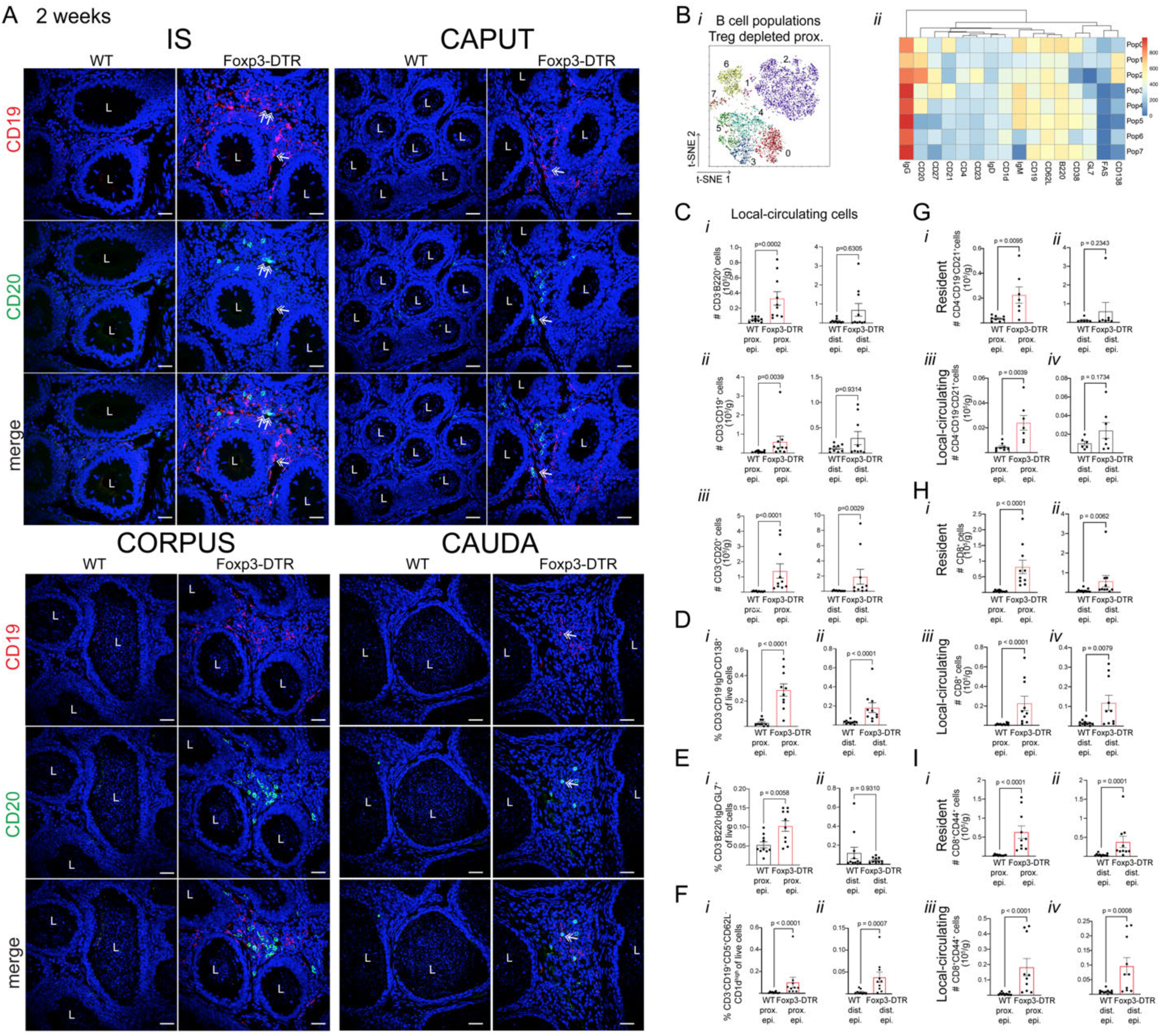
Localization and characterization of CD20^+^ and CD19^+^ B cells in the epididymis 2 weeks after Treg depletion. **A)** Immunolabeling of CD19 (red) and CD20 (green) B cells in the IS, caput, corpus, and cauda regions of DT-injected WT and Foxp3-DTR mice. Double-head arrows indicate double positive cells. **B)** tSNE and FlowSOM analysis of the B cell population of proximal epididymis from Foxp3-DTR mice (*i*) and its relative marker expression heatmap (*ii*). **C)** Absolute count (#) of local-circulating B cells (CD3^-^B220^+^ (*i*), CD3^-^ CD19^+^ (*ii*), and CD3^-^CD20^+^ (*iii*) cells) per gram of proximal and distal epididymis by flow cytometry. **D)** Relative abundance of local-circulating Plasma B cells (CD3⁻CD19⁺IgD⁻CD138⁺ of live cells) in the proximal and distal epididymides. **E)** Relative abundance of local-circulating GC B cells (CD3⁻B220⁺IgD⁻GL7⁺ of live cells) in the proximal and distal epididymides. **F)** Relative abundance of local-circulatory Bregs (CD3^-^CD19^+^CD5^+^CD62L^-^CD1d^hi^ of live cells) in the proximal and distal epididymides. **G)** Absolute count (#) of tissue-resident and local-circulating FDC (CD4^-^CD19^-^ CD21^+^ cells) in the proximal and distal epididymides. Absolute count (#) of tissue-resident and local-circulating CD8^+^ (CD4^-^CD8^+^) (**H**) and CD8^+^CD44^+^ T cells (**I**) per gram of proximal and distal epididymis. Nuclei are labeled with DAPI (blue). Bars: 20 μm. L: Lumen. Data were analyzed using Student’s t-test (E*i*, G*i*, *iii*, *iv*) or Mann-Whitney test (C*i-iii,* D, E*ii*, F, G*ii*, H*i-iv*, I*i-iv*). Data are shown as means ± SEM.

**Supplementary Figure 11.**
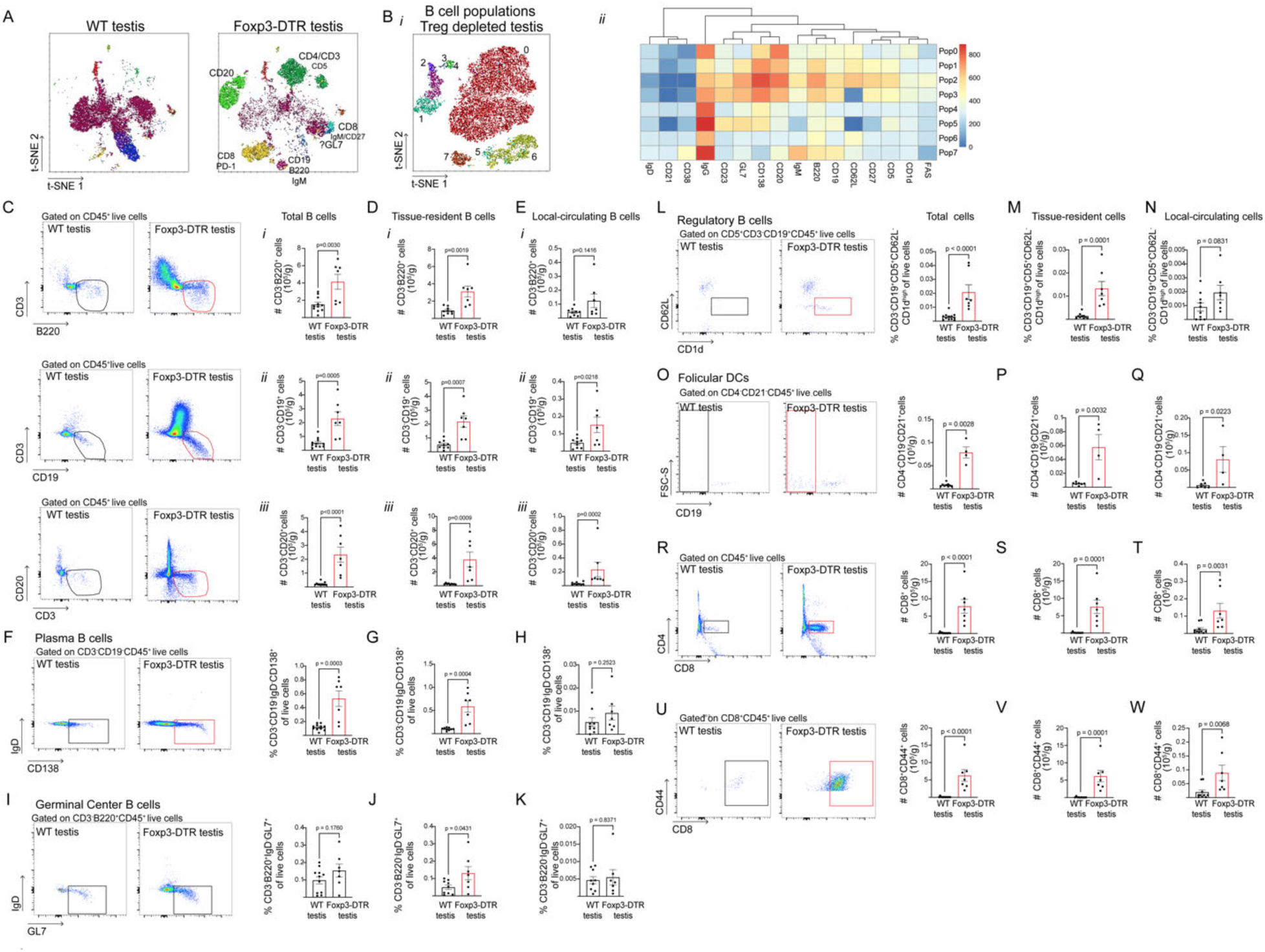
B cell populations analysis in the testis of Foxp3-DTR mice 2 weeks after Treg depletion by flow cytometry. tSNE and FlowSOM pan-populations analysis of testis from WT and Foxp3-DTR mice (**A**) and of the B cell populations of testis from Foxp3- DTR mice (B*i*) and its relative marker expression heatmap (B*ii*). Flow cytometry gating strategy and absolute count (#) of total (**C**), resident (**D**), and local circulating (**E**) B cells (CD3^-^B220^+^, CD3^-^ CD19^+^, and CD3^-^CD20^+^) per gram of testis. Flow cytometry gating strategy and relative abundance of total (**F**), resident (**G**), and local circulating (**H**) Plasma B cells (CD3⁻CD19⁺IgD⁻CD138⁺ of live cells) in the testes. Flow cytometry gating strategy and relative abundance of total (**I**), resident (**J**), and local circulating (**K**) GC B cells (CD3⁻B220⁺IgD⁻GL7⁺ of live cells) in the testes. Flow cytometry gating strategy and relative abundance of total (**L**), resident (**M**), and local circulating (**N**) Bregs (CD3-CD19+CD5+CD62L-CD1d^hi^ of live cells) in the testes. Flow cytometry gating strategy and absolute count (#) of total (**O**), resident (**P**), and local circulating (**Q**) FDC (CD4^-^CD19^-^CD21^+^ cells) in the testes. Flow cytometry gating strategy and absolute count (#) of total (**R**), resident (**S**), and local circulating (**T**) CD8^+^ T cells (CD4^-^CD8^+^) per gram of testis. Flow cytometry gating strategy and absolute count (#) of total (**U**), resident (**V**), and local circulating (**W**) activated CD8^+^ T cells (CD4^-^CD8^+^CD44^+^) per gram of testis. Data were analyzed using Student’s t-test (C*i*, *ii*, D*i-iii*, E*ii*, F, G, I, J, N, P, Q) or the Mann-Whitney test (C*iii*, E*i*, *iii*, H, K-M, O, R-W). Data are shown as means ± SEM.

**Supplementary Figure 12.**
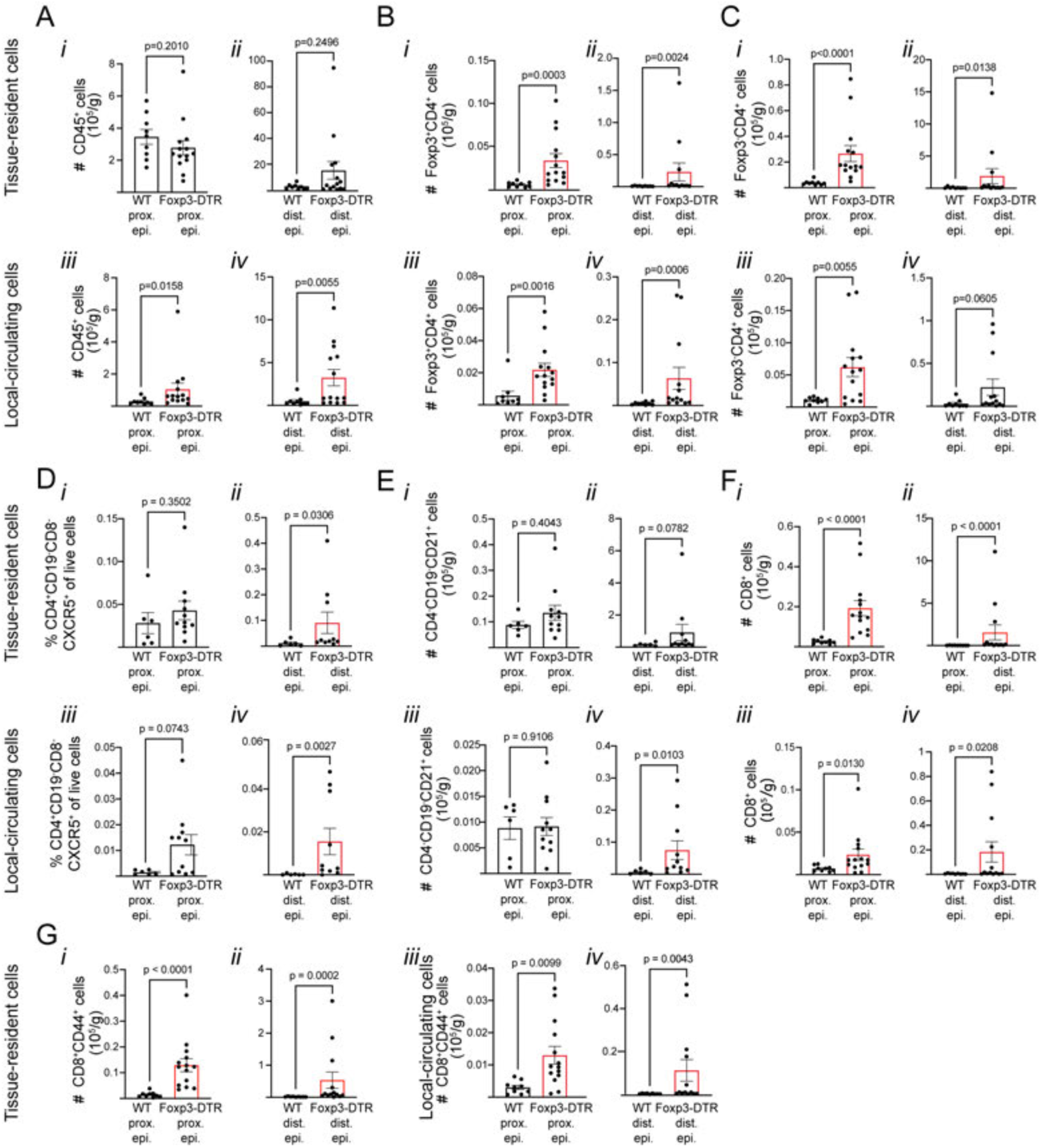
T and DC infiltration in the epididymis 8 weeks after Treg depletion by flow cytometry. **A)** Absolute count (#) of tissue-resident and local-circulatory of CD45^+^ cells per gram of proximal and distal epididymis. **B)** Absolute count (#) of tissue-resident and local-circulatory Tregs per gram of proximal and distal epididymides. **C)** Absolute count (#) of tissue-resident and local-circulatory CD4 T lymphocytes per gram of proximal and distal epididymis. **D)** Relative abundance of tissue-resident and local-circulatory T follicular helper cells (Tfh, CD4+CD8-CD19-CXCR5+ of live cells) in the proximal and distal epididymides. **E)** Absolute count (#) of tissue-resident and local-circulatory FDCs per gram of proximal and distal epididymis. Absolute count (#) of tissue-resident and local-circulatory CD8^+^ (**F**) and activated CD8^+^ (**G**) T lymphocytes per gram of proximal and distal epididymis. Data were analyzed using Student’s t- test (E*iii,* G*iii*) or Mann-Whitney test (A-D, E*i, ii, iv,* F, G*i-ii, iv*). Data are shown as means ± SEM.

**Supplementary Figure 13.**
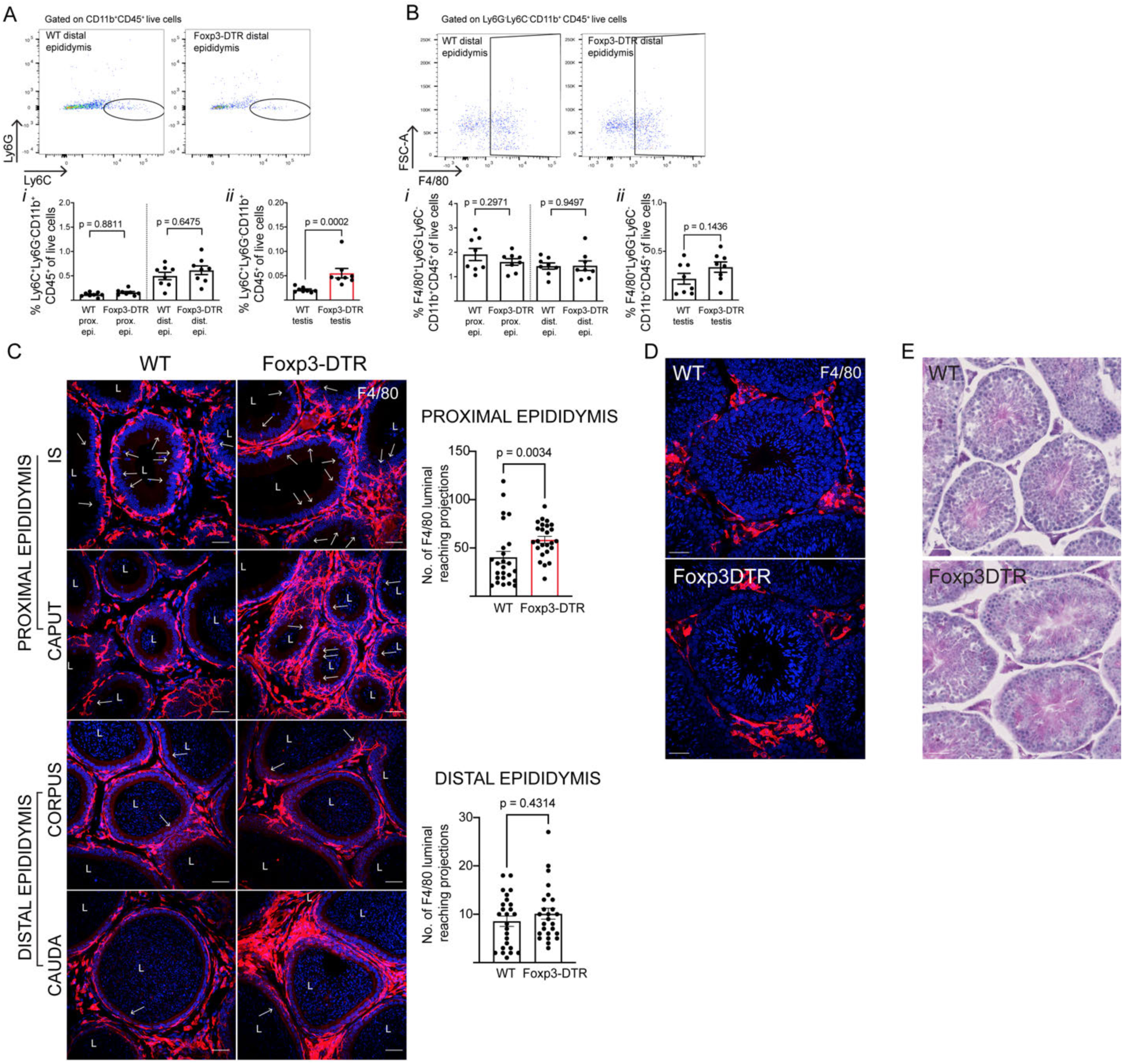
Myeloid cell analysis 8 weeks after Treg depletion. **A)** Flow cytometry gating strategy and relative abundance of monocyte (CD45^+^CD11b^+^Ly6G^-^Ly6C^+^ of live cells) in the proximal and distal epididymides. **B)** Flow cytometry gating strategy and relative abundance of macrophages (CD45^+^CD11b^+^Ly6G^-^Ly6C^-^F4/80^+^ of live cells) in the proximal and distal epididymides. **C)** Imaging and quantification of the number of F4/80 luminal-reaching projections (red, arrows) in the proximal (IS and caput) and distal (corpus and cauda) epididymis of DT-injected WT and Foxp3-DTR mice per area of tissue (110,540 µm^2^). Each image quantification is represented as a dot. **D)** Immunolabeling of F4/80^+^ cells (red) on testis from WT and Foxp3-DTR mice. **E)** H&E staining of WT and Foxp3-DTR testis. A complete process of spermatogenesis was observed in both groups. Nuclei are labeled with DAPI (blue). Bars: 20 μm. L: Lumen. Data were analyzed using Student’s t-test (A*i*, B*i-ii)* or Mann-Whitney test (A*ii*, C). Data are shown as means ± SEM.

**Supplementary Figure 14.**
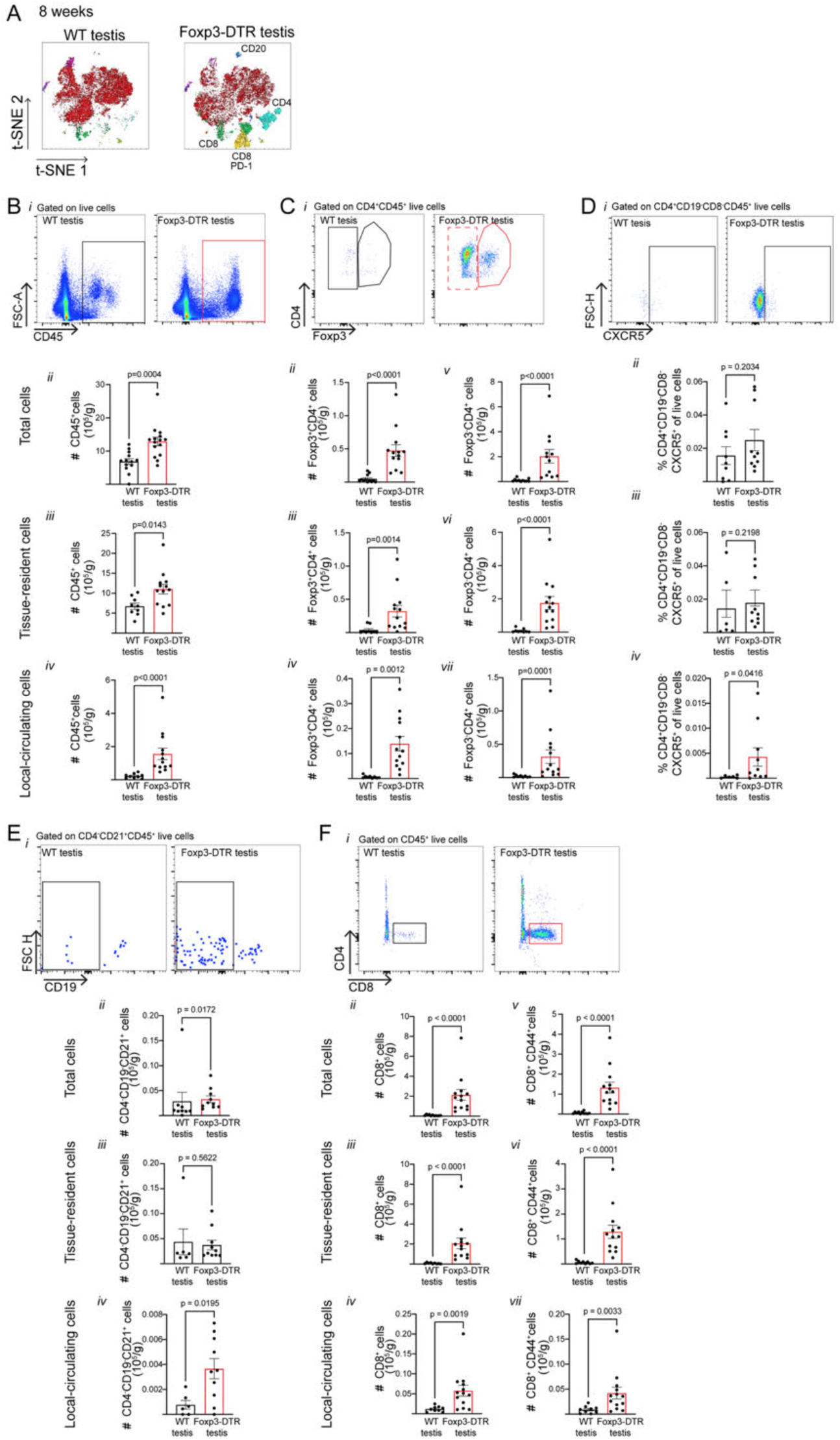
T cell and DC infiltration in the testis of DT-injected Foxp3-DTR mice 8 weeks after Treg ablation by flow cytometry. **A)** tSNE and FlowSOM pan-populations analysis of DT-treated WT and Foxp3-DTR testes. **B)** Flow cytometry gating strategy (*i*) and absolute count (#) of total *(ii*), resident (*iii*), and local circulating (*iv*) CD45^+^ cells per gram of testis. **C)** Flow cytometry gating strategy (*i*) and absolute count (#) of total (*ii, v*), tissue-resident (*iii, vi*), and local-circulatory (*iv, vii*) Treg and CD4 T lymphocytes per gram of testis. **D)** Flow cytometry gating strategy (*i*) and relative abundance of total (*ii*), tissue-resident (*iii*), and local-circulatory (*iv*) Tfh in the testes. **E)** Flow cytometry gating strategy (*i*) and absolute count (#) of total (*ii*), tissue-resident (*iii*), and local-circulatory (*iv*) FDCs cells per gram of testis. **F)** Flow cytometry gating strategy (*i*) and absolute count (#) of total (*ii, v*), tissue-resident (*iii, vi*), and local-circulatory (*iv, vii*) CD8 and activated CD8 T lymphocytes cells per gram of testis. Data were analyzed using Student’s t-test (B*iii*, C*iv*, E*iv*), or Mann-Whitney test (B*ii*, *iv*, C*ii*, *iii*, *v-vii*, D*ii-iv*, E*ii*, *iii*, F*ii-vii*). Data are shown as means ± SEM.

**Supplementary Figure 15.**
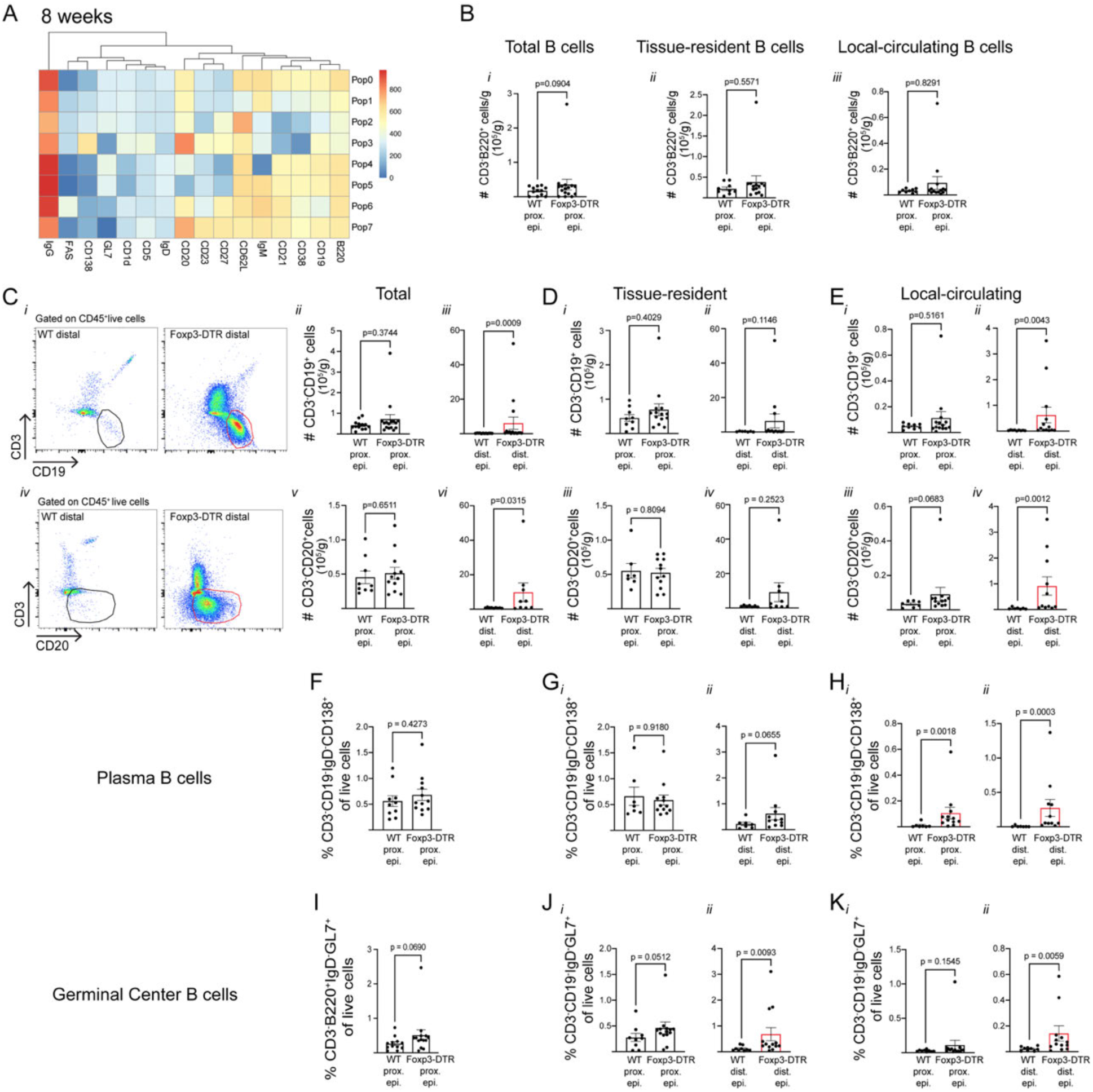
B cell analysis in the epididymis of Foxp3-DTR mice 8 weeks after Treg depletion by flow cytometry. **A)** Relative marker expression heatmap of tSNE and FlowSOM analysis of 8-week Treg depleted distal epididymis. **B)** Absolute count (#) of total (*i*), tissue-resident (*ii*), and local-circulating (*iii*) B cells per gram of proximal epididymis. **C)** Flow cytometry gating strategy and absolute count (#) of total B cells (CD3^-^CD19^+^ and CD3^-^CD20^+^ cells) per gram of proximal and distal epididymis. Absolute count (#) of tissue-resident (**D**) and local-circulating (**E**) B cells per gram of proximal and distal epididymis. Relative abundance of total (**F),** tissue-resident (**G**), and local-circulating (**H**) Plasma B cells in the proximal epididymides. Relative abundance of total (**I**), tissue-resident (**J**), and local-circulating (**K**) GC B cells in the proximal epididymides. Data were analyzed using Student’s t-test (D*iii*), or Mann-Whitney test (B*i- iii*, C*ii-iii, v-vi*, D*i-ii, iv*, E*i-iv*, F, G*i-ii*, H*i-ii*, I, J*i-ii*, K*i-ii*). Data are shown as means ± SEM.

**Supplementary Figure 16.**
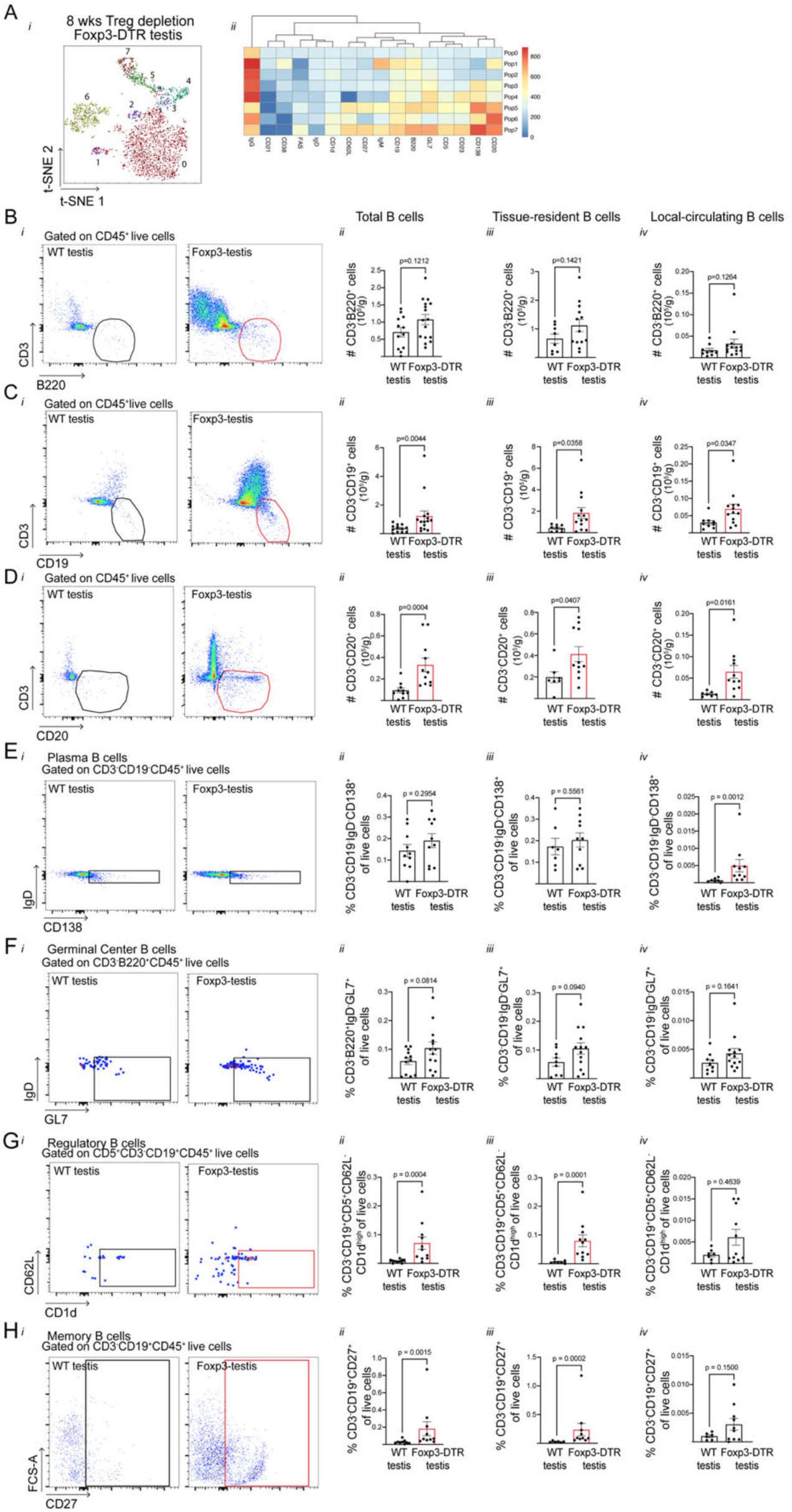
B cell analysis in the testis of Foxp3-DTR mice 8 weeks after Treg depletion by flow cytometry. **A)** tSNE and FlowSOM B cell populations analysis of testes from Foxp3-DTR mice (*i*) and relative marker expression heatmap (*ii*). Flow cytometry gating strategy (*i*) and absolute count (#) of total (*ii*), tissue-resident (*iii*) and local-circulating (*iv*) B220^+^ (**B**), CD19^+^ (**C**) and CD20^+^ (**D**) B cells per gram of testis from DT-treated WT and Foxp3-DTR mice. **E)** Flow cytometry gating strategy (*i*) and relative abundance of total (*ii*), tissue-resident (*iii*), and local-circulating (*iv*) plasma B cells in the testes. **F)** Flow cytometry gating strategy (*i*) and relative abundance of total (*ii*), tissue-resident (*iii*), and local-circulating (*iv*) GC B cells in the testes. **G)** Flow cytometry gating strategy (*i*) and relative abundance of total (*ii*), tissue-resident (*iii*) and local-circulating (*iv*) Bregs in the testes. **H)** Flow cytometry gating strategy (*i*) and relative abundance of total (*ii*), tissue-resident (*iii*), and local-circulating (*iv*) memory B cells in the testes. Data were analyzed using Student’s t-test (B*iii*, C*iii-iv,* D*iii-iv*, E*ii-iii*, F*ii-iii*, H*iv*), or Mann-Whitney test (B*ii-iv*, C*ii*, D*ii*, E*iv*, F*iv*, G*ii-iv*, H*ii-iii*). Data are shown as means ± SEM.

**Supplementary Figure 17.**
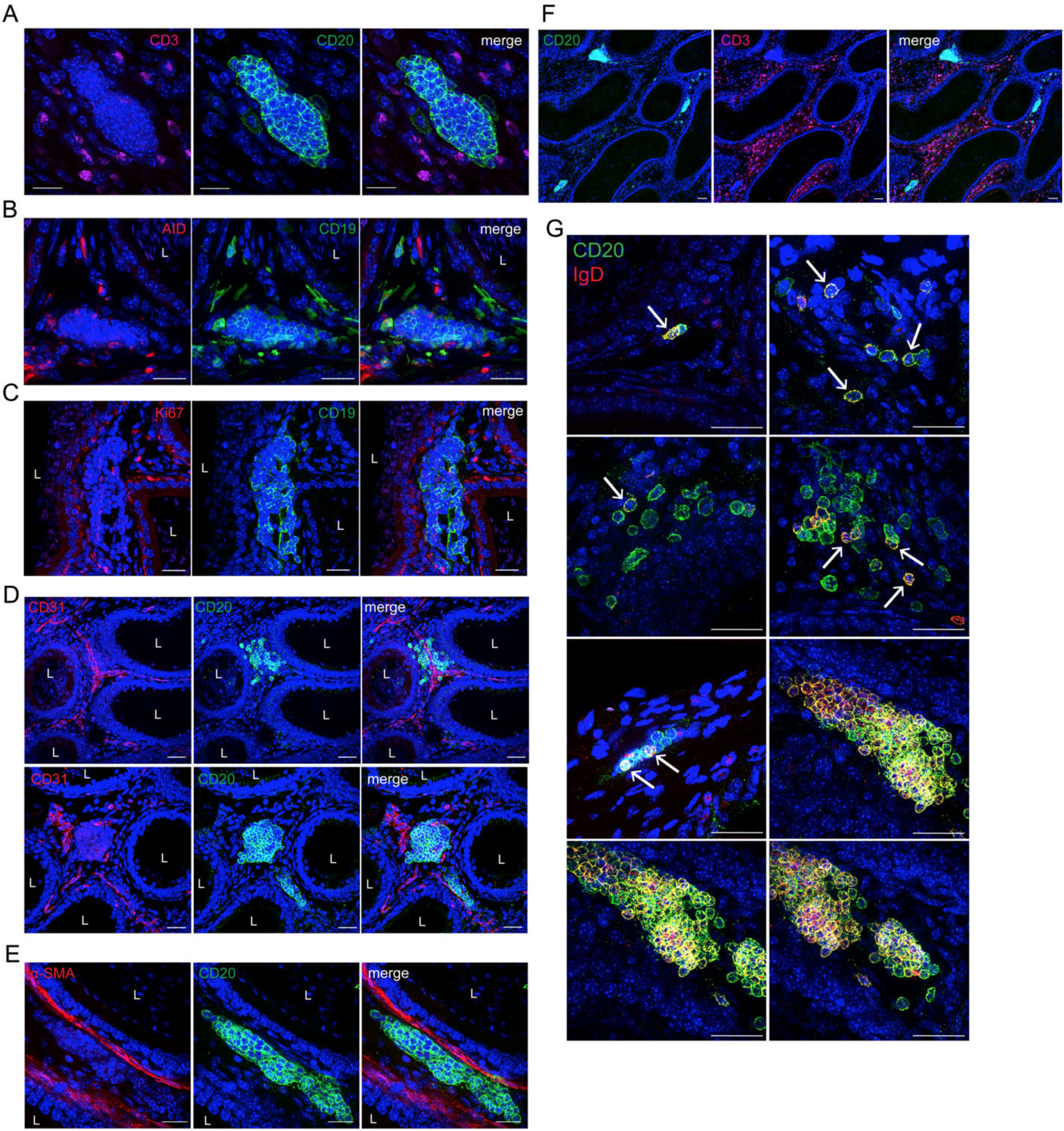
TLS characterization in the distal epididymides of 8 weeks DT-injected Foxp3-DTR mice. **A)** Immunolabeling of CD3 (magenta) and CD20 (green) in the epididymal distal region of DT-injected Foxp3-DTR mice. **B)** Immunolabeling of AID (red) and CD19 (green) in the epididymal distal region of DT-injected Foxp3-DTR mice. **C)** Immunolabeling of Ki-67 (red) and CD19 (green) in the epididymal distal region of DT-injected Foxp3-DTR mice. **C)** Immunolabeling of CD31 (red) and CD20 (green) in the epididymal distal region of DT-injected Foxp3-DTR mice. **E)** Immunolabeling of α-SMA (red) and CD20 (green) in the distal region of DT- injected Foxp3-DTR mice. **F)** Immunolabeling of CD3 (magenta) and CD20 (green) in the epididymal distal region of DT-injected Foxp3-DTR mice, low magnification. **G)** Immunolabeling of IgD (red) and CD20 (green) of different TLS development stages in the epididymal distal region of DT-injected Foxp3-DTR mice. Arrows indicate double positive cells. Please note that the three last pictures are different views of the same TLS shown in Fig. 6H. Nuclei are labeled with DAPI (blue). Bars: 20 μm.

**Supplementary Figure 18.**
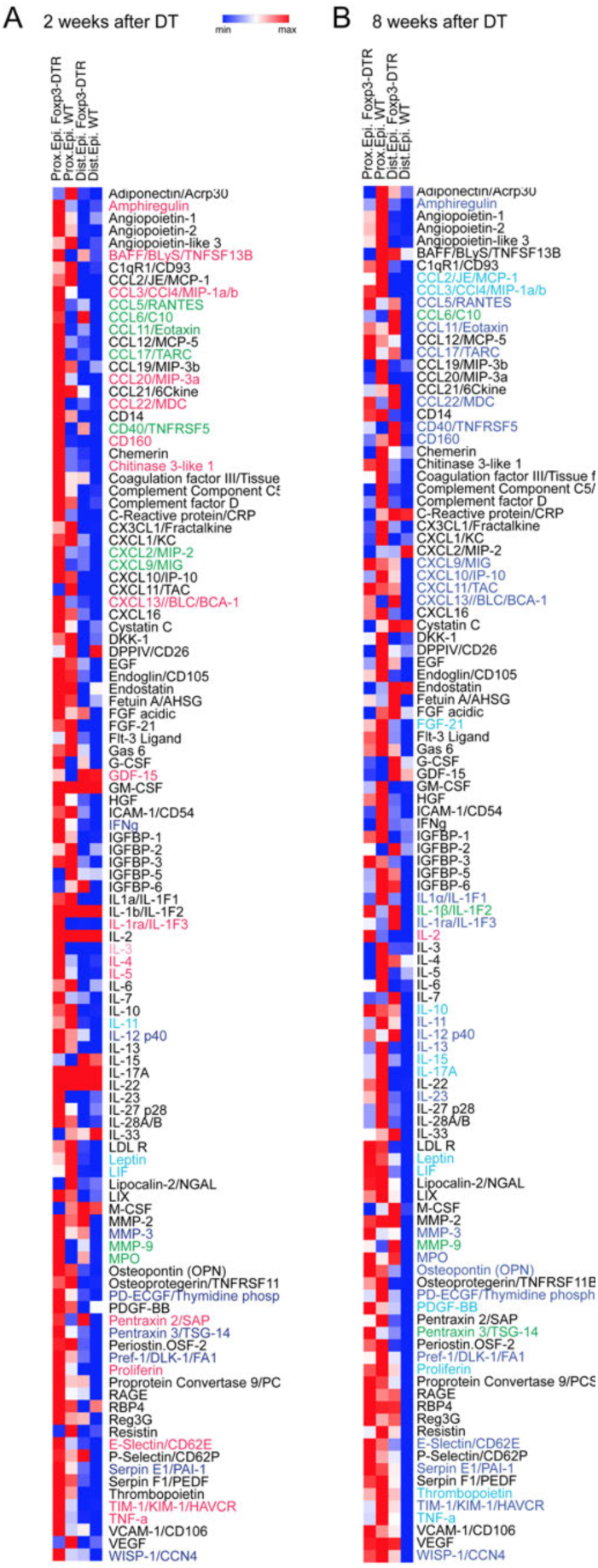
Soluble mediators within the proximal and distal epididymis at 2 and 8 weeks after DT treatment. Heatmap of the soluble mediator’s measurement by Proteome Profiler Mouse XL Cytokine Array (R&D) in proximal and distal epididymis tissues at 2 (**A**) and 8 (**B**) weeks after DT treatment. Pink: soluble mediators with a fold increase>2 in proximal epididymis. Blue: soluble mediators with a fold increase>2 in distal epididymis. Green: soluble mediators with a fold increase>2 in proximal and distal epididymis. Light pink: soluble mediators only present in Foxp3-DTR proximal epididymis. Light blue: soluble mediators only present in Foxp3-DTR distal epididymis.

**Supplementary Figure 19.**
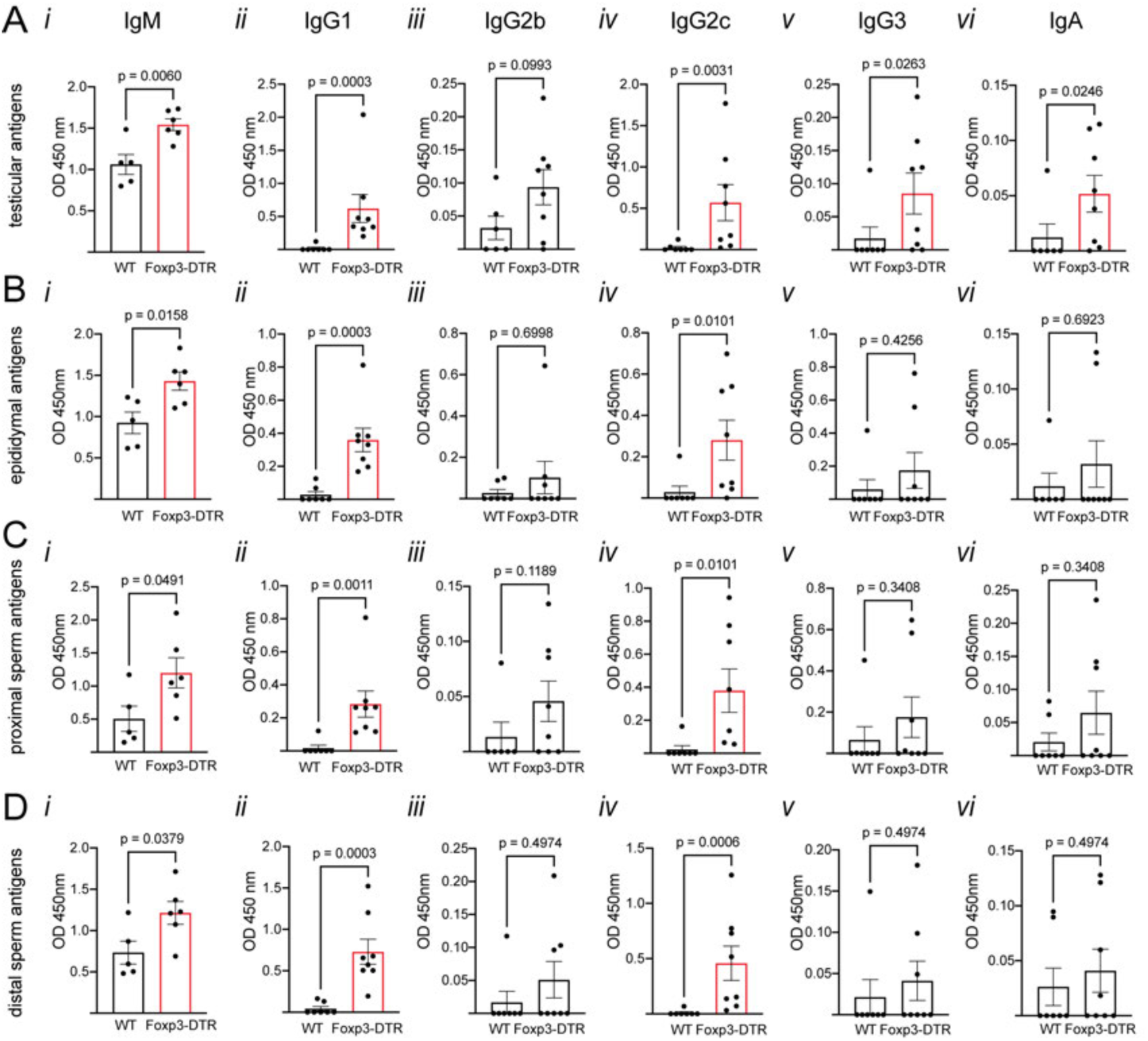
Serum autoantibody isotypification of Foxp3-DTR mice 8 weeks after DT treatment by ELISA. Autoantibody isotypification (IgM, IgG1, IgG2b, IgG2c, IgG3, and IgA) of DT-injected Foxp3-DTR and WT mouse serum against testicular **(A),** epididymal **(B)**, proximal sperm **(C)**, and distal sperm **(D)** antigens. Data were analyzed using Student’s t-test (A*i, iii*; B*i*; C*i*; D*i*) or Mann-Whitney test (A*ii, iv, v, vi*; B*ii, iii, iv, v, vi*; C*ii, iii, iv, v, vi*; D*ii, iii, v, vi*). Data are shown as means ± SEM.

**Supplementary Figure 20.**
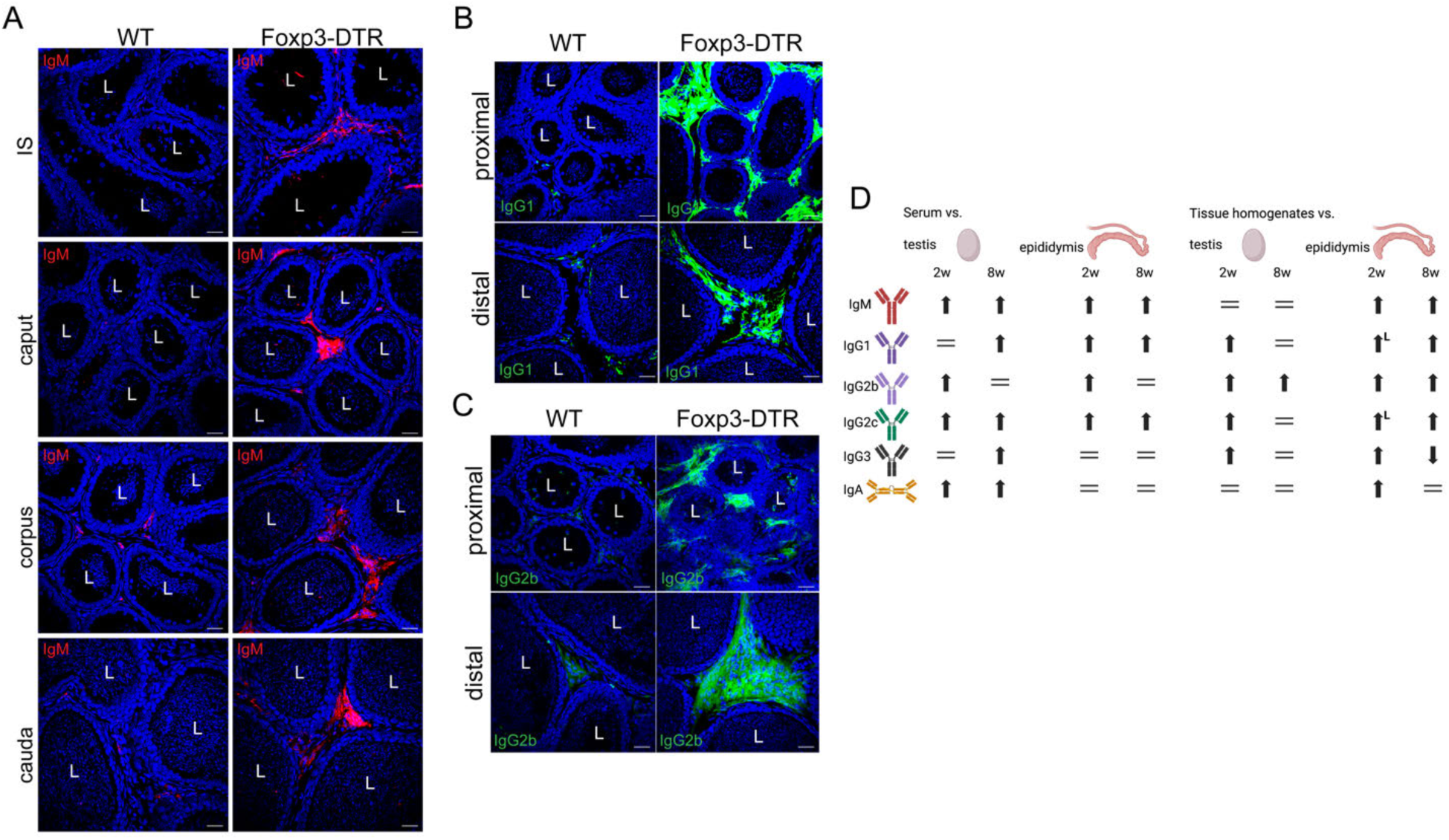
Autoantibody deposition in the epididymis 8 weeks after Treg depletion. **A)** Confocal microscopy images showing immunolabeling of IgM (red) on all epididymal regions of WT and Foxp3-DTR mice 8 weeks after DT treatment. **B)** Confocal microscopy images showing IgG1 (green) immunolabeling on proximal and distal epididymis of WT and Foxp3-DTR mice 8 weeks after DT treatment. **C)** Confocal microscopy images showing IgG2b (green) immunolabeling on proximal and distal epididymis of WT and Foxp3-DTR mice 8 weeks after DT treatment. Nuclei are labeled with DAPI (blue). Bars: 20 μm. L: Lumen. **D)** Summary of serum and tissue autoantibody profiles against epididymal and testicular antigens 2 and 8 weeks after DT treatment. L indicates the detection of an isotype in the epididymal lumen.

**Supplementary Figure 21.**
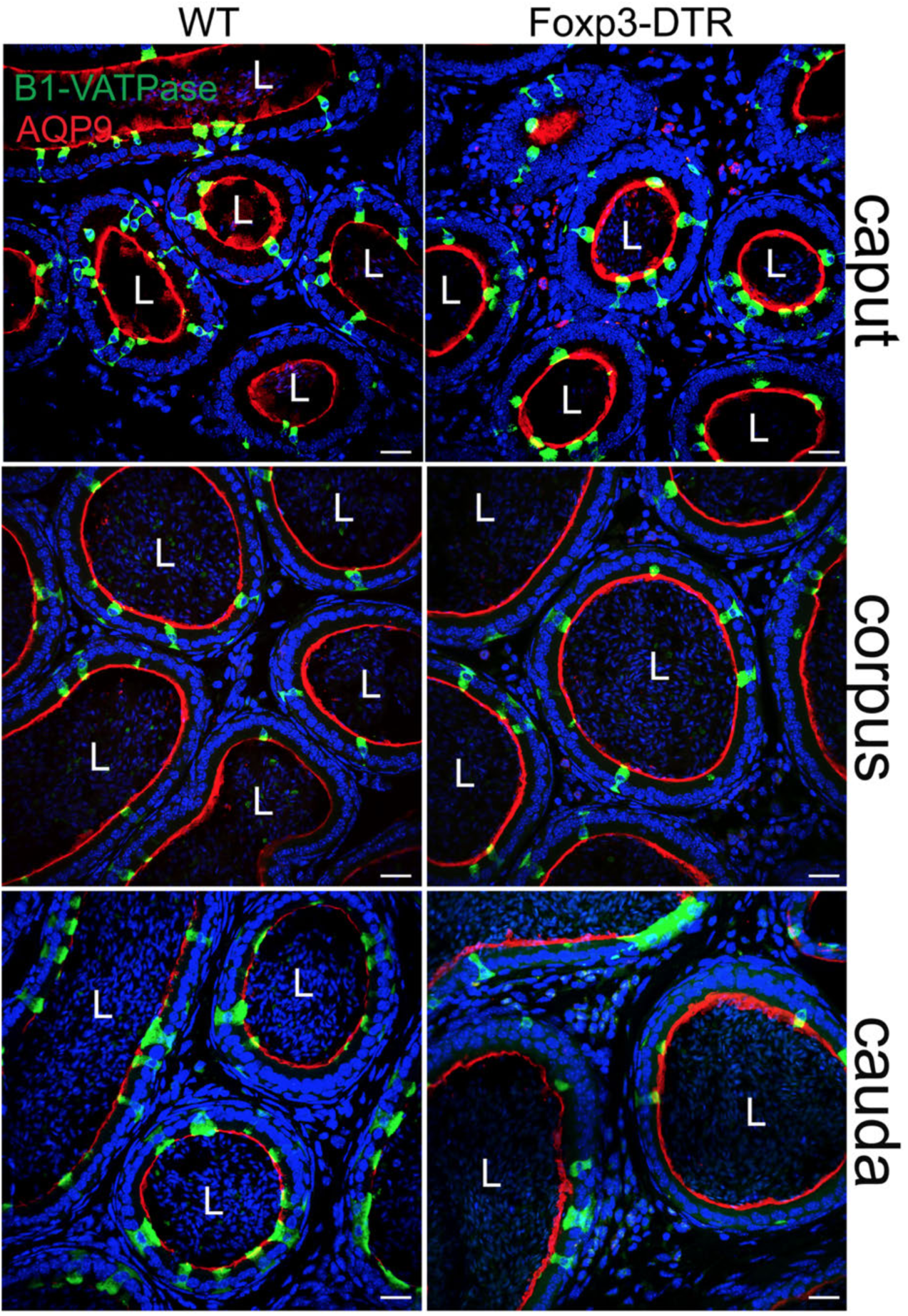
Epithelial integrity of the epididymal epithelia 8 weeks after Treg depletion. Confocal microscopy images showing B1 V-ATPase^+^ cells (a marker of clear cells (CCs); green) and AQP9^+^ cells (a marker of principal cells; red) in all epididymal regions of WT and Foxp3-DTR mice 8 weeks after DT injection. Nuclei are labeled with DAPI (blue). Bars: 20μm.

**Supplementary Table III.**
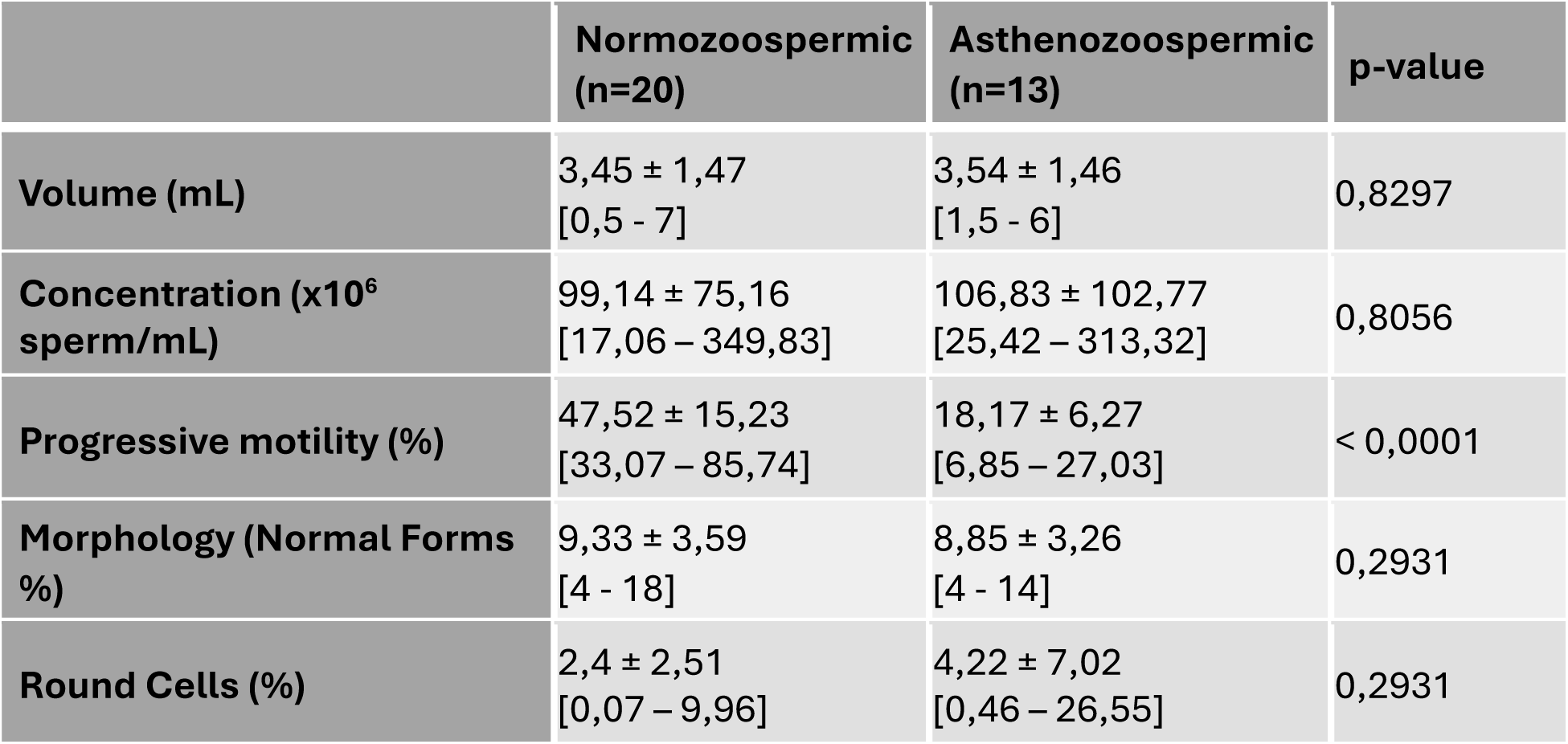
Descriptive parameters of the semen samples included in the study were divided into normozoospermic (n=20) and pathologic (n=13) groups. Results are indicated as mean ± standard deviation (SD), and [minimum and maximum] values. Student’s t-test was used to compare the groups.

## Auxiliary Material

**Supplementary Table I.** Proteomics analysis of sperm proteins recognized by serum ASAs of WT and Foxp3-DTR mice after 2 weeks of DT treatment.

**Supplementary Table II.** Proteomics analysis of sperm proteins recognized by the proximal and distal epididymal fluid ASAs of WT and Foxp3-DTR mice after 2 weeks of DT treatment.

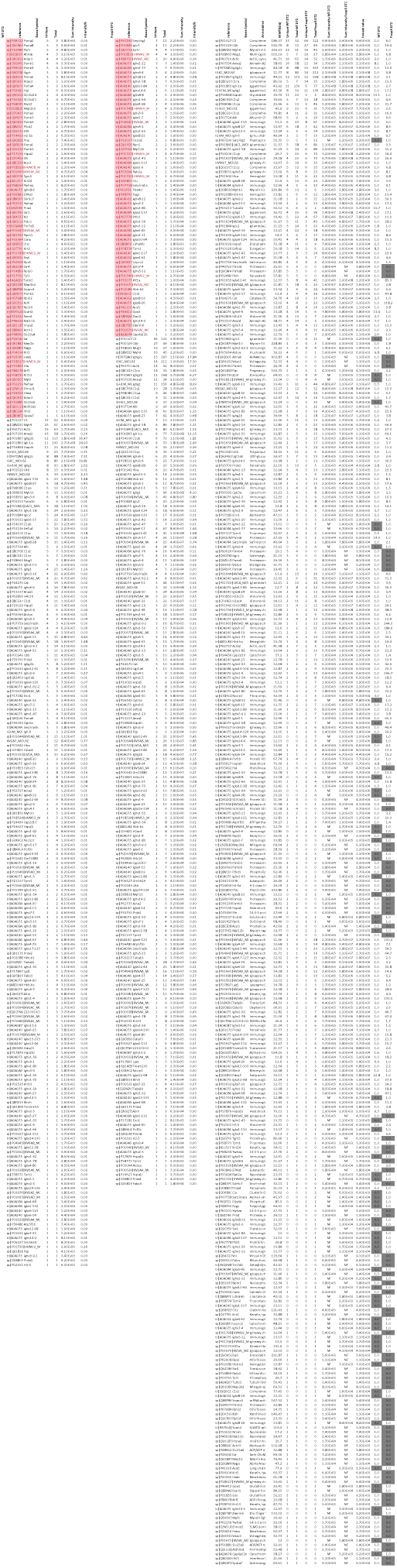
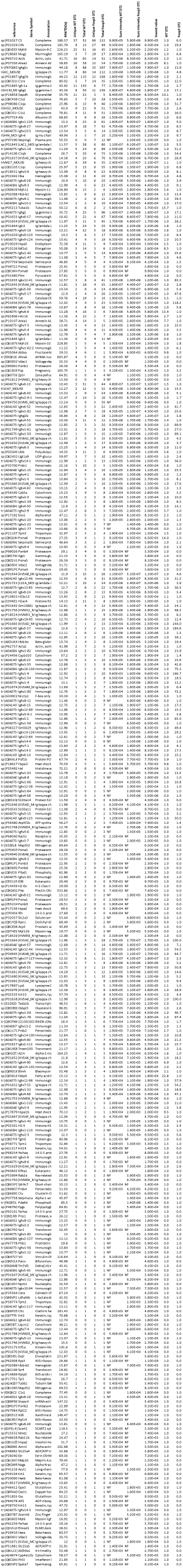
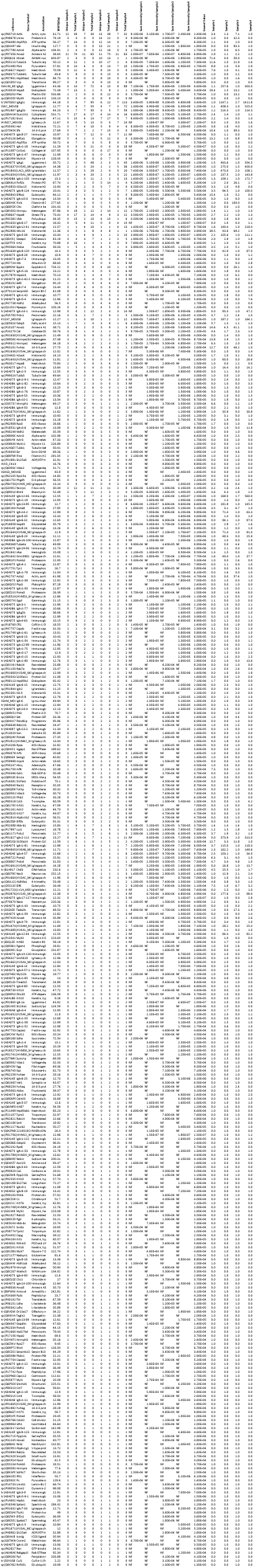
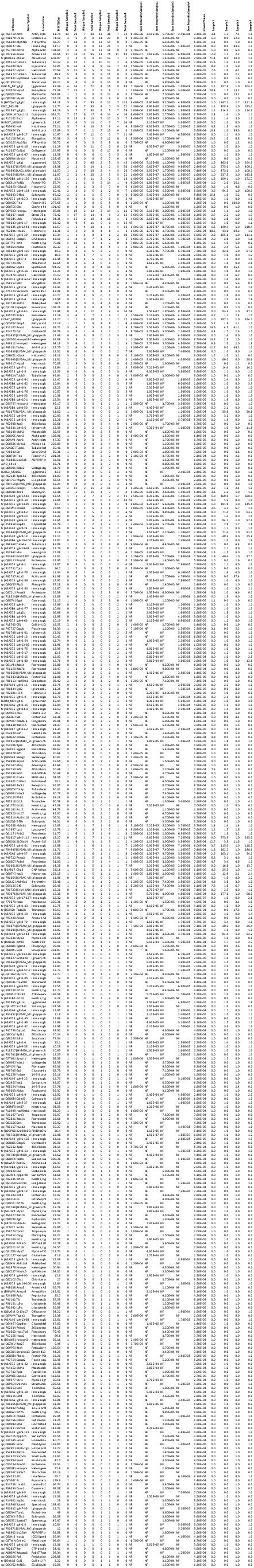

## Notes

### Competing Interest Statement

The authors have declared no competing interest.

